# The VelB intrinsically disordered domain promotes selective heterodimer formation of velvet domain regulatory proteins for fungal development

**DOI:** 10.1101/2025.02.27.640524

**Authors:** Anna M. Köhler, Sabine Thieme, Jennifer Gerke, Karl G. Thieme, Rebekka Harting, Kerstin Schmitt, Oliver Valerius, Wanping Chen, Annalena Höfer, Emmanouil Bastakis, Anja Strohdiek, Antje K. Heinrich, Helge B. Bode, Gerhard H. Braus

**Author notes:** equally contributed (Anna M. Köhler, Sabine Thieme).

## Abstract

Fungi possess several transcription factors with a characteristic velvet domain for DNA-binding and homo- or heterodimerization, which is structurally similar to the mammalian NF-ᴋB Rel homology domain. Velvet dimers control fungal development, virulence and mycotoxin formation. VelB is the only regulator, which carries an intrinsically disordered domain (IDD) within the velvet domain. The IDD as well as the positioning within VelB is conserved in the fungal kingdom. Intrinsically disordered regions contribute to transcription activation and DNA binding and frequently appear in eukaryotic transcription factors. The VelB IDD provides selective heterodimerization as well as protein stability control. The IDD is not required for the formation of the VelB-VeA heterodimer of *Aspergillus nidulans* or *Verticillium dahliae*, but promotes the formation of the VelB-VosA heterodimer. The IDD destabilizes VelB single molecules and also balances its distribution and ratio between both velvet heterodimers. These balances contribute to control appropriate mycotoxin production and sexual development. Herewith, the VelB IDD represents a novel control mechanism of velvet protein stability and heterodimer formation for precise priming of fungal development.

## Introduction

Intrinsically disordered proteins or domains possess an inherent flexibility with several possible conformations in solution instead of a well-defined 3D-structure [1]. This correlates with a significantly increased frequency of amino acid residues for higher net charge and lower mean hydrophobicity in comparison to ordered proteins [2]. Eukaryotic DNA-binding proteins are significantly enriched in disordered domains [3]. Posttranslational modifications of charged or other residues within these domains can alter functions or life time of the transcription factors by transient self-interactions or promiscuous binding to several partner molecules [4]. Intrinsically disorder is present in transcriptional activation domains, which recruit the transcriptional machinery. Disordered domains can provide electrostatic interactions within dynamic complexes by following induced fit mechanisms for binding [4]. A bioinformatic survey of human transcription factors revealed that DNA binding domains with significant order are often flanked by regions with significant disorder [5].

The fungal <u>Vel</u>vet-like <u>B</u> (VelB) transcription factor carries an intrinsically disordered domain within its DNA binding and dimerization domain [6]. VelB is one member of the conserved fungal velvet family of transcription factors which also includes <u>Ve</u>lvet <u>A</u> (VeA), <u>V</u>iability <u>o</u>f <u>s</u>pores <u>A</u> (VosA) and <u>Vel</u>vet-like <u>C</u> (VelC) [7]. Velvet proteins control and coordinate development, virulence and secondary metabolism including the formation of mycotoxins [7–14]. The velvet domain comprises approximately 100 to 200 amino acids and is a protein-protein interaction and DNA-binding domain with structural similarities to the Rel homology domain of the mammalian immune and infection response NF-ᴋB regulator [6,15]. Velvet domain proteins as well as NF-ᴋB regulators bind as homo- or heterodimers to a myriad of genomic sites. The NF-κB heterodimer p65p50 interface provides increased conformational plasticity resulting in significantly stronger affinity to DNA than the corresponding homodimers p50p50 or p65p65 [16].

Fungal VelB acts as light-dependent multifunctional regulator for fungal asexual and sexual development. VelB coordinates differentiation with the appropriate secondary metabolism and furthermore controls spore viability [17,18]. VelB can either form a heterodimer with VeA (VelB-VeA) or with VosA (VelB-VosA) [19]. Aspergillus asexual development is promoted by light and results in the release and dispersal of spores (conidia) into the air [7,20]. VelB activates the *brlA* (*bristle A*) gene encoding the central regulator of the progression of conidiation [21,22]. In contrast, VosA represents a negative regulator of asexual development and represses the *brlA* dependent genetic networks of asexual development, oxidative stress response and the corresponding secondary metabolism [23,24]. Aspergillus develops closed sexual fruiting bodies in darkness and low oxygen pressure representing overwintering structures of this mold in the soil. Specific multinuclear cells (Hülle cells) are formed to nurse the growing fruiting body and to protect it by mycotoxins against fungivores [19,20,25,26]. Nuclear VelB-VeA heterodimer activates sexual development. This transcriptional regulation is linked to epigenetic control by the formation of a trimeric complex of VelB-VeA with the methyltransferase LaeA (loss of the <u>a</u>flatoxin regulator <u>e</u>xpression <u>A</u>) as global regulator of secondary metabolite formation. LaeA is required to synthesize the aflatoxin family mycotoxin sterigmatocystin for protection of the sexual fruiting bodies [18,19,27,28]. VelB-VosA is important for trehalose biogenesis to support spore viability and germination [19,23].

The different VelB interactions are required in response to diverse environmental signals to support distinct developmental programs and the appropriate secondary metabolism [9,18]. The molecular mechanisms of how the fungal cell distributes VelB to the two alternative heterodimers to regulate different sets of target genes are yet unknown. VelB-VeA supports nuclear entry of VelB [18]. This suggests that nuclear VelB-VosA formation might happen by an exchange of the VelB binding partner after nuclear entry. This exchange of the VelB binding partner presumably depends on external signals and the cell-type specific status of *A. nidulans* development. Specific control mechanisms for differential gene expression as well as protein stability regulation, which change the protein homeostasis of velvet domain proteins, are essential for heterodimer formation control [20,29]. Internal protein signal sequences within velvet domain proteins for the promotion of selective heterodimer formation are yet unknown. We compared the VelB intrinsically disordered domain within the fungal kingdom and characterized and analyzed its molecular role in fungal cells or during development to gain further insights into the evolution and function of velvet domain regulatory proteins. The insertion in the VelB velvet domain is evolutionary conserved in filamentous fungi of different divisions. The intrinsically <u>d</u>isordered <u>d</u>omain (hereafter IDD) has a significant impact on protein stability and an even more remarkable potential to select and control VelB protein interactions. The IDD is required to control cellular ratios of different VelB heterodimers and therefore links fungal development to the required secondary metabolite production.

## Results

### VelB is the only member of the velvet family carrying an intrinsically disordered region within the DNA binding and dimerization domain

The crystal structures of the *A. nidulans* VosA and VelB velvet domains revealed a similar fold to the Rel homology domain of the mammalian transcription factor NF-ᴋB [6]. The amino acid sequence similarities between Rel homology and velvet domains comprise only approximately 14%, but important DNA contact sites are conserved. The VelB-VosA_1-190_ heterodimer crystal structure lacks a 99 amino acid (aa) insertion within the VelB velvet domain, which has been removed due to protease treatment during crystallization [6]. Sequence analysis has predicted that this 99 aa sequence is unstructured (Fig S1) and in the following is denominated as intrinsically <u>d</u>isordered <u>d</u>omain (IDD; Fig 1A). VelB is the only member of the *A. nidulans* velvet family with an IDD, which is absent in VeA, VelC or VosA [9].

**Fig 1:**
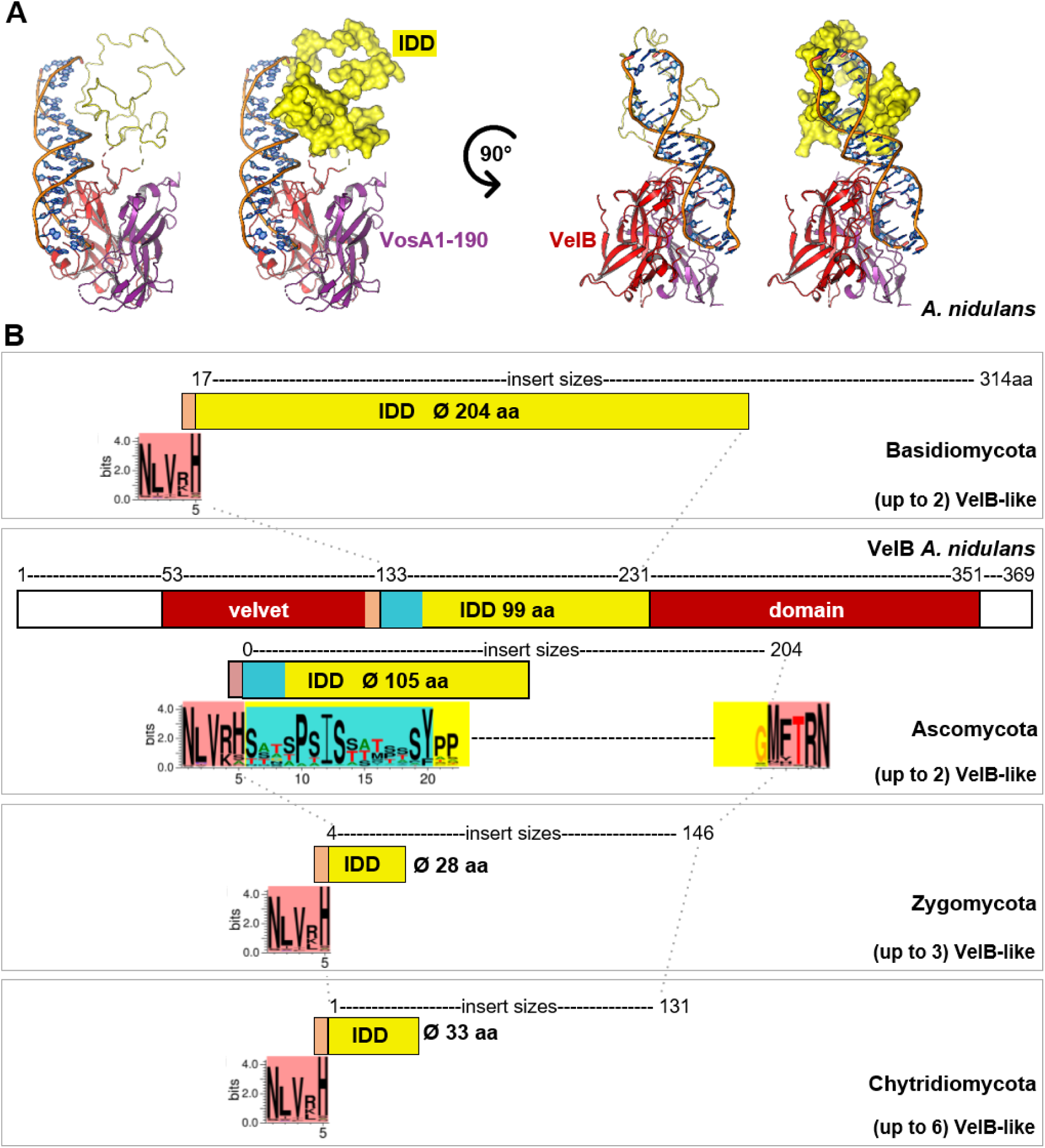
Genes encoding the VelB velvet transcription factors within the fungal kingdom share an intrinsically disordered domain, which is inserted at the same position within their DNA-binding and dimerization domain. **(A)** The crystal structure-derived model of the *A. nidulans* VelB (red)-VosA_1-190_ (purple) heterodimer bound to DNA (according to [6]; PDB4N6R) lacking the VelB intrinsically <u>d</u>isordered <u>d</u>omain (IDD, 99 amino acids; Fig S1). The unstructured PhoA1-150 NMR conformer 6 (PDB 2MLY) has a similar size as the structurally unknown VelB IDD and was used as template in a model to illustrate the size of the VelB IDD (yellow). **(B)** The VelB velvet domain N-terminal boundary of the IDD is highly conserved in the fungal kingdom and its C-terminal residue is in most cases a histidine. The *A. nidulans* VelB IDD of 99 amino acids includes a 15 amino acid motif (turquoise) and a 3’ boundary, which is conserved among Ascomycota. Averages of the amino acid sequence sizes of deduced VelB inserts differ among fungal divisions as indicated in the yellow boxes. The deduced IDD information and the corresponding genes are listed in Table S1 – source data 1 for fungal species with single genes for VelB or in Table S2 for species with at least two isogenes.

A comparison of the deduced VelB amino acid sequences of different fungal genomes provided by the JGI fungal genome database [30] revealed that the insertion of an IDD at a similar position into the velvet domain as in *A. nidulans* is conserved in numerous fungal VelB counterparts (Fig 1B). Deduced VelB IDD sequences of different species from the fungal kingdom (Ascomycota, Basidiomycota, Zygomycota, Chytridiomycota) share the presence of numerous serine residues, but differ considerably (Fig S2). There is a remarkable conservation of five aa residues at the N-terminal IDD boundary inserted into the VelB velvet domain in all four divisions. This boundary is mostly terminated with a positively charged amino acid residue as arginine or lysine, followed by a histidine residue (red box in Fig 1B & S2). Ascomycota share adjacent to the conserved 5’IDD boundary, a serine-rich conserved 15 aa motif (Motif_IDD_) at the N-terminus of the IDD (turquoise box), which is absent in the IDDs of the VelB counterparts of Basidiomycota, Zygomycota or Chytridiomycota (Fig 1 and Fig S2). The VelB velvet domain C-terminal boundary of the IDD is only conserved among ascomycetes (red box in Fig 1B) and includes a highly conserved central threonine residue, which is also found in the 3’ boundary region of various basidiomycetes (Fig S2).

The different fungal groups show strong variations in the size of the analyzed VelB IDDs. Ascomycota and Basidiomycota have larger average sizes of 105 and 204 aa, respectively compared to the IDDs of Zygomycota or Chytridiomycota which are significantly smaller (28 and 33 aa in average, Fig 1 and Fig S3). Ascomycota or Basidiomycota normally possess one *velB* gene. Zygomycota or Chytridiomycota species often acquired two or more *velB*-like gene copies. Especially among the Zygomycota one isoform can have either no (or only a very short) IDD, whereas the other isoform carries a longer IDD (Table S1 and S2).

Taken together, the fungal VelB orthologs share a region of intrinsically disorder at the same position within the velvet domain, and which differs in size and sequence with increasing evolutionary distance. These different IDD sequences and sizes combined with up to six *velB*-like genes present in fungal groups as the Chytridiomycota suggest a rapid *velB* gene evolution, including gene duplications and subsequent IDD variations. A common molecular IDD function is yet elusive, but these fungal VelB insertions presumably would have not been conserved during evolution if they had only occurred randomly or unspecifically.

### The intrinsically disordered domain destabilizes *A. nidulans* VelB

Control of the relative amounts of transcription factors is essential to ensure appropriate heterodimer formation for the regulation of target genes in response to changing environments. VelB full length protein stability was compared to a variant lacking the IDD in fungal strains expressing the proteins C-terminally fused to GFP (VelB*-*GFP, VelB^ΔIDD^-GFP) under the control of the endogenous *velB* promoter. The fungal cultures were supplemented with cycloheximide after 24h of vegetative growth to inhibit protein biosynthesis. Protein crude extracts were analyzed by Western experiments hourly from 0 to 5h after supplementation. Cycloheximide assays revealed that full length VelB-GFP is less stable than VelB^ΔIDD^-GFP during filamentous growth. Whereas approximately 90% of VelB^ΔIDD^-GFP was still present after five hours, the relative protein amounts of VelB-GFP with the IDD was reduced to approximately 56% (Fig 2A). This suggests that the IDD provides a destabilizing function, which allows the fungal cell to control VelB protein turnover under specific environmental conditions.

**Fig 2:**
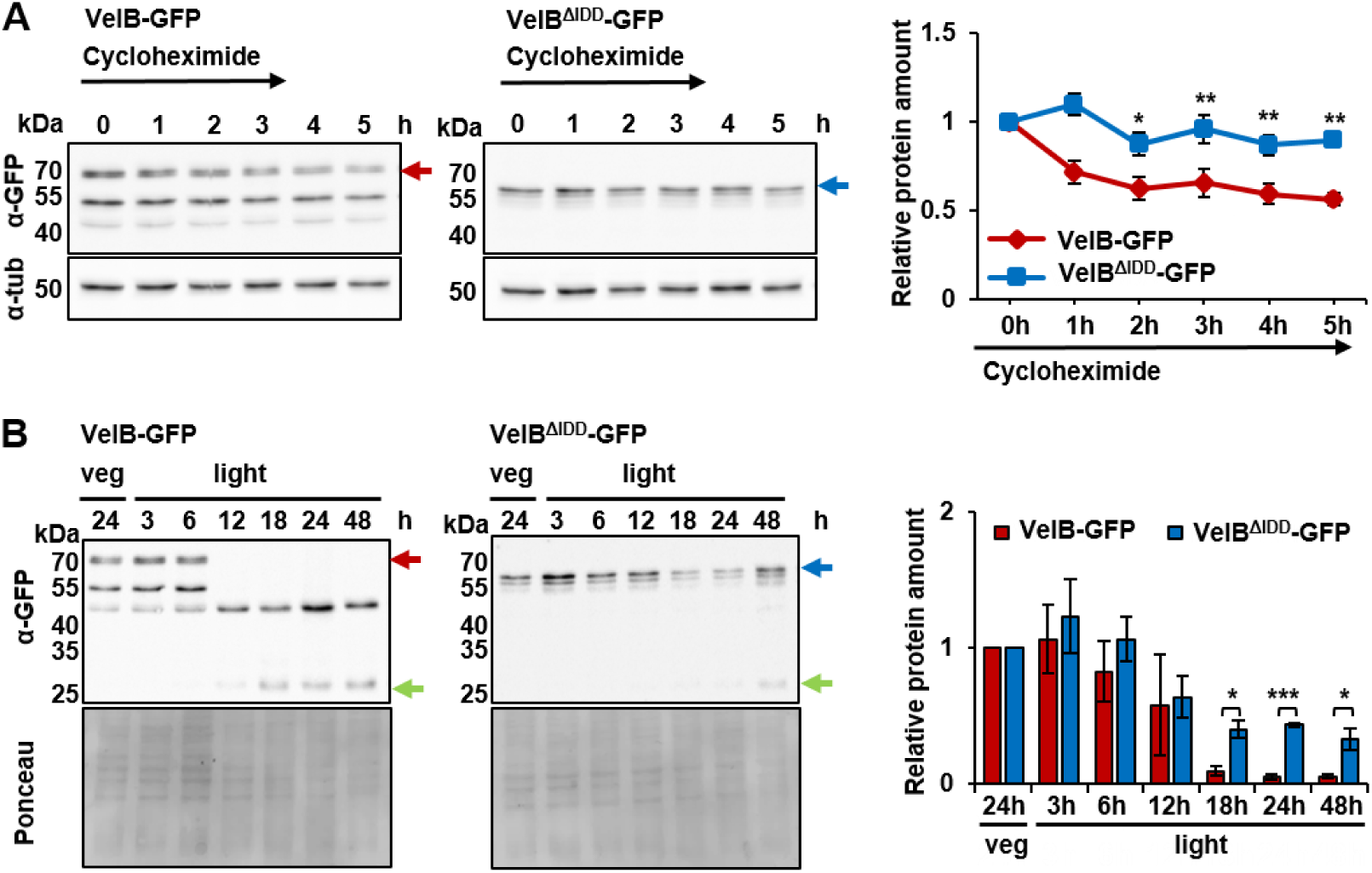
The IDD destabilizes *A. nidulans* VelB. **(A)** Cycloheximide chase analysis of VelB-GFP and VelB^ΔIDD^-GFP protein degradation. Cycloheximide was added after 24 hours (h) of vegetative fungal growth at 37°C to submerged cultures. Samples were taken from zero to five hours post supplementation. Western experiments applying the α-GFP antibody to crude extracts show less stable VelB-GFP compared to VelB^ΔIDD^-GFP. α-Tubulin was used as loading control. Lack of the IDD increases protein stability. Error bars indicate standard error of the mean (SEM) of four biological replicates normalized against the tubulin signal. *p*-value was calculated with standard deviation. * *p* < 0.05, ** *p* < 0.01. **(B)** Relative abundance of VelB-GFP or VelB^ΔIDD^-GFP during fungal development. Vegetative grown mycelia were shifted after 24h to solid minimal medium and cultivated for indicated time periods in light at 37°C for induction of asexual development. Western analysis shows that VelB-GFP (red arrow) is degraded during early asexual development when degradation products and free GFP (green arrow) become visible. VelB^ΔIDD^-GFP (blue arrow) is stable and still detectable after 48h cultivation in light. The diagram depicts the quantification of VelB-GFP and VelB^ΔIDD^-GFP relative to the protein amount of vegetative grown cultures normalized against Ponceau. Error bars indicate the SEM of three biological replicates. *p*-value was calculated with standard deviation. * *p* < 0.05; *** *p* < 0.005.

Relative VelB protein abundances in protein crude extracts of both versions were compared between vegetative growth and upon induction of asexual development when the VelB-VosA heterodimer is active. The relative normalized expression of *velB^ΔIDD^* compared to *velB* was similar from six to 18h incubation under asexual conditions (Fig S4). Vegetative cultures incubated for 24h comprise the 67 kDa VelB-GFP protein (Fig 2B, red arrow) or the 56 kDa VelB^ΔIDD^-GFP protein (blue arrow), respectively. Full length VelB-GFP was only detectable during early asexual development but is mostly degraded after 18h of differentiation when only free GFP is still present (∼ 27 kDa, Fig 2B, green arrow). This indicates a rapid VelB degradation during ongoing asexual development. In contrast, VelB^ΔIDD^-GFP fusion protein degradation is slower, with approximately 30% of the relative protein abundance present after 48h incubation under asexual development inducing conditions (Fig 2B).

This corroborates that in *A. nidulans* the presence of the IDD in the wildtype VelB protein contributes to protein degradation, especially during light-induced asexual development. In contrast, truncated VelB without IDD is more stable under the same cultivation conditions, which supports the destabilizing impact of IDD on the VelB protein. In conclusion, this data suggest a VelB IDD-dependent degradation control as mechanism to restrict cellular VelB abundance for its adjusted channeling to complex formation with either VeA or VosA during fungal development.

### The VelB intrinsically disordered domain enables selective heterodimer formation with VosA, which ensures nuclear localization of VelB

Independently of light, VelB is localized in the cytoplasm prior to its nuclear import as well as in the nucleus for DNA binding [18]. Fluorescence microcopy was applied to examine whether the IDD affects VelB nuclear localization. Strains with full length VelB fused to GFP (VelB*-*GFP) were compared to ones with the truncated version without IDD (VelB^ΔIDD^-GFP). A constitutively expressing GFP (OE GFP) strain and the wildtype strain served as controls to exclude unspecific GFP background signals (Fig S5A). Independent from the presence of the IDD, VelB accumulated in nuclei of vegetative growing hyphae, however, an additional subpopulation appeared in the cytoplasm (Fig 3A).

**Fig 3:**
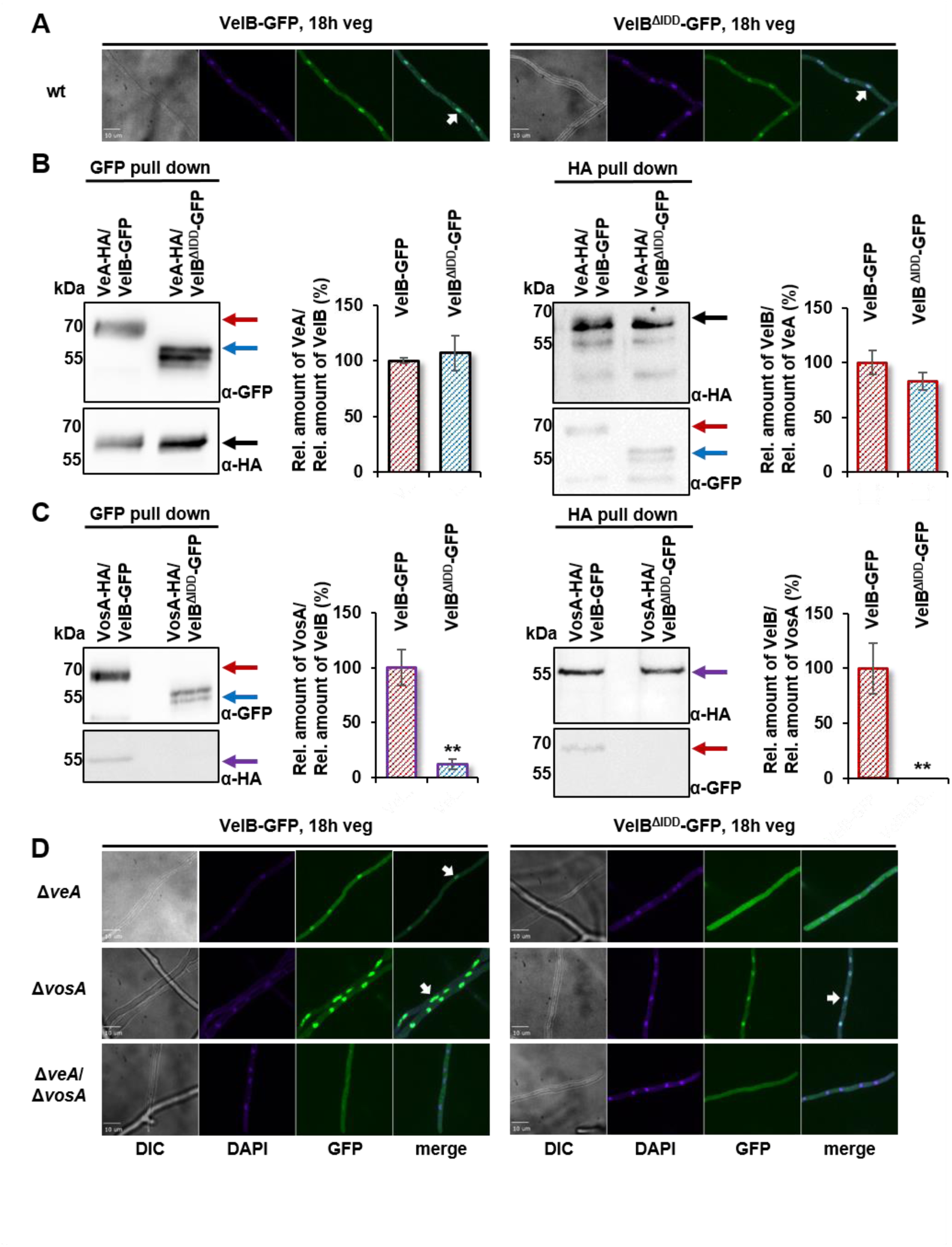
*A. nidulans* VelB IDD is required for VelB-VosA heterodimer formation but not for VelB-VeA dimer formation or VelB nuclear localization. **(A)** Fluorescence microscopy of VelB-GFP with or without IDD revealed nuclear accumulation (white arrows). *A. nidulans* strains were grown vegetatively in submerged cultures for 18 h at 30°C. Nuclei were visualized with DAPI (4′,6-<u>dia</u>midin-2-<u>p</u>henylindol). wt = wildtype, DIC = differential interference contrast, scale bar = 10 µm. **(B)** Predominant nuclear localization of VelB-GFP is similar in wt, *veA* (Δ*veA*) and *vosA* (Δ*vosA*) single deletion strains, but changes to cytoplasmic localization in the veA/vosA double deletion strain (Δ*veA*/ Δ*vosA).* VelB^ΔIDD^-GFP is localized in the nucleus in wt and *vosA* deletion strains but found in the cytoplasm in *veA* or *vosA*/*veA* deletion strains. **(C)** Co-immunoprecipitation experiments included GFP and HA pull downs of strains expressing either VeA-HA, VelB-GFP or VelB^ΔIDD^-GFP fusion proteins followed by western experiment detection with GFP and HA antibodies. The relative amount of VeA against VelB (black diagram) or the relative amount of VelB against VeA (red diagram) was quantified and revealed that VelB and VeA form a heterodimeric velvet complex with and without the IDD. **(D)** Co-immunoprecipitation experiments of strains expressing VosA-HA and VelB-GFP or VelB^ΔIDD^-GFP fusion proteins were followed by western experiment detection with GFP and HA antibodies. The relative amount of VosA against VelB (violet diagram) or the relative amount of VelB against VosA (red diagram) was quantified revealing that VelB without IDD did not form a VelB-VosA complex. Error bars indicate standard error of the mean (SEM) of three biological replicates. *p-*value was calculated with standard deviation. ** *p* < 0.01. Black arrows = VeA-HA, red arrows = VelB-GFP, blue arrows = VelB^ΔIDD^-GFP, violet arrows = VosA-HA.

Intrinsically disordered proteins can function as hubs in protein interaction networks [31–33]. VeA enhances the transport of VelB through the heterodimer VelB-VeA from the cytoplasm into the nucleus in the dark [18], whereas the VelB-VosA heterodimer resides predominantly in the nucleus [19].

GFP pull down experiments were conducted to investigate whether VelB without the IDD still interacts with VeA or VosA within the vegetative fungal cell. Mycelium from vegetative cultures was used for GFP pull downs followed by LCMS analysis for the identification of interacting proteins. The bait proteins VelB-GFP and VelB^ΔIDD^-GFP were detected with similar LFQ intensities and MS/MS counts in biological replicates (Table 1). Comparable numbers of unique peptides were identified, indicating that the pull downs worked equally well for both VelB variants. Proteins were filtered for detection in at least two out of three biological replicates with MS/MS counts ≥ 4, unique peptides ≥ 3 and logarithmised LFQ intensity ≥ 20 and were absent in the control strain (Table 1, Table S3).

**Table 1:** Velvet domain proteins identified from GFP pull downs of VelB-GFP and VelB^ΔIDD^-GFP.

| Sys. name | Std. name | LFQ intensity |  | MS/MS counts |  | Unique peptides |  |
| --- | --- | --- | --- | --- | --- | --- | --- |
|  |  | *velB:gfp* | *velB<sup>ΔIDD</sup>:gfp* | *velB:gfp* | *velB<sup>ΔIDD</sup>:gfp* | *velB:gfp* | *velB<sup>ΔIDD</sup>:gfp* |
| AN0363 | VelB | 28.88 | 29.77 | 735 | 728 | 15 | 16 |
|  |  | 26.18 | 27.77 | 191 | 139 | 11 | 9 |
|  |  | 25.22 | 28.05 | 112 | 36 | 13 | 9 |
| AN1052 | VeA | 25.51 | 26.35 | 406 | 405 | 17 | 16 |
|  |  | 23.75 | 25.85 | 125 | 98 | 13 | 10 |
|  |  | 23.78 | 26.01 | 89 | 47 | 16 | 12 |
| AN1959 | VosA | 24.32 | 14.92 | 166 | 1 | 11 | 1 |
|  |  | 22.03 | 15.86 | 8 | 0 | 3 | 0 |
|  |  | 16.89 | 15.24 | 1 | 0 | 1 | 0 |
| AN0807 | LaeA | 21.35 | 20.68 | 47 | 30 | 7 | 6 |
|  |  | 17.39 | 16.38 | 0 | 0 | 0 | 0 |
|  |  | 19.74 | 17.22 | 2 | 0 | 2 | 0 |

VelB-GFP with and without the IDD reliably pulled the VeA protein. VosA was only identified in experiments with the full length VelB-GFP but not with VelB<sup>ΔIDD</sup>-GFP. In contrast, the IDD is dispensable for VelB-VeA-LaeA velvet complex formation. Sys. Name = systematic name, Std. name = standard name.

The GFP pull downs revealed 32 interaction candidates exclusively found for VelB^ΔIDD^-GFP, which were not detected with full length VelB-GFP as bait (Fig S6). These proteins were sorted and grouped according to their known or predicted cellular function or localization: mRNA translation, primary metabolism, RNA maturation and processing, membrane/cell wall, signaling, cell compartments, DNA binding and unknown function (Table S3, Fig S6A). For almost half of the putative interaction partners (14 proteins) a nuclear localization signal with a score ≤ 5 was predicted by employing the cNLS Mapper program (Kosugi et al., 2009; Table S3 underlined AN numbers). Proteins with this score presumably can shuttle between, and can be localized in both, the cytoplasm and the nucleus. The velvet domain protein VeA was pulled by both VelB-GFP protein variants with similar efficiency in all experiments. Similarly, the catalase CatB was also always identified as VelB interacting protein supporting a link to the fungal oxidative stress response (Table 1 and Table S3). Only one protein was exclusively pulled with VelB-GFP: the velvet domain protein VosA.

Co-immunoprecipitation (Co-IP) experiments from vegetative mycelium of strains expressing *veA:HA* or *vosA:HA* and *velB:gfp* or *velB^ΔIDD^:gfp* under their native promoters were conducted to verify the velvet heterodimer formation of VelB with VeA and VosA. Western experiments revealed that VeA-HA signals with equal intensity in both GFP pull downs (Fig 3B, black arrow) and the VeA-HA pull down resulted in comparable abundance of VelB-GFP (HA pull down, red arrow) with or without IDD (HA pulldown, blue arrow). These experiments confirmed that VelB-VeA interaction is independent of the VelB IDD. The Co-IPs with VelB variants and VosA show VelB-GFP and VelB^ΔIDD^-GFP with similar abundance in the GFP pulldown experiments (Fig 3C red/blue arrows). In contrast, VosA-HA was only detected for the full length VelB-GFP pull down (Fig 3C, GFP pull down, violet arrow). Quantification of western blot signal intensities revealed a ten-fold increased abundance of VosA in the VelB-GFP strain compared to the VelB^ΔIDD^-GFP strain. The reciprocal experiment resulted in the identification of VosA-HA in equal amounts for both HA pull downs (Fig 3C, HA pull down, violet arrow). VosA-HA co-enriched VelB-GFP (HA pull down, red arrow) but never VelB^ΔIDD^- GFP. These findings corroborate that the IDD prevents multiple additional interactions, which were monitored in the VelB^ΔIDD^-GFP pulldown. The IDD specifically promotes the formation of the VelB-VosA heterodimer in *A. nidulans in vivo* during vegetative growth conditions.

Equivalent pull downs were conducted with VelB orthologous proteins Vel2-GFP and Vel2^ΔIDD^-GFP of the plant pathogenic ascomycete *V. dahliae* to investigate whether the involvement of the IDD for the interaction with VosA is conserved in different fungi. The interactome of *V. dahliae* Vel2^ΔIDD^-GFP showed a decrease in potential interaction partners compared to *A. nidulans* Vel2^ΔIDD^-GFP (Fig S6B, C and Table S4). *V. dahliae* Vel1 (VeA) can interact with Vel2 with or without IDD. In contrast and as found in *A. nidulans*, Vos1 as counterpart of VosA can only significantly interact with the Vel2 full length protein with the IDD (Fig S6B, C). These experiments revealed and support an evolutionary conserved IDD function in heterodimer partner selection.

Microscopy experiments were carried out to investigate whether the IDD affects VelB localization in dependence of VeA or VosA. VelB-GFP and VelB^ΔIDD^-GFP localization was analyzed in *veA* and *vosA* single and double deletion strains after 18h of vegetative growth. VelB-GFP is localized in the nuclei whereas VelB^ΔIDD^-GFP is dispensed throughout the hyphae without nuclear accumulation in absence of *veA* (Fig 3D). Nuclear VelB accumulation is not altered when *vosA* is missing but both VelB variants are absent from nuclei in strains lacking *veA* and *vosA* (Fig 3D). Protein abundance was analyzed by western experiments and shows similar protein levels for all strains investigated (Fig S5B). The same localizations were observed after 18h sexual and asexual development (Fig S5C, D). These results indicate that external factors like light or darkness do not influence VelB localization in the early developmental state. However, all these data support that VelB needs the interaction of either VeA or of VosA for nuclear localization in all tested conditions.

### VeA counteracts VelB-VosA heterodimer formation and suggests VeA as preferred VelB interaction partner

VelB-VosA heterodimer formation according to the previously determined crystal structure was aimed for by recombinant expression of the full length VelB protein with a truncated version of VosA encompassing residues 1-190 (VosA_1-190_) in *E. coli* [6]. Therefore, it was investigated whether a full length VosA requires the VelB IDD for heterodimer formation *in vitro.* Full length fusion proteins of VosA-GST and VelB-His or VelB^ΔIDD^-His were recombinantly expressed in *E. coli* followed by GST-pull downs where the lysate of VosA-GST was either mixed with purified VelB-His or VelB^ΔIDD^-His. VosA-GST (Fig S7, violet arrow) co-enriched with VelB-His (red arrow) and VelB^ΔIDD^-His (blue arrow). This result demonstrates that the VelB IDD itself is not directly required for the interaction of VelB with VosA *in vitro.* This supports indirect effects of other cellular components that impact the IDD to prevent the VelB-VosA interaction *in vivo*.

Fungal Δ*velB*/Δ*veA* and *velB^ΔIDD^*/Δ*veA* strains were generated to analyze whether the VelB-VosA heterodimer formation is affected by VeA and the IDD *in vivo*. Samples of these strains were analyzed from vegetative mycelium as well as from light-induced asexually grown mycelium because the VelB-VosA heterodimer is a regulator of this developmental program [23]. Western experiments show similar protein intensities of VelB-GFP and VelB^ΔIDD^-GFP in the *veA* deletion strain from vegetative and asexual mycelium (Fig S8A). GFP pull downs with VelB-GFP and VelB^ΔIDD^-GFP in *veA* deletion background were performed. In absence of the *veA* gene, VosA is able to bind to VelB independently of the IDD (Fig S8B, Table S5). CoIP experiments with VelB-GFP or VelB^ΔIDD^-GFP and VosA-HA in strains deleted in *veA* show a similar result. VosA-HA was able to recruit VelB^ΔIDD^-GFP and *vice versa* in Δ*veA* (Fig S9).

These data suggest that heterodimerization of VelB^ΔIDD^ with VosA is possible in absence of *veA in vivo* and accordingly is also possible *in vitro.* The presence of an intact *veA* gene favors heterodimerization with VeA as preferred interaction partner of VelB *in vivo*.

### The VelB IDD promotes asexual development in *A. nidulans*

The VelB-VeA and VelB-VosA heterodimers coordinate fungal development [18]. VelB-VeA is required for the sexual and VelB-VosA for the asexual pathway, respectively [18]. The impact of the VelB IDD on *A. nidulans* developmental programs was examined by comparing phenotypes of wildtype, *velB* deletion (Δ*velB*) or *velB^ΔIDD^* mutant strains.

Light/illumination promotes asexual development and results in conidiophores with green asexual spores on agar plates for the wildtype as well as the Δ*velB* strain (Fig 4A). Point inoculations of the strains lead to similar colony morphology of the *velB^ΔIDD^* and the wild type strain when incubated in asexual development-inducing conditions (Fig S10A). In contrast, spreading spores area-wide on culture plates resulted in a different phenotype. The strain lacking the VelB IDD forms increased amounts of aerial hyphae as precursors of conidiophores resulting in a white fluffy appearance (Fig 4A, red arrow). Conidiophores with green conidiospores are rare and only formed at the edge of the agar plate (Fig 4A, blue arrow). Quantification of conidiospores after growth with light revealed significantly reduced amounts (12% after three days, 14% for the *vel2^ΔIDD^* strain after seven days) relative to wildtype (Fig 4B). This is similar to the *vel2* deletion strain, which also shows a decrease in conidiospore production. This phenotype was complemented by *in locus* reintroduction of the functional *A. nidulans velB:gfp* fusion construct (*velB^AnIDD^:gfp*) or with orthologous sequences from the ascomycetes *A. fumigatus* or *V. dahliae* (*velB^AfIDD^, velB^VdIDD^*). This corroborates that IDD exchanges with different fungi result in functional proteins and can support *A. nidulans* VelB in providing appropriate asexual development and spore formation.

**Fig 4:**
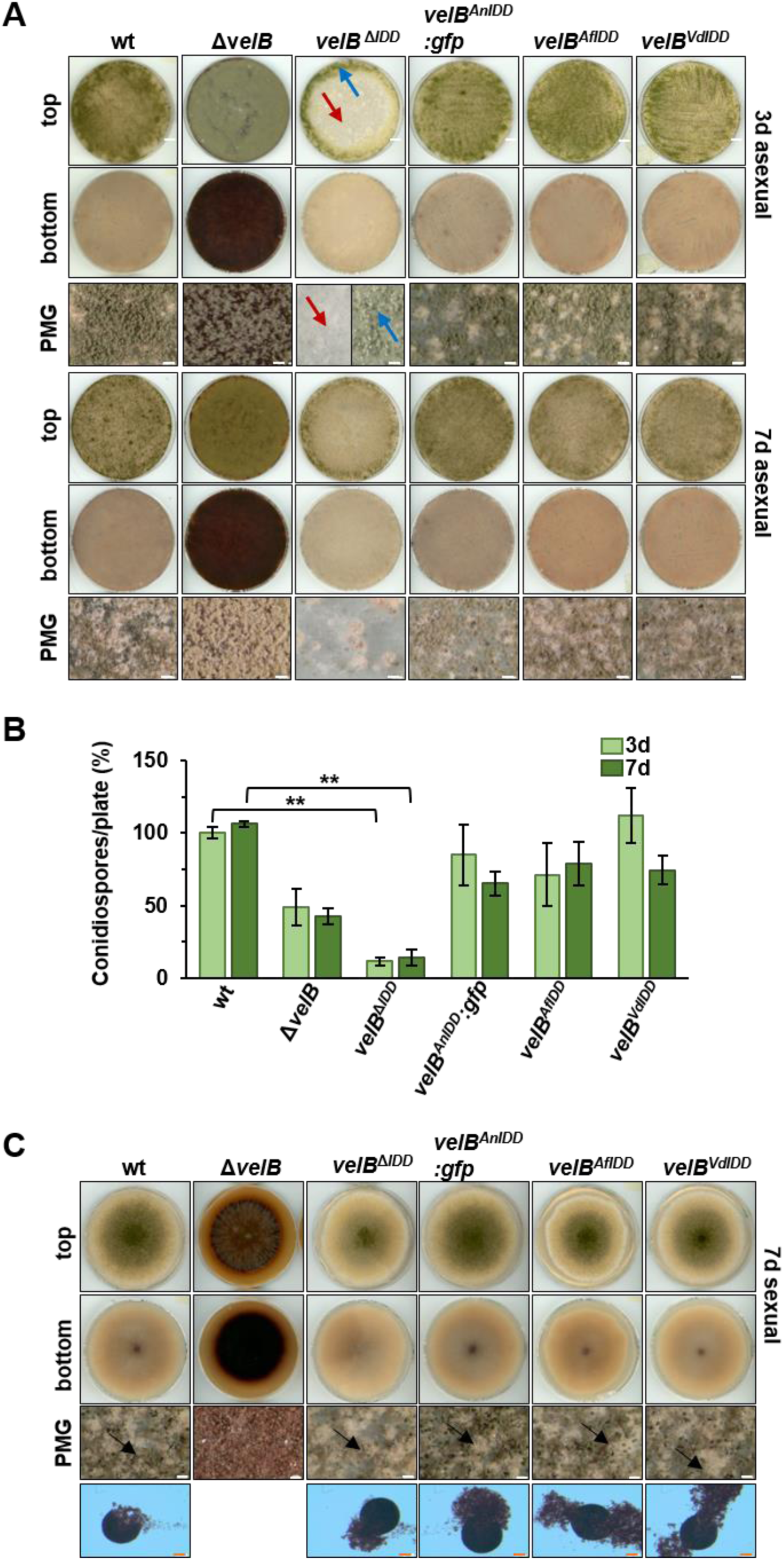
The VelB IDD is required for efficient asexual spore formation in *A. nidulans*. **(A)** Phenotypes of wildtype (wt), *velB* deletion (Δ*velB*)*, velB^IDD^* deletion (*velB**^Δ^**^IDD^*) and complementation strains with the IDD of *A. nidulans* (*velB^AnIDD^:gfp*)*, A. fumigatus* (*velB^AfIDD^*) or *Verticillium dahliae* (*velB^VdIDD^*) on solid minimal medium. 10^5^ spores were distributed over the plate and grown in light for three or seven days (d) at 37°C. Red arrows indicate aerial hyphae and blue arrows conidiophores. Scale bar = 100 µm. **(B)** Quantification of conidiospores from strains shown in (A). Error bars indicate standard error of the mean (SEM) of three biological replicates. *p*-value was calculated with standard deviation. *** *p* < 0.005. **(C)** Phenotypes of wildtype, *velB* deletion*, velB^IDD^* deletion and complementation with the IDD of *A. nidulans, A. fumigatus* or *V. dahliae* into *velB^ΔIDD^* strains. Strains were point inoculated with 10^5^ spores on solid minimal medium and incubated in dark for seven days (d) at 37°C. Black arrows indicate mature cleistothecia. PMG = photomicrograph, scale bar (white) = 100 µm, (red) = 50 µm.

Dehydroaustinol, product of the *aus* gene cluster-encoded proteins, is one of two compounds which signal the induction of sporulation of *A. nidulans* [9,35]. The impact of VelB on the *aus* cluster genes was examined in more detail. The *aus* cluster comprises fourteen genes including the four genes *ausI, ausJ, ausM* and aus*N,* which are essential for austinol and dehydroaustinol biosynthesis (Lo *et al.*, 2012). They were chosen to test their expression via qRT-PCR in the Δ*velB* and *velB^ΔIDD^* strains. Transcript levels of the *ausI*, *ausJ ausM* and *ausN* genes were significantly reduced in the *velB* deletion strain, suggesting that they are controlled by the encoded transcription factor (Fig S10B). Transcription of the *ausJ*, *ausM* and *ausN* is significantly reduced in the *velB*^ΔIDD^ strain, implying an IDD dependent function in regulation. This suggests that the IDD-dependent VelB-VosA complex is required to activate several genes involved in the biosynthesis of austinol and dehydroaustinol which stimulates asexual development. These findings are consistent with the decreased conidiospore amounts in the *velB* deletion and the *velB^ΔIDD^* mutant strains (Fig 4B).

*A. nidulans* reproduces sexually through the formation of cleistothecia harboring sexual ascospores as overwintering structures in the soil. The *velB* deletion mutant is completely inhibited in sexual reproduction in the dark (Fig 4C). In contrast, the *velB^ΔIDD^* mutant strain develops mature cleistothecia filled with ascospores after seven days in the dark. Therefore, the IDD is not required for sexual development and can also be exchanged by orthologous IDD sequences from *A. fumigatus* or *V. dahliae* without impact on the sexual development.

Whereas VelB IDD and, therefore, the heterodimer VelB-VosA are dispensable for sexual development, the VelB-VeA heterodimer, which still can be formed without the VelB IDD, is absolutely necessary for *A. nidulans* asexual conidiation. The IDD reduces VelB-VeA formation and promotes VelB-VosA formation for induction of the asexual developmental program linked to the appropriate corresponding secondary metabolism.

### The IDD-dependent VelB-VosA heterodimer controls sterigmatocystin production, whereas VelB-VeA controls secondary metabolites relevant for sexual development

Velvet domain proteins connect fungal development with secondary metabolism [18]. A *velB* deletion strain secretes dark red-brown pigments into the agar plate, whereas the *velB^ΔIDD^* strain shows a lighter color on the bottom of the plate compared to wildtype (Fig 4A, C, Fig 5). This reflects differences in the regulation of <u>s</u>econdary <u>m</u>etabolism (SM) controlled by the interplay of VelB with its interaction partners. SM analyses were conducted for *velB, velB^ΔIDD^, veA* and *vosA* single, double and triple deletion strains cultivated for three days under asexual or seven days under sexual development-inducing conditions. The *veA* deletion strain has a similar coloring phenotype like strains without *velB* but Δ*vosA* shows no red-brown colored medium and hyphae. SM analysis revealed accumulation of the *orsellinic* (*ors*) cluster product F9775A/B (**2**) in all strains with red-brown color (Fig 5, Fig S11 and S13, Table S6). Loss of *veA* or *vosA* and additional loss of the IDD has no significant impact on the colony color, because double mutants *velB^ΔIDD^*/Δ*veA* and *velB^ΔIDD^*/Δ*vosA* resemble the corresponding *veA* and *vosA* single deletion phenotypes, respectively (Fig 5).

**Fig 5:**
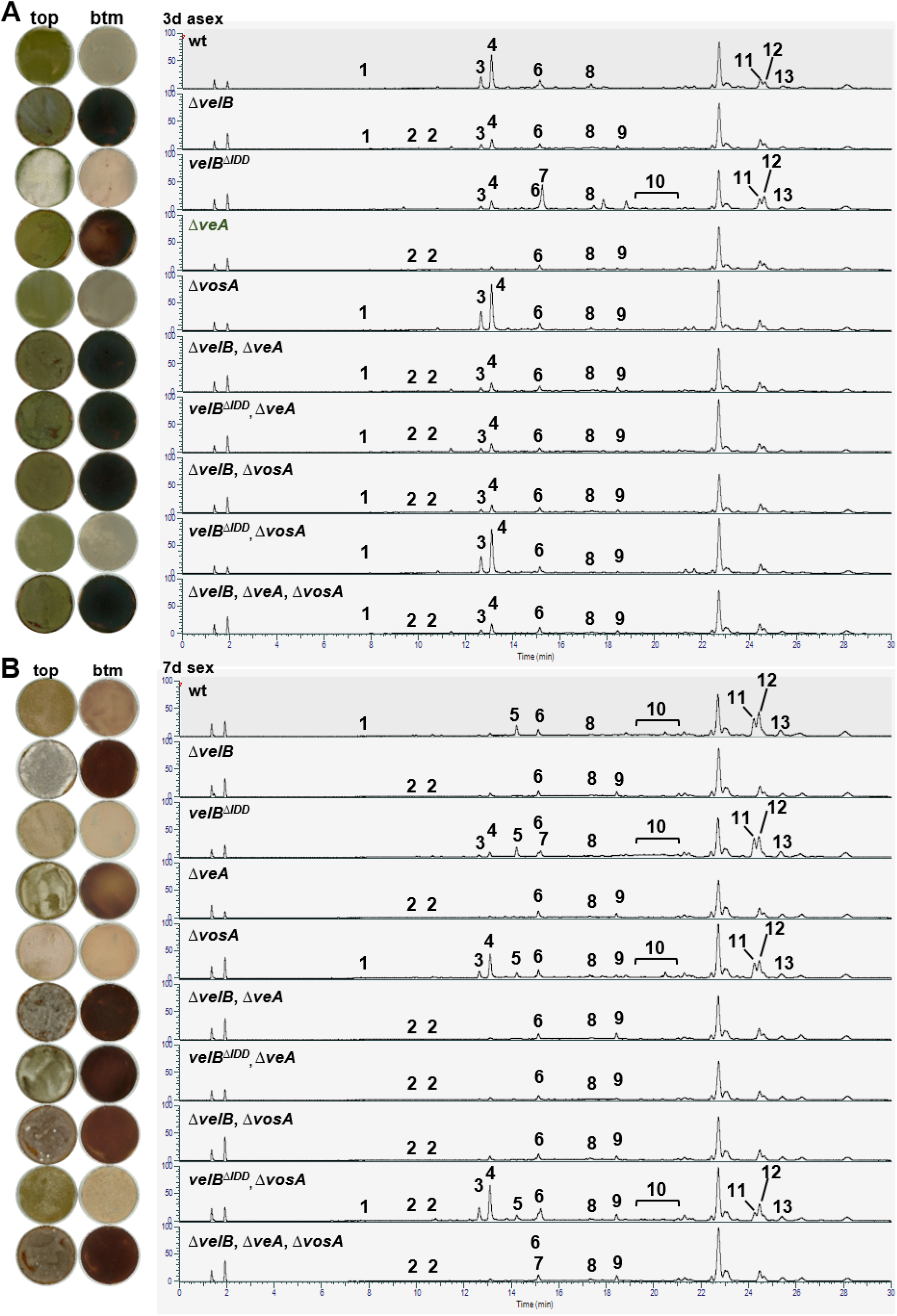
VelB, VeA and VosA regulate secondary metabolite production, including VelB IDD control of sterigmatocystin biosynthesis. Charged aerosol detector (CAD) chromatogram of secondary metabolite extracts obtained from 3 days asexual **(A)** and 7 days sexual **(B)** incubated strains. The numbers highlight peaks with identified metabolites listed in Table S6. Cichorine (**1**), F-9775A/B (**2**), Austinol (**3**), Dehydroaustinol (**4**), Arugosin H (**5**), Emericellamide C (**6**), Sterigmatocystin (**7**), Emericellamide E (**8**), Terrequinone A (**9**), Arugosin A (**10**), Emericellin (**11**), Shamixanthone (**12**), Epishamixanthone (**13**). Structures are shown in Fig S13.

SM extracts of *ΔvelB, velB^ΔIDD^*, Δ*veA* and all double deletions strains including Δ*velB* or Δ*veA* display less austinol (**3**) and dehydroaustinol (**4**) compared to the wildtype or the complementation strain, with the Δ*vosA* strains as the only exceptions (Fig S10A, Fig S11). These data combined with *aus* gene expression experiments underline the importance of VelB for asexual spore formation.

Sterigmatocystin is usually found with low abundances in laboratory wild type/reference strains. The *velB^ΔIDD^* strain produces high amounts of this mycotoxin during asexual development (**7**, Fig 5A, Fig S11A, Fig S12A). LCMS analysis of SM extracts from sexual development revealed additional sterigmatocystin production in the Δ*vosA* or *velB^ΔIDD^*/ΔvosA strain (Fig 5B, Fig S12B). <u>T</u>hin layer <u>c</u>hromatography (TLC) visualized sterigmatocystin (**7**) after derivatization of the compounds on the silica plates with AlCl_3_. A five-fold increased sterigmatocystin abundance was detected at 366 nm in the *velB^ΔIDD^* mutant strain after three and seven days of growth in light compared to wildtype or *velB^AnIDD^:gfp* complementation strains (Fig S11B, C). Therefore, the VelB-VosA heterodimer possibly acts as repressor for sterigmatocystin biosynthesis. The *velB^ΔIDD^*, Δ*vosA* and *velB^ΔIDD^*/ΔvosA strain show wild type- like production of the antraquinones arugosin H (**5**), arugosin A (**10**) as well as the xanthones emericellin (**11**), shamixanthone (**12**) and epishamixanthone (**13**) in sexual developmental samples (Fig 5B). These metabolites were not detected in asexual samples, except in the *velB^ΔIDD^* strain. The VelB-VeA, but not the VelB-VosA complex influences antraquinone and xanthone production of the *monodictophenone* (*mdp*) cluster, which are important metabolites for sexual development [26].

The LCMS analysis additionally showed that chichorine (**1**) was increased in Δ*vosA* and *velB^ΔIDD^*/Δ*vosA*, which is low or absent in all other strains. This suggests that the VelB-VeA heterodimer has either an activating function or the VelB-VosA heterodimer an inhibitory function on the cichorine cluster. Emericellamide production (**6** and **8**) was decreased in all strains compared to wild type under asexual but increased under sexual development inducing conditions. VelB-VosA positively controls terrequinone A (**9**) synthesis, because it was found in *veA* or *vosA* and especially in the *velB* deletion strains but was absent in the *velB^ΔIDD^* strain. This analysis of differences in secondary metabolite formation highlights the importance of precise and accurate control of velvet heterodimer formation to fulfill the distinct molecular functions of the various velvet heterodimer complexes. The IDD allows that VelB can operate specifically depending on its interaction partner as promoting or as inhibiting regulator for the formation of appropriate secondary metabolites that are tightly connected with the distinct fungal differentiation programs.

## Discussion

Intrinsically disordered regions within regulators of gene expression contribute to transcriptional activation or DNA binding. Here we show that the acquisition of an additional intrinsically disordered domain within the DNA binding and dimerization domain of a single member of a gene family of transcriptional regulators, which can form homo- as well as heterodimers allows selective heterodimer formation adjusted to different developmental eukaryotic differentiation programs. A single member of the conserved fungal velvet regulatory gene family allows selective formation of VelB heterodimers with either the velvet domain protein VeA or with VosA. This enables VelB to operate depending on its interaction partner as positive or negative regulator for a specific differentiation program. VelB dependent differentiation is connected with the production of appropriate secondary metabolites to communicate with the environment. The ratio of heterodimers of VelB without the IDD is significantly changed towards increased VelB-VeA and strongly decreased VelB-VosA heterodimers (Fig 6). *In vitro* VosA can form a heterodimer with VelB independently of the IDD. VelB-VosA heterodimer formation without IDD is also possible *in vivo* in a *veA* deletion strain. More VelB proteins are accessible for VosA binding without the competition of VeA. This suggests a higher VelB-VeA binding affinity than VelB-VosA. It can be assumed that VelB IDD switches into a conformation that restricts VosA interaction under specific conditions during fungal life cycle.

**Fig 6:**
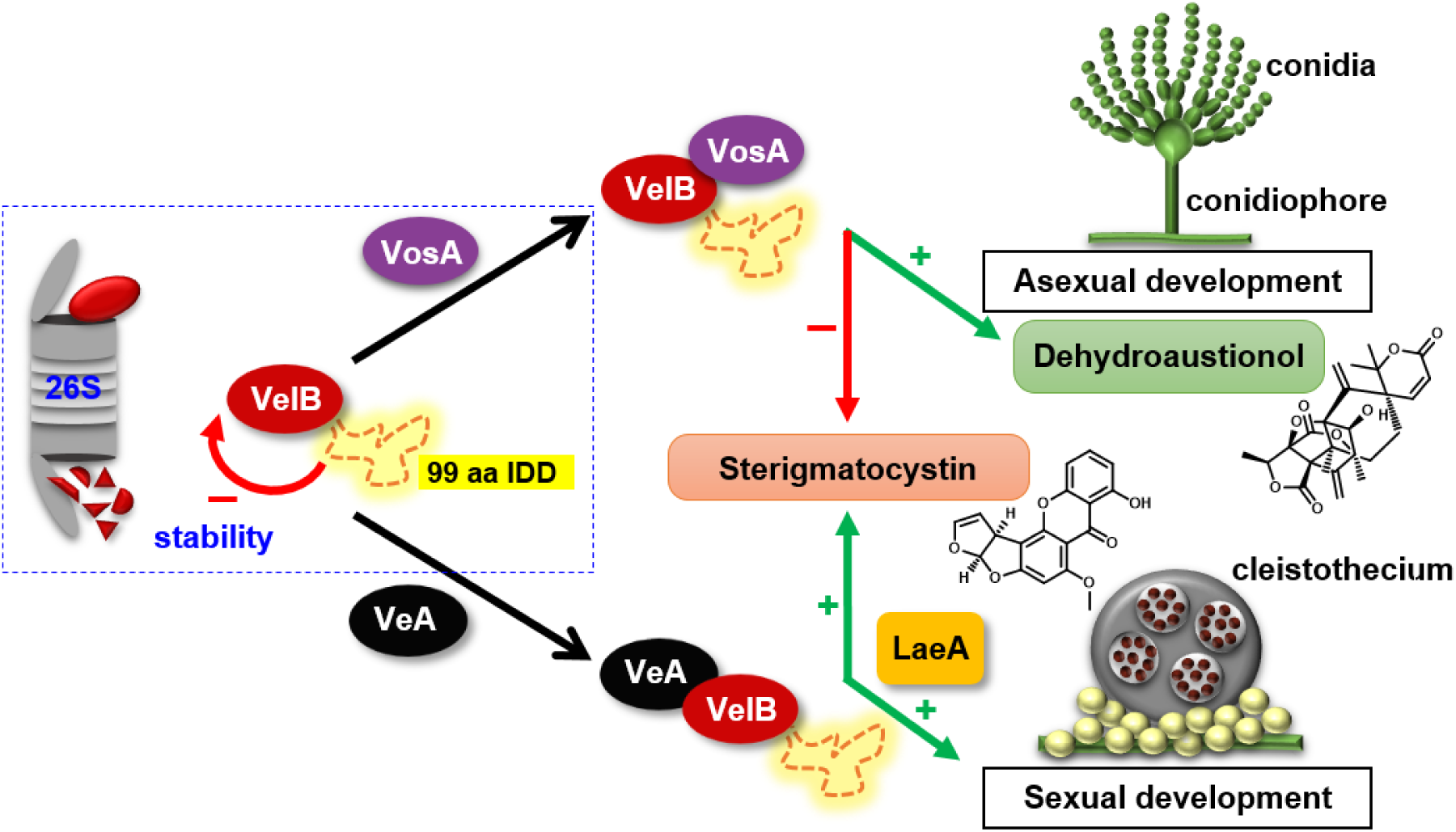
Model of selective VelB IDD heterodimer formation in *A. nidulans*. The VelB dimerization control in *A. nidulans* includes protein stability control and selective heterodimerization. The IDD destabilizes VelB for 26S proteasome degradation. VelB IDD controls selective molecular heterodimer formation. VelB-VosA activates dehydroaustinol production as signal for asexual conidia formation. VelB-VosA reduces sterigmatocystin production either by direct repression or indirectly by reducing VelB levels by competitive VelB-VeA heterodimer formation. Sexual development and sterigmatocystin biosynthesis are induced by VelB-VeA-LaeA.

Fungal velvet and mammalian Rel homology domain (RHD) share a common structure and might even have a common evolutionary ancestor [6]. The genetic architecture of these gene families presumably results from gene duplication events with subsequent codon changes for sub-functionalization of original genes [36]. Thus, additional functions or interactions to other proteins or DNA can be developed. Mutations in promoters or signal sequences can further alter temporal or spatial protein concentration levels within fungal cells through modulated gene expression [37]. DNA acquisition for an intrinsically disordered domain is only found within the fungal *velB* gene family, but neither in other velvet genes nor in the RHD domain family.

Higher differentiated Asco- or Basidiomycota mostly contain only one VelB ortholog with IDD. In contrast, members from the phyla Chytridiomycota and Zygomycota have more than one VelB-like isoform where one of them is predicted to be (nearly) continuous, whereas the other isoform mostly has a short interruption of the velvet domain. The VelB copy without IDD might have been lost during evolution. The insertion site of the *velB* DNA for the IDD is conserved during evolution. This supports that the original integration event happened in a common fungal ancestor and was further evolved to the current high diversity of IDD lengths and sequences. Accordingly, chimeric VelB proteins with the IDD region of *A. fumigatus* or *V. dahliae* are functional. The presence of an IDD in the VelB protein is presumably more important than the protein sequence. The VelB IDD involvement in VosA heterodimer formation is conserved between *A. nidulans* and *V. dahliae*, although the Vel2-Vos1 function might differ in the plant pathogen [14].

*A. nidulans* VelB-VeA supports sexual development but delays asexual sporulation as default mode of VelB heterodimer formation for fungal differentiation. Asexual development is accelerated by illumination, which is promoted by VelB-VosA heterodimer through increased dehydroaustinol synthesis [19,38,39]. Conidiospore numbers or production of the sporulation induction signal dehydroaustinol [35] are significantly reduced in strains without VelB IDD. Consistently, the phenotype of this strain is reminiscent to the appearance of deletion strains impaired in several genes for upstream developmental regulators of conidiation [24,40]. The IDD-dependent VelB-VosA heterodimer further promotes spore survival, whereas deletion of the IDD in *velB* results in reduced spore survival, as described for spores of *velB* or *vosA* deletion strains [41]. The IDD-dependent VelB-VosA heterodimer also triggers biosynthesis of secondary metabolites including xanthones from the *mdp* gene cluster [26,42,43] or the phytotoxin chicorine produced by the nonreducing polyketide synthase CicF [44–47]. VelB-VosA targets the *mcrA* gene encoding another global regulator for secondary metabolism. Accordingly, the *velB^ΔIDD^* and a *mcrA* deletion strain share the reduced sporulation phenotype, loss of spore viability and increased sterigmatocystin production [48].

More than 10% of annually harvested crops are spoiled by fungi and their bioactive metabolites, with a prominent role of the cancerogenic aflatoxin family [49,50]. The *velB^ΔIDD^* strain forms five-fold increased amounts of the aflatoxin family compound sterigmatocystin compared to wildtype. VelB-VeA together with methyltransferase LaeA are activators for sterigmatocystin biosynthesis [18], whereas VelB-VosA reduces this metabolite. Formations of VelB-VeA and presumably VelB-VeA-LaeA are enhanced in absence of the VelB IDD, whereas VelB-VosA heterodimer formation is nearly abolished, which elevates sterigmatocystin levels (Fig 6). This is consistent with increased expression of biosynthetic genes and sterigmatocystin production in ascospores of a *vosA* deletion strain [51]. The comparative analysis of *A. nidulans* wildtype with different combinations of velvet deletion strains revealed that each strain produced a different secondary metabolite peak profile (Fig 5). This suggests that the IDD fine-tunes fungal secondary metabolism and provides a valid tool to dissect the impact of the different VelB heterodimers on fungal secondary metabolite control.

The acquisition of the VelB IDD provides a protein destabilizing function, in addition to the control of heterodimer partner selection. Half-lives of fungal transcription factors controlling multiple genes in different fungal differentiation or pathogenicity programs vary considerably [52–54].

The interplay between expression of the *velB* gene and IDD-mediated protein stability determines the cellular VelB levels and adjusts fungal development and secondary metabolism. The ubiquitin-mediated 26S proteasomal degradation system reduces half-lives of various proteins with internal disordered domains of more than 40 residues [55,56]. The different VelB IDDs of the fungal kingdom share numerous serine residues (Fig S2). These are potential sites of phosphorylation as priming reaction for subsequent ubiquitination and degradation [57,58]. Posttranslational modifications as well masking or demasking of the IDD could provide molecular mechanism to increase or decrease VelB stability and shift the ratio of VelB protein complexes in response to different environmental stimuli in *A. nidulans* (Fig 6). Stability of the VelB interaction partner VeA depends on a complex interplay between the destructing ubiquitinating F-box23 containing E3 cullin-RING ligase for labeling for proteasomal degradation and the reversal stabilization by the deubiquitinating enzyme UspA [29,59]. The VelB IDD presumably interferes with numerous protein interactions, because interaction partners with different cellular functions were exclusively identified in pull downs of VelB^ΔIDD^-GFP but not with VelB-GFP. Many of these carry a conventional nuclear localization signal [34]. The IDD could provide VelB conformational flexibility and structural plasticity as variable hub, which allows the interaction with different proteins during various environmental conditions. Different interaction partners control or adjust cellular locations of distinct VelB complexes within or outside of the nucleus. This is illustrated by the finding that a VelB protein without IDD and therefore unable to interact to VosA is unable enter the nucleus, when the VeA protein is also missing in the cell.

The IDD mediated proportion of cellular VelB heterodimers therefore includes two control levels: (i) the IDD selects between VeA and VosA and (ii) the IDD stabilizes or destabilizes VelB as heterodimer binding partner. This allows to respond to different environmental stimuli in favor of specific fungal developmental programs linked to the appropriate secondary metabolism.

Here we discovered a novel mechanism for an intrinsically disordered domain within a velvet transcription factor, which has been specifically introduced and further evolved within the VelB protein family in the fungal kingdom. Velvet proteins control fungal defense, including the entire genetic network of fungal development, virulence and secondary metabolism, whereas the mammalian NF-ᴋB proteins with a similar DNA binding fold are relevant for infection or immune defense. The VelB IDD coordinates and fine-tunes the fungal chemical language by controlling the ratio of VelB heterodimer formation and their stability. This might be a promising starting point for a better understanding of fungal communication. In addition, it will be interesting to examine whether there will be IDDs with similar functions for heterodimeric transcription factors in other organisms than the fungi.

## Material and Methods

### Strains and growth conditions

*A. nidulans* strains were cultivated in liquid or solid minimal medium (MM) [60] in light with oxygen supply inducing asexual development or in darkness with limited oxygen supply by sealing the plates with parafilm inducing sexual development. For details see [24]. *V. dahliae* strains were cultivated as described in [14].

### Plasmid and strain preparation

All strains used in this study are listed in Table S7. A list of plasmids used in this study is shown in Table S8, and oligonucleotides in Table S9. Genomic DNA of FGSC A4 (*A. nidulans* WT, *veA*^+^) was used as template for amplification of DNA fragments for plasmid constructions. Gene targeting by homologous recombination was performed with recyclable marker (RM) cassettes [61]. Amplified DNA fragments and recyclable marker cassettes, which were excised from pME4304 and pME4305 with *Sfi*I, are called natRM and phleoRM, respectively, were cloned into the *EcoR*V multiple cloning site of pBluescript SK(+) using a seamless cloning reaction (Invitrogen). Transformation of plasmid excised cassettes into *A. nidulans* was performed by polyethylene glycol-mediated protoplast fusion as described earlier [62]. *V. dahliae* was transformed as described in [63]. Transformation of plasmids into *E. coli* was conducted as described before [64,65]. *E. coli* strains were cultivated in lysogeny broth (LB) (Bertani, 1951) medium (1% (w/v) tryptophan, 0.5% (w/v) yeast extract, 1% (w/v) NaCl).

### Plasmid and strain construction of *velB^ΔIDD^*

For construction of a *velB^ΔIDD^* strain 1 the 5’ flanking region and half of the velvet domain of the *velB* gene (to exclude the IDD) was amplified with SR110/SR109. The second half of the *velB* velvet domain was amplified with SR108/SR111. The 3’ flanking region was amplified with SR112/SR113. These fragments and the phleoRM cassette were cloned to pBluescript SK(*) resulting in pME4686. The *velB^ΔIDD^* cassette was excised with *Pme*I and transformed to AGB551 resulting in AGB1131, into AGB1066 giving AGB1140 and AGB1057 giving AGB1142.

### Plasmid and strain construction of *velB:gfp*

*gfp* was amplified with primers SR18/SR20 from pME4292. The 5’ flanking region and the *velB* gene was amplified with SR05/SR24. The 3’ flanking region was amplified with SR07/SR08. These three fragments and the natRM marker cassette were cloned into pBluescript SK(+), resulting in pME4687. The *velB:gfp* cassette was excised with *Pme*I, followed by transformation in AGB551 and AGB1066, which resulted in the AGB1132 and AGB1190, respectively.

### Plasmid and strain construction of *velB^ΔIDD^*:*gfp*

Fragments from constructing pME4686 were used for the assembly of this construct, but the second half of the *velB* velvet domain was amplified with primers SR108/SR24. The *velB* velvet domain fragments amplified with SR109/SR110 and SR108/SR24 were joined by fusion PCR. This desired fragment and *gfp* (amplified with SR18/SR20 from pME4292), the 3’ flanking region and the phleoRM marker cassette were cloned into pBluescript SK(+) giving plasmid pME4688. The *velB^ΔIDD^*:*gfp* cassette was excised with *Pme*I, followed by transformation in AGB551 and AGB1066, which resulted AGB1133 and AGB1191, respectively.

### Plasmid and strain construction of *velB^AfIDD^* complementation

The 3’ flanking region was amplified with SR112/SR113 and cloned into the *Eco72I* restriction site of pME4319, resulting in pME4689. The 5’ flanking region and half of the velvet domain of the *A. nidulans velB* gene (until the IDD) was amplified with primers SR110/SR253. The 0.3 kb IDD of *A. fumigatus velB* was amplified with SR266/SR267 from *A. fumigatus* Afs35 gDNA. The second half of the *A. nidulans velB* velvet domain was amplified with SR108/SR111. All fragments were joined by fusion PCR. The fused fragment was cloned into the *Swa*I restriction site of pME4689, resulting in pME4690. The *velB^AfIDD^* complementation cassette was excised with *Pme*I, followed by transformation in AGB1133, resulting in AGB1134.

### Plasmid and strain construction of *velB^VdIDD^* complementation

The 0.4 kb IDD of *V. dahliae vel2* was amplified with SR254/SR255 from *V. dahliae* JR2 gDNA. The fragments SR110/253, SR254/255 and SR108/111 were joined by fusion PCR. The fused fragment was cloned into the *Swa*I restriction site of pME4689, resulting in pME4691. The *velB^VdIDD^*complementation cassette was excised with *Pme*I, followed by transformation in AGB1133, yielding AGB1135.

### Plasmid construction of pME4692 for recombinant expression of *velB^ΔIDD^*:*his* in *E. coli*

The fragments for the two parts of the *velB* velvet domain were amplified from pME3815 with primers JG45/SR109 and JG46/SR108. These fragments were joined by fusion PCR containing an overhang for the *Nco*I or *Xho*I restriction site, respectively. After subcloning in the pJET plasmid, the construct was excised with *Nco*I and *Xho*I and cloned into *Nco*I/*Xho*I site of pETM-13, which contains a sequence encoding a C-terminal His-tag, resulting in pME4692. The plasmid was transformed to Rosetta II *E. coli* strain (NOVAGEN^®^, MERCK) for recombinant protein expression.

### Plasmid and strain construction of *vosA:ha*

The 5’ flanking region and the *vosA* gene were amplified with primers SR76/SR201, introducing a sequence encoding a HA (<u>h</u>emaglutinin <u>a</u>ntigen) tag. The 3’ flanking region was amplified with primers SR49/SR75. The two fragments and the phleoRM marker cassette were cloned into the *EcoR*V multiple cloning site of pBluescript SK(+), resulting in pME4693. The *vosA:ha* cassette was excised with *Pme*I and transformed into AGB1132 and AGB1133, resulting in AGB1136 and AGB1137, respectively.

### Plasmid and strain construction of *veA:ha*

The *veA* 5’ flanking region was amplified with *veA* using KT197/KT166. This template was used for another PCR introducing a sequencing encoding a HA tag using KT197/KT163. The seamless cloning kit was used to ligate the 5’:*veA:ha* fragment with the natRM and the *veA* 3’ flanking region (KT142/KT198) into pBluescript SK(+) resulting in pME4748. The *veA:ha* cassette was excised with *Pme*I and transformed into AGB1132 and AGB1133, resulting in AGB1149 and AGB1150, respectively.

### Plasmid and strain construction of *V. dahliae vel2^ΔIDD^:GFP*

The construction of a *VEL2* strain without *IDD* fused to *GFP* was conducted in several steps. In the first step, the primers AO74 and AO75 were used to amplify the 5’ flanking region of the gene and *VEL2* until the start of the *IDD* from gDNA. In another PCR, AO76 and AO77 were utilized to amplify the part of *VEL2* downstream of the *IDD* from gDNA. As AO75 was constructed with an overhang to *VEL2* after the *IDD*, the two fragments were fused by PCR using AO74 and AO77. The fragment was ligated in the *Eco*RV-linearized pPK2 and the resulting plasmid was named pME5075. In the next step, pME5075 was cut with *Xba*I and the 3’ flanking region of *VEL2* was amplified with AO78 and AO79. Ligation of pME5075 and the PCR fragment resulted in a plasmid named pME5076. In the last step, the 5’ flanking region and *VEL2* without *IDD* and stop codon were amplified with AO167 and AO168 from pME5076. The 3’ flanking region was amplified with the primers AO169 and AO170. *GFP* (without start codon) and a flexible linker (protein sequence GGSGG) were amplified from pME4990 with AO165 and RH514. The hygromycin resistance cassette was amplified from the same plasmid with the primers RH590 and RO4. The *GFP*-linker fragment and the hygromycin marker were fused by PCR with the primers AO165 and RO4. The three generated fragments were ligated into pME4564 cut with *Eco*RV and *Stu*I. The created plasmid was named pME5077. The wild type (JR2) was transformed with the plasmid resulting in VGB468. The gDNA of the constructed strain was treated with *Hinc*II and *Pst*I and tested by Southern hybridization with the 5’ flanking region as probe. The strain was also confirmed by cutting the gDNA with *Eco*RI and conducting a Southern hybridization with the 3’ flanking region as probe.

### Extraction of fungal genomic DNA

Extraction of *A. nidulans* genomic DNA from liquid cultures was performed as described before [24]. Genomic DNA of *V. dahliae* was extracted as described in [66].

### Isolation of fungal RNA and cDNA synthesis and quantitative real-time-PCR

Fungal RNA was isolated from vegetative cultures. Subsequent cDNA synthesis and quantitative real-time PCR was performed as described before [24]. cDNA synthesis for checking the expression of *velB* was performed with 0.8 µg RNA and for checking the expression of *aus* genes with 2.0 µg RNA. Primers used for qRT-PCRs are listed in Table S10.

### Sterigmatocystin isolation

1*10^5^ spores were distributed on solid MM and grown for three or seven days at 37°C in light or dark. Two agar plugs were excised with a 50 mL centrifuge tube (SARSTEDT), which were cut into small pieces. The agar pieces were transferred into 50 mL Falcon tubes and six glass beads and 3 mL H_2_O were added. Samples were shaken roughly for 30 min at room temperature (rt). Subsequently, 3 mL chloroform were added and samples were incubated for another 30 min at rt. This was followed by centrifugation at 1000 rpm for 10 min at 4°C for phase separation and the lower chloroform metabolite-containing chloroform phase was transferred into glass tubes and evaporated o/n at rt under the hood.

### Thin layer chromatography

Samples of sterigmatocystin isolation were resuspended in 50 µL methanol and three times 5 µL of isolated sterigmatocystin per sample was applied spot-wise to pre-coated SIL G/UV254 Polygram® DC-foil TLC-sheets (MACHEREY-NAGEL) (thin layer chromatography plates). TLC plates were run in 1:4 (v/v) acetone:chloroform for 40-50 min, dried for 5 min and photographed at 366 and 254 nm with a Camag TLC Visualizer 2 system (CAMAG). Afterwards, TLC plates were sprayed with 20% (v/v) aluminum chloride in 95% (v/v) ethanol and incubated at 70°C for 10 min. Derivatized TLC plates were photographed again at 366 and 254 nm UV light and white light with a Camag TLC visualizer 2 system and processed with the winCATS 1.4.4 software (CAMAG).

### Secondary metabolite extraction

For secondary metabolite extraction 1*10^5^ spores were distributed over the whole agar plate and grown for three days at 37°C in light, promoting asexual development. Subsequently, four round agar pieces with a diameter of 2.5 cm were excised and homogenized with a syringe. Metabolites were extracted with a mixture of 8 ml water and 8 ml ethyl acetate overnight. After centrifugation, 5 ml of the organic phase were dried, dissolved in 500 µl methanol and subjected to LCMS analysis.

### LCMS analysis of secondary metabolites

The reconstituted metabolites were analyzed using a Q Exactive^TM^ Focus orbitrap mass spectrometer coupled with an UltiMate 3000 HPLC (THERMO FISHER SCIENTIFIC). 1 µl of each sample was injected on a HPLC column (Acclaim^TM^ 120, C18, 5 µm, 120 Å, 4.6 x 100 mm (THERMO FISHER SCIENTIFIC)) applying a linear acetonitrile/0.1% (v/v) formic acid in H_2_O/0.1% (v/v) formic acid gradient (from 5% to 95% (v/v) acetonitrile/0.1 formic acid in 20 min, plus additional 10 min with 95% (v/v) acetonitrile/0.1 formic acid) with a flow rate of 0.8 ml/min at 30°C. The measurements were performed in a mass range of 70-1050 m/z in positive or negative mode. Data analysis was performed with Thermo Scientific Xcalibur 4.1 (Thermo Fisher Scientific) and FreeStyle™ 1.4 (Thermo Fisher Scientific).

### Phenotypic analyses of fungal strains

For phenotypic analyses either 2000 spores were spotted in the middle of the agar plate or 10^5^ spores were distributed over the whole plate. Agar plates were incubated for three or seven days under either asexual or sexual development promoting conditions. Photomicrographic pictures of *A. nidulans* colonies were obtained by utilization of an Axiolab microscope (ZEISS) or with the help of a binocular microscope SZX12-ILLB2-200 (OLYMPUS). Visualization was performed with an OLYMPUS SC30 digital camera and pictures were processed with the cellSens software (OLYMPUS).

### Fluorescence microscopy

Fluorescence microscopy was conducted with a Zeiss AxioObserver Z.1 inverted confocal microscope, equipped with Plan-Neofluar 63x/0.75 (air), Plan-Apochromat 63x/1.4 oil and a Plan-Apochromat 100x/1.4 oil objectives (ZEISS) and a QuantEM:512SC camera (PHOTOMETRICS). Pictures were processed with the SlideBook 6.0 software package (INTELLIGENT IMAGING INNOVATIONS).

For fluorescence microscopy 2000 spores per strain were inoculated in 8-well borosilicate cover glass system (THERMO FISHER SCIENTIFIC) in 400 μL liquid MM or on agar slants for vegetative growth and grown for 18h at 30°C. Nuclei were visualized via staining with 0.1% (w/v) 4’,6’-<u>dia</u>midino-2-<u>p</u>henylindole (DAPI) (ROTH) and incubation for 10 min at rt prior to microscopy.

### Conidiospore quantification

Conidiospore numbers were determined by utilization of Coulter Z2 particle counter (BECKMAN COULTER) or with a Thoma cell counting chamber (hemocytometer) (PAUL MARIENFELD).

### Protein isolation

1*10^7^ spores were inoculated in MM and strains were grown vegetatively in submerged cultures. For protein isolation of asexually or sexually grown cultures, strains were grown vegetatively for 24h and subsequently shifted onto solid MM plates and grown for indicated time points in light or dark. For cycloheximide assay, after 24h of vegetative growth, 250 µL cycloheximide (10 mg/mL) was added to the cultures, which were incubated for 0 to 5h prior protein extraction. Mycelia were collected and protein crude extracts were obtained as described [24].

### Immunoprecipitation with GFP-tagged fusion proteins

5*10^8^ spores of *A. nidulans* strains were inoculated in 500 mL MM and grown for 24h in submerged cultures (vegetative samples) at 37°C. Sexual samples were shifted afterwards to solid MM, sealed with Parafilm^®^ and incubated for 48h in the dark at 37°C. Protein GFP pull downs were conducted by employing GFP-Trap^®^_A beads from (CHROMOTEK) as described earlier [47,54].

### Pull downs with HA-tagged fusion proteins

Protein HA pulldowns were conducted by utilizing Monoclonal Anti-HA-Agarose beads A2095 (SIGMA ALDRICH). *A. nidulans* strains were inoculated in a concentration of 5*10^8^ spores in 500 mL MM and vegetatively grown for 24h in submerged cultures at 37°C. Mycelia were treated like for GFP trap. HA beads were washed with PBS, equilibrated with B^+^ buffer, added to the filtered supernatant and incubated rotating for 3h or o/n at 4°C. The supernatant with the HA beads was loaded onto fresh Poly-Prep^®^ Chromatography Columns (BIO-RAD), which were equilibrated with B^+^ buffer before, and washed with W500 buffer. Proteins were eluted with 100 µL 0.1 M glycine pH 2.5 and 2.5 µL Tris pH 10.4 and analyzed by western experiments.

### SDS PAGE and western experiments

SDS-polyacrylamide gel electrophoresis (SDS-PAGE) and western experiments were performed as described in [54]. Primary antibodies anti-GFP (sc-9996, SANTA CRUZ BIOTECHNOLOGY), anti-tubulin (T0926, SIGMA-ALDRICH), 6x-His Tag Monoclonal Antibody (R930-25, THERMO FISHER SCIENTIFIC), anti-GST (Z-5, SANTA CRUZ BIOTECHNOLOGY) and anti-hemagglutinin (anti-HA, clone HA-7; SIGMA-ALDRICH) were diluted in TBST-M (TBST buffer, supplemented with 5% (w/v) skim milk powder) and incubated o/n at 4°C. secondary antibodies, either horseradish peroxidase-coupled rabbit antibody (G21234, INVITROGEN) or mouse antibody (115-035-003, JACKSON IMMUNORESEARCH) in a dilution of 1:1000 in TBST-M. As loading control membranes were stained with Ponceau staining.

### *In vitro* co-immunoprecipitation of recombinant expressed proteins

For *in vitro* co-IPs, LB was inoculated with a pre-culture of the respective *E. coli* cells and incubated on a rotary shaker. Protein expression was induced with 1 mM IPTG at OD_600_ of 0.8. After o/n incubation at 20°C on a rotary shaker, cells were harvested by centrifugation at 4000 rpm for 20 min at 4°C, washed with buffer W (100 mM Tris pH 8, 150 mM NaCl, 1 mM EDTA) and centrifuged again. For protein purification and *in vitro* co-IP the cell pellets were resuspended in buffer W supplemented with 1 mM PMSF and cell lysis was performed by sonication. VosA-GST cells were centrifuged at 20000 rpm for 30 min at 4°C. The supernatant was incubated with GST beads for 3h rotating at 4°C. After incubation the lysate was centrifuged at 1000 rpm for 2 min at 4°C and washed with buffer W + 1 mM IPTG. The beads were divided into two 15 mL reaction tubes, the supernatant of lysed cells containing either pME3815 or pME4692 was added and incubated for 2h on a rotator at 4°C followed by centrifugation at 1000 rpm for 2 min at 4°C. The beads were washed twice with buffer W1 mM IPTG, transferred to 1.5 mL reaction tubes and washed twice again. The supernatant was removed carefully from the beads, 3x SDS sample buffer was added and boiled for 10 min at 95°C. The samples were analyzed by 12% SDS PAGE and experiments for specific detection of His- or GST-tagged proteins.

### Tryptic protein digestion and peptide analysis with LCMS

Protein samples were separated by SDS PAGE. The gel was incubated in fixing solution (40% (v/v) ethanol, 10% (v/v) acetic acid) for 1 h, washed with dH_2_O and incubated o/n in colloidal Coomassie [67] (5% (w/v) aluminum-sulfate-(14-18)-hydrate, 10% (v/v) methanol, 0.1 (w/v) Coomassie Brilliant Blue G-250, 2% (v/v) ortho-phosphoric acid). Each lane was excised completely and cut into small pieces of approximately 2 mm. Protein tryptic digestion was performed as described earlier using Sequencing Grade Modified Tryspin (PROMEGA) [24,68], followed by stage tip purification as described in [69,70].

### Identification of proteins by LCMS/MS

Peptide solutions were analyzed with mass spectrometry coupled to liquid chromatography (LCMS) at the LCMS facility at the Institute of Microbiology and Genetics as described in [14,29,47]. The samples were subjected to reverse phase liquid chromatography for peptide separation using an RSLCnano Ultimate 3000 system (Thermo Fisher Scientific). Peptides were loaded on an Acclaim PepMap 100 pre-column (100 µm x 2 cm, C18, 5 µm, 100 Å; Thermo Fisher Scientific) with 0.07% trifluoroacetic acid at a flow rate of 20 µL/min for 3 min. Analytical separation of peptides was done on an Acclaim PepMap RSLC column (75 µm x 50 cm, C18, 2 µm, 100 Å; Thermo Fisher Scientific) at a flow rate of 300 nL/min. The solvent composition was gradually changed within 94 min from 96 % solvent A (0.1 % formic acid) and 4 % solvent B (80 % acetonitrile, 0.1 % formic acid) to 10 % solvent B within 2 minutes, to 30 % solvent B within the next 58 min, to 45% solvent B within the following 22 min, and to 90 % solvent B within the last 12 min of the gradient. Eluting peptides were on-line ionized by nano-electrospray (nESI) using the Nanospray Flex Ion Source (Thermo Fisher Scientific) and transferred into an Orbitrap mass spectrometer (Thermo Fisher Scientific). Full scans in a mass range of 300 to 1650 m/z were recorded at a resolution of 30,000 followed by data-dependent top 10 HCD fragmentation at a resolution of 15,000 (dynamic exclusion enabled). LC-MS method programming and data acquisition was performed with the XCalibur 4.0 software (Thermo Fisher Scientific). Protein database searches and subsequent data analyses were carried out using MaxQuant (V.2.2.0.0) and Perseus (V.2.0.7.0) respectively.

## Acknowledgements

The authors thank Verena Große, Nicole Scheiter and Helen Stupperich and Sarah E. Eubanks for their technical assistance. We would also like to thank Prof. Dr. Ralf Ficner and Dr. Piotr Neumann for providing the structure and the resulting figure.

## Supplementary Information Captions

**Fig S1: *A. nidulans* VelB is interrupted by a 99 amino acid intrinsically disordered domain (IDD).** Structure prediction of *A. nidulans* VelB using the DISOPRED program [71]. The velvet domain (VD: in red) for DNA binding and dimerization is interrupted by an intrinsically disordered domain (IDD, yellow) of 99 amino acids. The dotted line represents the cut-off of 0.5 of the confidence score (disorder probability).

**Fig S2: Within the fungal kingdom VelB IDDs vary in length and amino acid sequence.** Alignments of deduced protein sequences for the *A. nidulans* VelB intrinsically <u>d</u>isordered <u>d</u>omain (IDD) (yellow box) against VelB proteins of indicated fungal species as examples for the **(A)** Ascomycota, **(B)** Basidiomycota, **(C)** Zygomycota or **(D)** Chytridiomycota. Serine residues as putative phosphorylation sites are highlighted in pink. The VelB IDDs of Ascomycota contain the conserved Motif_IDD_ (turquoise box). The N-terminal boundary of the VelB <u>v</u>elvet <u>d</u>omain (VD) is conserved in all analyzed fungal genomes (red box).

**Fig S3: VelB IDD variations between different fungal divisions.** X-axis represents IDD lengths in amino acid residues and y-axis the number of fungal species within the four divisions (corresponding orthologs and deduced IDD information in Table S1 and S2).

**Fig S4: Relative expression of *velB:gfp* transcripts is similar with or without IDD.** Strains were grown in submerged cultures for 24h, shifted to solid minimal medium and incubated for 6 or 18h in the light at 37°C. qRT-PCR analyses indicate that transcription of the truncated *velB^ΔIDD^:gfp* gene is similar to the full length *velB:gfp* gene. Data from two biological and three technical replicates are shown. *15S rRNA* and *h2A* were used as references for normalization.

**Fig S5: VelB IDD is required for selective heterodimer formation with VosA and subsequent nuclear localization. (A)** Fluorescence micrographs of 18h vegetatively grown wild type (wt) and constitutively expressing GFP (OE GFP) strains as control for microscopy. **(B)** Western blot experiment for VelB-GFP and VelB^ΔIDD^-GFP in wild type (wt), *veA* (Δ*veA*) and *vosA* (Δ*vosA*) single and double deletion background used in microscopy. The crude extracts were obtained from mycelium of vegetatively grown cultures (18h). The pixel density was calculated using the BioID software (Vilber Lourmat), normalized against the loading control Ponceau and calculated relative to VelB-GFP (red bars) or VelB^ΔIDD^-GFP (blue bars) in wild type background. **(C+D)** Fluorescence microscopy of the same strains after growth in 18h sexual **(C)** and 18h asexual **(D)** development inducing conditions. White arrows indicate nuclei. Size bar represents 10 µm.

**Fig S6: The VelB IDD allows selective VelB-VosA formation in *A. nidulans* and *V. dahliae*. (A)** GFP pull downs of VelB-GFP and VelB^ΔIDD^-GFP from vegetatively grown *A. nidulans* cultures followed by LC-MS/MS revealed that the lack of the IDD leads to an increased number of putative interaction partners (Table S3). The three proteins found in both GFP pulldowns (left column) are the corresponding bait protein itself, the catalase CatB and the velvet protein VeA. The single protein, which is only recruited by VelB-GFP (middle column) is the velvet protein VosA. The diagram depicts the predicted cellular function or localization of the 32 putative interaction partners of VelB^ΔIDD^-GFP. Numbers represent proteins identified in at least two out of three biological repetitions with MS/MS counts ≥ 4, unique peptides ≥ 3 and LFQ intensity ≥ 20 and which were absent in the control strain. **(B)** *V. dahliae* Vel2-GFP or Vel2^ΔIDD^-GFP strains were used for GFP pull down experiments, followed by LC-MS analysis. The x-axis are fold-differences of LFQ values of the strains indicated (mean of three independent experiments). The y-axis shows the -Log p value resulting from a two-tailed t-test. Significantly interacting proteins are found in the upper right part of the plot (Table S4). Missing values were replaced four times with imputed values and proteins, which showed a significant interaction in all 4 repetitions, are colored (Vel1: yellow, Vel2 respectively Vel2^ΔIDD^: orange, Vos1: turquois, other proteins: dark blue, GFP: lime). **(C)** Comparison of significantly enriched proteins in Vel2 and Vel2^ΔIDD^ pull downs. Among the 39 significantly enriched proteins in Vel2 and four enriched proteins in Vel2^ΔIDD^ are two overlaps, Vel1 and Vel2. Within the proteins enriched in Vel2 is also Vos1. Vos1 was not significantly enriched in the Vel2^ΔIDD^ strain.

**Fig S7: *In vitro* VelB binds full length VosA independently of the IDD.** *In vitro* co- immunoprecipitation of VosA-GST and VelB-His or VelB^ΔIDD^-His fusion proteins recombinantly expressed in *E. coli.* Specific detection of fusion proteins was achieved with α-His or α-GST antibodies. VosA-GST (violet arrow) pulls VelB-His (red arrow) and VelB^ΔIDD^-His (blue arrow) fusion proteins. SN = supernatant of lysed cells after centrifugation.

**Fig S8: Heterodimer formation of VosA with VelB missing IDD is restored in absence of VeA. (A)** Western experiment of 24h vegetative and six hour light incubated *A. nidulans* strains shows GFP signals of *velB:gfp* and *velB^ΔIDD^:gfp* in wild type and Δ*veA* background. OE = GFP over expression. **(B)** Summary of identified velvet proteins VelB, VeA and VosA in each GFP pull down (*velB:gfp, velB:gfp/*Δ*veA, velB^ΔIDD^:gfp* and *velB^ΔIDD^:gfp/*Δ*veA*). Plus (+) means identified as interaction partner in at least two replicates, minus (-) means not identified or only with MS/MS or unique peptide counts lower than one (Table S5).

**Fig S9: VelB-VosA heterodimer formation is reduced in absence of the IDD and *veA.*** Western experiment of CoIP samples from 20h vegetative mycelium of indicated strains. Co- IPs were conducted with GFP and HA beads and the elution fractions (E) were subjected to SDS PAGE following western experiments using anti-GFP and anti-HA antibodies.

**Fig S10: Phenotypes of *A. nidulans veA* or *vosA* deletion strains are epistatic to *velB^ΔIDD^*in corresponding double mutant strains. (A)** Phenotypes of wildtype (wt), *velB* deletion (Δ*velB*), *velB^IDD^* deletion (*velB^ΔIDD^*) and complementation strains with the IDD of *A. nidulans*, *A. fumigatus* or *Verticillium dahliae* into *velB^ΔIDD^* (*velB^AnIDD^:sgfp*, *velB^AfIDD^* and *velB^VdIDD^*) on solid minimal medium. 2000 spores were spotted in the middle of the plate and grown in light for 3 or 7 days (d) at 37°C. PMG photomicrograph, scale bar = 100 µm. **(B)** Differential expression of genes, essential for the synthesis of austinol and dehydroaustinol, in Δ*velB* and *velB^ΔIDD^* strains. mRNAs were extracted from mycelia grown in submerged cultures shaking at 37°C for 20h in light. For normalization of gene expression the reference gene *h2A* was used and the expression of all samples was calibrated towards the wt. Data presented are the averages and standard deviations derived by at least three biological replicates, each with at least three technical replicates. Statistics were performed by Student’s t test: * p < 0.05, ** p < 0.01 and *** p < 0.001, n.s.: not significant.

**Fig S11: Sterigmatocystin production depends on the VelB IDD. (A)** Secondary metabolite production is altered in the *velB^ΔIDD^* strain. <u>H</u>igh <u>p</u>erformance liquid <u>c</u>hromatography coupled with <u>m</u>ass <u>s</u>pectrometry (HPLC-MS) was performed with metabolites extracted after 3 days of growth in light. Metabolites were isolated of wildtype (wt), velB deletion (Δ*velB*), *velB^IDD^* deletion (*velB^ΔIDD^*) and *velB^ΔIDD^*complementation (*velB^AnIDD^:gfp*). The chromatogram shows identified secondary metabolites at 250 nm, indicated with numbers (peak 3: austinol, peak 4: dehydroaustinol, peak 7: sterigmatocystin, peak 11: emericellin) and listed in Table S6. **(B)** <u>T</u>hin layer <u>c</u>hromatography (TLC) shows increased sterigmatocystin production in the *velB^ΔIDD^* strains (red arrow) after 3 or 7 days of growth in the light. The signals of the TLC plate were detected at 366 nm after derivatization with AlCl_3_. ST = Sterigmatocystin standard. **(C)** Quantification of the sterigmatocystin signal relative to wt. Error bars indicate the SEM. Statistics were performed by Student’s t test: * *p* < 0.05.

**Fig S12 Extracted ion chromatograms (EICs) of detected peaks.** The EICs show the identified metabolites (M ± 5 ppm) from Table S6 and Fig S13 measured from 3 days asexual and 7 days sexual developmental samples in positive ionization mode [M+H]^+^. Masses are given in Table S6. Cichorine (1), F-9775A/B (2), Austinol (3), Dehydroaustinol (4), Arugosin H (5), Emericellamide C (6), Sterigmatocystin (7), Emericellamide E (8), Terrequinone A (9), Arugosin A (10), Emericellin (11), Shamixanthone (12), Epishamixanthone (13).

**Fig S13: Overview of identified metabolites and their structures from Table S6.**

**Table S1: Summary on IDD information (Excel file)**

**Table S2: Collection of fungal genomes with multiple VelB homologs (Excel file)**

**Table S3: Putative interaction partners identified in the GFP pull down of VelB-GFP and VelB^ΔIDD^-GFP expressing strains.** Proteins were identified in at least two out of three biological repetitions with MS/MS counts ≥ 4, unique peptides ≥ 3 and LFQ intensity ≥ 20 and which were absent in the control strain. Proteins with underlined AN numbers contain a nuclear localization signal predicted with cNLS mapper [34] with a score ≤ 5 (indicating a localization in both, nucleus and cytoplasm), Sys. Name = systematic name, Std. name = standard name, unchar. = uncharacterized. Descriptions were obtained and adapted from FungiDB and NCBI [72,73]

**Table S4: Proteins significantly enriched with *V. dahliae* Vel2-GFP and Vel2^ΔIDD^-GFP, their predicted domains and potential *A. nidulans* counterparts.** For identification of proteins and their conserved domains, EnsemblFungi was used [74].

**Table S5: VelB-VosA heterodimer formation is independent of the IDD in a *veA* deletion background.** Identification of VeA, VosA and LaeA as interaction partner of VelB with or without IDD in wild type or *veA* deletion (Δ*veA*) background. Thresholds were set as followed: MS/MS counts ≥ 3, LFQ intensity ≥ 20, unique peptides ≥ 2 and identified proteins are absent in the GFP OE control strain in both biological replicates. Std. name = systematic name, Sys. name = systematic name.

**Table S6: Secondary metabolites identified in this study.** Table corresponds to Fig 5, Fig S11, Fig S12 and Fig S13. PKS = polyketide synthase, NR-PKS = non-reducing polyketide synthase, NRPS = non ribosomal polyketide synthase, cic = chicorine, ors = orsellinic acid, aus = austinol, mdp = monodictophenone, eas = emericellamide, stc = sterigmatocystein, tdi = (tryptophane-derived) terrequinone gene clusters. Structures are depicted in Fig S13.

**Table S7: Fungal strains used in this study.** Strains were constructed by employment of recyclable marker cassettes (see material and methods section in the main text). These marker cassettes leave only a small six site (100 nucleotides) as scar after recycling of the marker off the genome. FGSC = Fungal genetics stock center, Kansas, USA.

**Table S8: Plasmids constructed and used in this study**. pBluescript SK+ was used as backbone for all plasmids constructed in this study, if not stated otherwise. ^P^ = promoter, ^t^ = terminator, ^R^ = resistance, natRM = recyclable nat^R^ resistance cassette from pME4304, phleoRM = recyclable phleo^R^ resistance cassette from pME4305, p.c. = personal communication; hyg^R^ = hygromycine resistance gene; Kan^R^ = kanamycine resistance.

**Table S9: Oligonucleotides used for amplifications and plasmid constructions.**

**Table S10: Primers used for qRT-PCR in this study.**

## References

1. Holehouse AS, Kragelund BB. The molecular basis for cellular function of intrinsically disordered protein regions. Nature Reviews Molecular Cell Biology 2023 25:3. 2023;25: 187–211. doi:10.1038/s41580-023-00673-0

2. Habchi J, Tompa P, Longhi S, Uversky VN. Introducing protein intrinsic disorder. Chem Rev. 2014;114: 6561–88. doi:10.1021/cr400514h

3. Wang C, Uversky VN, Kurgan L. Disordered nucleiome: Abundance of intrinsic disorder in the DNA- and RNA-binding proteins in 1121 species from Eukaryota, Bacteria and Archaea. Proteomics. 2016;16: 1486–1498. doi:10.1002/pmic.201500177

4. Smith NC, Kuravsky M, Shammas SL, Matthews JM. Binding and folding in transcriptional complexes. Curr Opin Struct Biol. 2021;66: 156–162. doi:10.1016/j.sbi.2020.10.026

5. Guo X, Bulyk ML, Hartemink AJ. Intrinsic disorder within and flanking the DNA-binding domains of human transcription factors. Pacific Symposium on Biocomputing. 2012. pp. 104–115. doi:10.1142/9789814366496_0011

6. Ahmed YL, Gerke J, Park H-S, Bayram Ö, Neumann P, Ni M, et al. The Velvet family of fungal regulators contains a DNA-binding domain structurally similar to NF-κB. Stock AM, editor. PLoS Biol. 2013;11: e1001750. doi:10.1371/journal.pbio.1001750

7. Bayram Ö, Braus GH. Coordination of secondary metabolism and development in fungi: the velvet familyof regulatory proteins. FEMS Microbiol Rev. 2012;36: 1–24. doi:10.1111/j.1574-6976.2011.00285.x

8. Myung K, Zitomer NC, Duvall M, Glenn AE, Riley RT, Calvo AM. The conserved global regulator VeA is necessary for symptom production and mycotoxin synthesis in maize seedlings by *Fusarium verticillioides*. Plant Pathol. 2012;61: 152–160. doi:10.1111/j.1365-3059.2011.02504.x

9. Gerke J, Braus GH. Manipulation of fungal development as source of novel secondary metabolites for biotechnology. Appl Microbiol Biotechnol. 2014/08/21. 2014;98: 8443–8455. doi:10.1007/s00253-014-5997-8

10. Schumacher J, Simon A, Cohrs KC, Traeger S, Porquier A, Dalmais B, et al. The velvet vomplex in the gray mold fungus *Botrytis cinerea*: impact of BcLAE1 on differentiation, secondary metabolism, and virulence. Mol Plant Microbe Interact. 2015;28: 659–674. doi:10.1094/MPMI-12-14-0411-R

11. Calvo AM, Lohmar JM, Ibarra B, Satterlee T. Velvet regulation of fungal development. 3rd ed. In: Wendland J, editor. The Mycota I: Growth, Differentiation and Sexuality. 3rd ed. Cham: Springer International Publishing Switzerland; 2016. pp. 475–497. doi:10.1007/978-3-319-25844-7_18

12. Lee M-K, Kwon N-J, Lee I-S, Jung S, Kim S-C, Yu J-H, et al. Negative regulation and developmental competence in Aspergillus. Sci Rep. 2016;6: 28874. doi:10.1038/srep28874

13. Müller N, Leroch M, Schumacher J, Zimmer D, Könnel A, Klug K, et al. Investigations on VELVET regulatory mutants confirm the role of host tissue acidification and secretion of proteins in the pathogenesis of *Botrytis cinerea*. New Phytologist. 2018;219: 1062–1074. doi:10.1111/nph.15221

14. Höfer AM, Harting R, Aßmann NF, Gerke J, Schmitt K, Starke J, et al. The velvet protein Vel1 controls initial plant root colonization and conidia formation for xylem distribution in Verticillium wilt. PLoS Genet. 2021;17. doi:10.1371/JOURNAL.PGEN.1009434

15. Becker K, Ziemons S, Lentz K, Freitag M, Kück U. Genome-Wide Chromatin Immunoprecipitation Sequencing Analysis of the *Penicillium chrysogenum* Velvet Protein PcVelA Identifies Methyltransferase PcLlmA as a Novel Downstream Regulator of Fungal Development. mSphere. 2016;1: e00149–16. doi:10.1128/mSphere.00149-16

16. Potoyan DA, Bueno C, Zheng W, Komives EA, Wolynes PG. Resolving the NFκB hetero-dimer binding paradox: Strain and frustration guide the binding of dimeric transcription factors. J Am Chem Soc. 2017;139: 18558–18566. doi:10.1021/jacs.7b08741

17. Park HS, Ni M, Jeong KC, Kim YH, Yu JH. The role, interaction and regulation of the velvet regulator VelB in Aspergillus nidulans. PLoS One. 2012;7: e45935. doi: 10.1371/journal.pone.0045935. doi:10.1371/journal.pone.0045935

18. Bayram O, Krappmann S, Ni M, Bok JW, Helmstaedt K, Valerius O, et al. VelB/VeA/LaeA complex coordinates light signal with fungal development and secondary metabolism. Science. 2008;320: 1504–6. doi:10.1126/science.1155888

19. Sarikaya Bayram O, Bayram O, Valerius O, Park HS, Irniger S, Gerke J, et al. LaeA control of velvet family regulatory proteins for light-dependent development and fungal cell-type specificity. Brakhage AA, editor. PLoS Genet. 2010;6: e1001226. doi:10.1371/journal.pgen.1001226

20. Gerke J, Köhler AM, Meister C, Thieme KG, Amoedo H, Braus GH. 8 Coordination of Fungal Secondary Metabolism and Development. Genetics and Biotechnology. Cham: Springer International Publishing; 2020. pp. 173–205. doi:10.1007/978-3-030-49924-2_8

21. Park HS, Yu JH. Genetic control of asexual sporulation in filamentous fungi. Curr Opin Microbiol. 2012;15: 669–677. doi:10.1016/j.mib.2012.09.006

22. Adams TH, Wieser JK, Yu JH. Asexual sporulation in *Aspergillus nidulans*. Microbiol Mol Biol Rev. 1998;62: 35–54.

23. Ni M, Yu J-H. A novel regulator couples sporogenesis and trehalose biogenesis in *Aspergillus nidulans*. d’Enfert C, editor. PLoS One. 2007;2: e970. doi:10.1371/journal.pone.0000970

24. Thieme KG, Gerke J, Sasse C, Valerius O, Thieme S, Karimi R, et al. Velvet domain protein VosA represses the zinc cluster transcription factor SclB regulatory network for *Aspergillus nidulans* asexual development, oxidative stress response and secondary metabolism. Copenhaver GP, editor. PLoS Genet. 2018;14: e1007511. doi:10.1371/journal.pgen.1007511

25. Troppens DM, Köhler AM, Schlüter R, Hoppert M, Gerke J, Braus GH. Hülle Cells of Aspergillus nidulans with Nuclear Storage and Developmental Backup Functions Are Reminiscent of Multipotent Stem Cells. Di Pietro A, editor. mBio. 2020;11. doi:10.1128/mBio.01673-20

26. Liu L, Sasse C, Dirnberger B, Valerius O, Fekete-Szücs E, Harting R, et al. Secondary metabolites of hülle cells mediate protection of fungal reproductive and overwintering structures against fungivorous animals. Elife. 2021;10. doi:10.7554/ELIFE.68058

27. Sarikaya-Bayram Ö, Bayram Ö, Feussner K, Kim J-H, Kim H-S, Kaever A, et al. Membrane-Bound Methyltransferase Complex VapA-VipC-VapB Guides Epigenetic Control of Fungal Development. Dev Cell. 2014;29: 406–420. doi:10.1016/j.devcel.2014.03.020

28. Palmer JM, Theisen JM, Duran RM, Grayburn WS, Calvo AM, Keller NP. Secondary Metabolism and Development Is Mediated by LlmF Control of VeA Subcellular Localization in Aspergillus nidulans. Heitman J, editor. PLoS Genet. 2013;9: e1003193. doi:10.1371/journal.pgen.1003193

29. Meister C, Thieme KG, Thieme S, Köhler AM, Schmitt K, Valerius O, et al. COP9 Signalosome interaction with UspA/Usp15 deubiquitinase controls VeA-mediated fungal multicellular development. Biomolecules. 2019;9: 238. doi:10.3390/biom9060238

30. Grigoriev I V, Nikitin R, Haridas S, Kuo A, Ohm R, Otillar R, et al. MycoCosm portal: gearing up for 1000 fungal genomes. Nucleic Acids Res. 2014;42: D699–704. doi:10.1093/nar/gkt1183

31. Ota M, Gonja H, Koike R, Fukuchi S. Multiple-localization and hub proteins. PLoS One. 2016;11: e0156455. doi: 10.1371/journal.pone.0156455. doi:10.1371/journal.pone.0156455

32. Haynes C, Oldfield CJ, Ji F, Klitgord N, Cusick ME, Radivojac P, et al. Intrinsic disorder is a common feature of hub proteins from four eukaryotic interactomes. PLoS Comput Biol. 2006;2: e100. doi: 10.1371/journal.pcbi.0020100. doi:10.1371/journal.pcbi.0020100

33. Patil A, Kinoshita K, Nakamura H. Hub promiscuity in protein-protein interaction networks. Int J Mol Sci. 2010;11: 1930–1943. doi:10.3390/ijms11041930

34. Kosugi S, Hasebe M, Tomita M, Yanagawa H. Systematic identification of cell cycle-dependent yeast nucleocytoplasmic shuttling proteins by prediction of composite motifs. Proceedings of the National Academy of Sciences. 2009;106: 10171–10176. doi:10.1073/pnas.0900604106

35. Rodríguez-Urra AB, Jiménez C, Nieto MI, Rodríguez J, Hayashi H, Ugalde U. Signaling the induction of sporulation involves the interaction of two secondary metabolites in *Aspergillus nidulans*. ACS Chem Biol. 2012;7: 599–606. doi:10.1021/cb200455u

36. Stark TL, Liberles DA, Holland BR, O’Reilly MM. Analysis of a mechanistic Markov model for gene duplicates evolving under subfunctionalization. BMC Evol Biol. 2017;17: 1–16. doi:10.1186/s12862-016-0848-0

37. Liberles DA, Kolesov G, Dittmar K. Understanding Gene Duplication Through Biochemistry and Population Genetics. Evolution after Gene Duplication. Hoboken, NJ, USA: John Wiley & Sons, Inc.; 2011. pp. 1–21. doi:10.1002/9780470619902.ch1

38. Bayram Ö, Braus GH, Fischer R, Rodriguez-Romero J. Spotlight on Aspergillus nidulans photosensory systems. Fungal Genet Biol. 2010;47: 900–8. doi:10.1016/j.fgb.2010.05.008

39. Bayram Ö, Feussner K, Dumkow M, Herrfurth C, Feussner I, Braus GH. Changes of global gene expression and secondary metabolite accumulation during light-dependent Aspergillus nidulans development. Fungal Genet Biol. 2016;87: 30–53. doi:10.1016/j.fgb.2016.01.004

40. Oiartzabal-Arano E, Perez-de-Nanclares-Arregi E, Espeso EA, Etxebeste O. Apical control of conidiation in Aspergillus nidulans. Curr Genet. 2016;62: 371–377. doi:10.1007/s00294-015-0556-0

41. Wu M-Y, Mead ME, Lee M-K, Neuhaus GF, Adpressa DA, Martien JI, et al. Transcriptomic, protein-DNA interaction, and metabolomic studies of VosA, VelB, and WetA in *Aspergillus nidulans* asexual spores. bioRxiv. 2020; 2020.09.09.290809. doi:10.1101/2020.09.09.290809

42. Inoue N, Wakana D, Takeda H, Yaguchi T, Hosoe T. Production of an emericellin and its analogues as fungal biological responses for Shimbu-to extract. J Nat Med. 2018;72: 357–363. doi:10.1007/s11418-017-1156-8

43. Sanchez JF, Entwistle R, Hung J-H, Yaegashi J, Jain S, Chiang Y-M, et al. Genome-based deletion analysis reveals the prenyl xanthone biosynthesis pathway in Aspergillus nidulans. J Am Chem Soc. 2011;133: 4010–7. doi:10.1021/ja1096682

44. Sanchez JF, Entwistle R, Corcoran D, Oakley BR, Wang CCC. Identification and molecular genetic analysis of the cichorine gene cluster in Aspergillus nidulans. Medchemcomm. 2012;3: 997–1002. doi:10.1039/C2MD20055D

45. Lin H, Lyu H, Zhou S, Yu J, Keller NP, Chen L, et al. Deletion of a global regulator LaeB leads to the discovery of novel polyketides in *Aspergillus nidulans*. Org Biomol Chem. 2018;16: 4973–4976. doi:10.1039/C8OB01326H

46. Pfannenstiel BT, Zhao X, Wortman J, Wiemann P, Throckmorton K, Spraker JE, et al. Revitalization of a forward genetic screen identifies three new regulators of fungal secondary metabolism in the genus Aspergillus. mBio. 2017;8: e01246–17. doi:10.1128/mBio.01246-17

47. Köhler AM, Harting R, Langeneckert AE, Valerius O, Gerke J, Meister C, et al. Integration of fungus-specific CandA-C1 into a trimeric CandA complex allowed splitting of the gene for the conserved receptor exchange factor of CullinA E3 ubiquitin ligases in Aspergilli. mBio. 2019;10. doi:10.1128/mBio.01094-19

48. Lee MK, Son YE, Park HS, Alshannaq A, Han KH, Yu JH. Velvet activated McrA plays a key role in cellular and metabolic development in Aspergillus nidulans. Sci Rep. 2020;10. doi:10.1038/s41598-020-72224-y

49. Normile D. Spoiling for a Fight With Mold. Science (1979). 2010;327: 807–807. doi:10.1126/science.327.5967.807

50. Meyer V, Andersen MR, Brakhage AA, Braus GH, Caddick MX, Cairns TC, et al. Current challenges of research on filamentous fungi in relation to human welfare and a sustainable bio-economy: a white paper Fungal Biology and Biotechnology. Fungal Biol Biotechnol. 2016;3: 6. doi:10.1186/s40694-016-0024-8

51. Kim M-J, Lee M-K, Pham HQ, Gu MJ, Zhu B, Son S-H, et al. The velvet Regulator VosA Governs Survival and Secondary Metabolism of Sexual Spores in Aspergillus nidulans. Genes (Basel). 2020;11: 103. doi:10.3390/genes11010103

52. Krappmann S, Bignell EM, Reichard U, Rogers T, Haynes K, Braus GH. The *Aspergillus fumigatus* transcriptional activator CpcA contributes significantly to the virulence of this fungal pathogen. Mol Microbiol. 2004;52: 785–799. doi:10.1111/j.1365-2958.2004.04015.x

53. Timpner C, Braus-Stromeyer SA, Tran VT, Braus GH. The Cpc1 Regulator of the Cross-Pathway Control of Amino Acid Biosynthesis Is Required for Pathogenicity of the Vascular Pathogen *Verticillium longisporum*. Molecular Plant-Microbe Interactions. 2013;26: 1312–1324. doi:10.1094/MPMI-06-13-0181-R

54. Jöhnk B, Bayram Ö, Abelmann A, Heinekamp T, Mattern DJ, Brakhage AA, et al. SCF ubiquitin ligase F-box protein Fbx15 controls nuclear co-repressor localization, stress response and virulence of the human pathogen *Aspergillus fumigatus*. Lin X, editor. PLoS Pathog. 2016;12: e1005899. doi:10.1371/journal.ppat.1005899

55. van der Lee R, Lang B, Kruse K, Gsponer J, de Groot NS, Huynen MA, et al. Intrinsically disordered segments affect protein half-life in the cell and during evolution. Cell Rep. 2014;8: 1832–1844. doi:10.1016/j.celrep.2014.07.055

56. da Fonseca PCA, He J, Morris EP. Molecular model of the human 26S proteasome. Mol Cell. 2012;46: 54–66. doi:10.1016/j.molcel.2012.03.026

57. Ciechanover A, Orian A, Schwartz AL. Ubiquitin-mediated proteolysis: biological regulation via destruction. BioEssays. 2000;22: 442–451. doi:10.1002/(SICI)1521-1878(200005)22:5<442::AID-BIES6>3.0.CO;2-Q

58. Ravid T, Hochstrasser M. Diversity of degradation signals in the ubiquitin– proteasome system. Nat Rev Mol Cell Biol. 2008;9: 679–689. doi:10.1038/nrm2468

59. von Zeska Kress MR, Harting R, Bayram Ö, Christmann M, Irmer H, Valerius O, et al. The COP9 signalosome counteracts the accumulation of cullin SCF ubiquitin E3 RING ligases during fungal development. Mol Microbiol. 2012;83: 1162–1177. doi:10.1111/j.1365-2958.2012.07999.x

60. Käfer E. Meiotic and Mitotic Recombination in Aspergillus and Its Chromosomal Aberrations. Adv Genet. 1977;19: 33–131. doi:10.1016/S0065-2660(08)60245-X

61. Hartmann T, Dümig M, Jaber BM, Szewczyk E, Olbermann P, Morschhäuser J, et al. Validation of a self-excising marker in the human pathogen Aspergillus fumigatus by employing the beta-rec/six site-specific recombination system. Appl Environ Microbiol. 2010;76: 6313–7. doi:10.1128/AEM.00882-10

62. Punt PJ, van den Hondel CAMJJ. Transformation of filamentous fungi based on hygromycin b and phleomycin resistance markers. Methods Enzymol. 1992. pp. 447–457. doi:10.1016/0076-6879(92)16041-H

63. Bui T, Harting R, Braus-Stromeyer SA, Tran V, Leonard M, Höfer A, et al. *Verticillium dahliae* transcription factors Som1 and Vta3 control microsclerotia formation and sequential steps of plant root penetration and colonisation to induce disease. New Phytologist. 2019;221: 2138–2159. doi:10.1111/nph.15514

64. Inoue H, Nojima H, Okayama H. High efficiency transformation of Escherichia coli with plasmids. Gene. 1990;96. doi:10.1016/0378-1119(90)90336-P

65. Hanahan D, Jessee J, Bloom FR. Plasmid transformation of Escherichia coli and other bacteria. Methods Enzymol. 1991;204: 63–113.

66. Starke J, Harting R, Maurus I, Leonard M, Bremenkamp R, Heimel K, et al. Unfolded Protein Response and Scaffold Independent Pheromone MAP Kinase Signaling Control Verticillium dahliae Growth, Development, and Plant Pathogenesis. J Fungi (Basel). 2021;7. doi:10.3390/JOF7040305

67. Kang D, Gho YS, Suh M, Kang C. Highly sensitive and fast protein detection with Coomassie Brilliant Blue in sodium dodecyl sulfate-polyacrylamide gel electrophoresis. Communications. 2002;23: 1511–1512. doi:10.5012/bkcs.2002.23.11.1511

68. Shevchenko A, Jensen ON, Podtelejnikov AV, Sagliocco F, Wilm M, Vorm O, et al. Linking genome and proteome by mass spectrometry: Large-scale identification of yeast proteins from two dimensional gels. Biochemistry. 1996;93: 14440–14445.

69. Opitz N, Schmitt K, Hofer-Pretz V, Neumann B, Krebber H, Braus GH, et al. Capturing the Asc1p/Receptor for Activated C Kinase 1 (RACK1) Microenvironment at the Head Region of the 40S Ribosome with Quantitative BioID in Yeast. Mol Cell Proteomics. 2017;16: 2199–2218. doi:10.1074/mcp.M116.066654

70. Schmitt K, Smolinski N, Neumann P, Schmaul S, Hofer-Pretz V, Braus GH, et al. Asc1p/RACK1 Connects Ribosomes to Eukaryotic Phosphosignaling. Mol Cell Biol. 2017;37: e00279–16. doi:10.1128/MCB.00279-16

71. Ward JJ, McGuffin LJ, Bryson K, Buxton BF, Jones DT. The DISOPRED server for the prediction of protein disorder. Bioinformatics. 2004;20: 2138–2139. doi:10.1093/bioinformatics/bth195

72. Alvarez-Jarreta J, Amos B, Aurrecoechea C, Bah S, Barba M, Barreto A, et al. VEuPathDB: the eukaryotic pathogen, vector and host bioinformatics resource center in 2023. Nucleic Acids Res. 2024;52: D808–D816. doi:10.1093/NAR/GKAD1003

73. Sayers EW, Bolton EE, Brister JR, Canese K, Chan J, Comeau DC, et al. Database resources of the national center for biotechnology information. Nucleic Acids Res. 2022;50: D20–D26. doi:10.1093/NAR/GKAB1112

74. Harrison PW, Amode MR, Austine-Orimoloye O, Azov AG, Barba M, Barnes I, et al. Ensembl 2024. Nucleic Acids Res. 2024;52: D891–D899. doi:10.1093/nar/gkad1049

