## Supplemental Material for "The VelB intrinsically disordered domain promotes selective heterodimer formation of velvet domain regulatory proteins for fungal development"

### Supplementary Figures and Tables

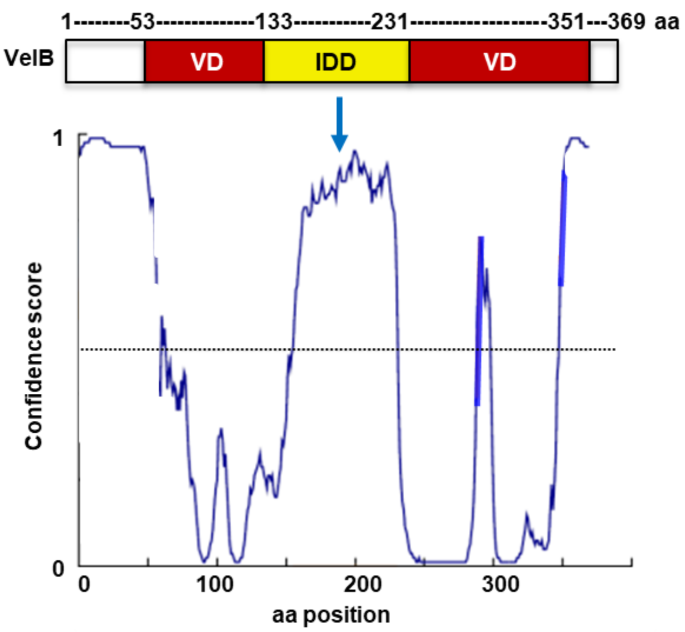

**Fig S1: *A. nidulans* VelB is interrupted by a 99 amino acid intrinsically disordered domain (IDD).** Structure prediction of *A. nidulans* VelB using the DISOPRED program [1]. The velvet domain (VD: in red) for DNA binding and dimerization is interrupted by an intrinsically disordered domain (IDD, yellow) of 99 amino acids. The dotted line represents the cut-off of 0.5 of the confidence score (disorder probability).

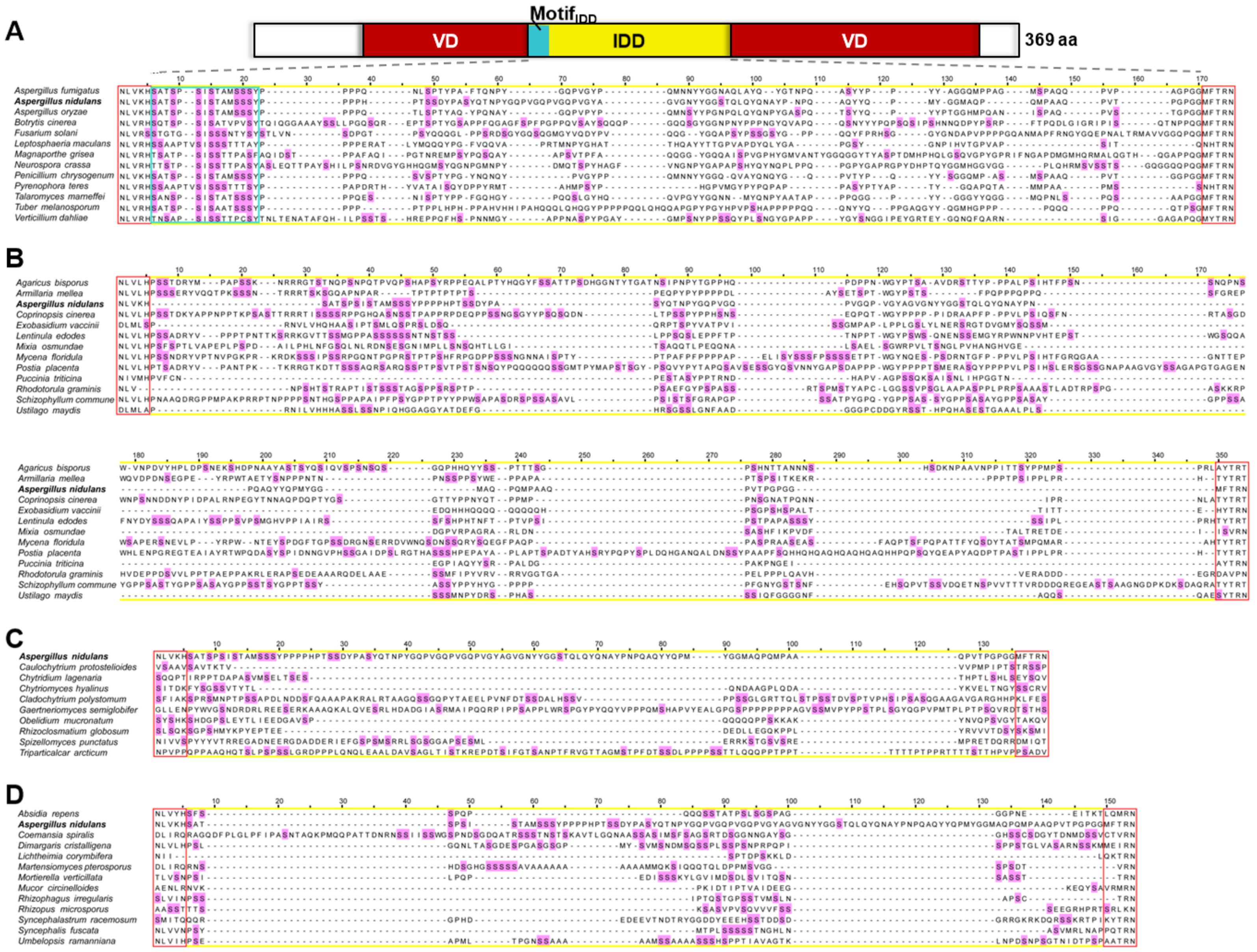
A
Motif IDD
VD
IDD
VD
369 aa
Aspergillus fumigatus
Aspergillus nidulans
Aspergillus oryzae
Botrytis cinerea
Fusarium solani
Leptosphaeria maculans
Magnaporthe grisea
Neurospora crassa
Penicillium chrysogenum
Pyrenophora teres
Talaromyces marneffei
Tuber melanosporum
Verticillium dahliae
B
Agaricus bisporus
Armillaria mellea
Aspergillus nidulans
Coprinopsis cinerea
Exobasidium vaccinii
Lentinula edodes
Mixia osmundae
Mycena floridula
Postia placenta
Puccinia triticina
Rhodotorula graminis
Schizophyllum commune
Ustilago maydis
C
Aspergillus nidulans
Caulochytrium protostelioides
Chytridium lagenaria
Chytriomyces hyalinus
Cladochytrium polystomum
Gaertneriomyces semiglobifer
Obelidium mucronatum
Rhizoclosmatium globosum
Spizellomyces punctatus
Triparticalcar arcticum
D
Absidia repens
Aspergillus nidulans
Coemansia spiralis
Dimargaris cristalligena
Lichtheimia corymbifera
Martensiomyces pterosporus
Mortierella verticillata
Mucor circinelloides
Rhizophagus irregularis
Rhizopus microsporus
Syncephalastrum racemosum
Syncephalis fuscata
Umbelopsis ramanniana

**Fig S2: Within the fungal kingdom VelB IDDs vary in length and amino acid sequence.** Alignments of deduced protein sequences for the *A. nidulans* VelB intrinsically disordered domain (IDD) (yellow box) against VelB proteins of indicated fungal species as examples for the **(A)** Ascomycota, **(B)** Basidiomycota, **(C)** Zygomycota or **(D)** Chytridiomycota. Serine residues as putative phosphorylation sites are highlighted in pink. The VelB IDDs of Ascomycota contain the conserved $\text{Motif}_{IDD}$ (turquoise box). The N-terminal boundary of the VelB velvet domain (VD) is conserved in all analyzed fungal genomes (red box).

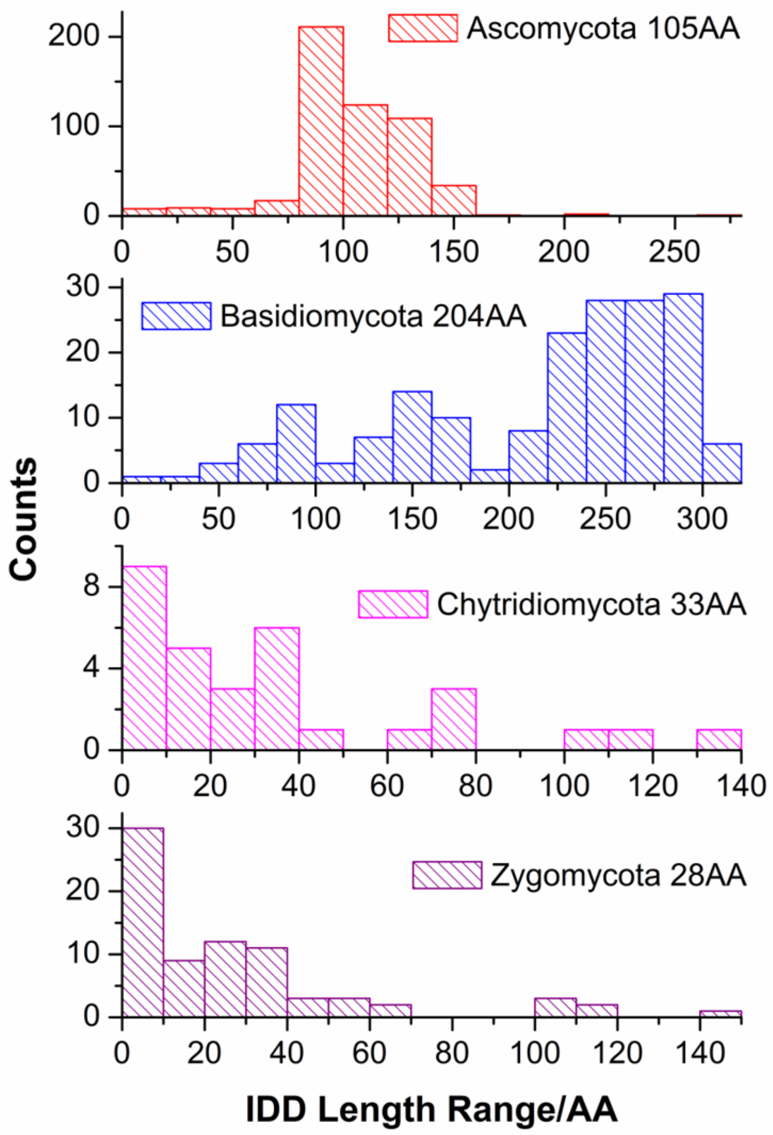

**Fig S3: VelB IDD variations between different fungal divisions.** X-axis represents IDD lengths in amino acid residues and y-axis the number of fungal species within the four divisions (corresponding orthologs and deduced IDD information in Table S1 and S2).

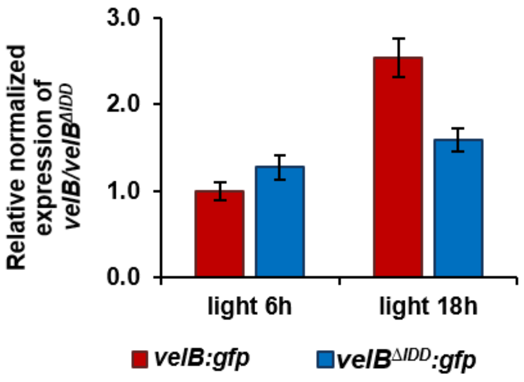

**Fig S4: Relative expression of *velB:gfp* transcripts is similar with or without IDD.** Strains were grown in submerged cultures for 24h, shifted to solid minimal medium and incubated for 6 or 18h in the light at 37°C. qRT-PCR analyses indicate that transcription of the truncated *velB*$^{\Delta IDD}$*:gfp* gene is similar to the full length *velB:gfp* gene. Data from two biological and three technical replicates are shown. *15S rRNA* and *h2A* were used as references for normalization.

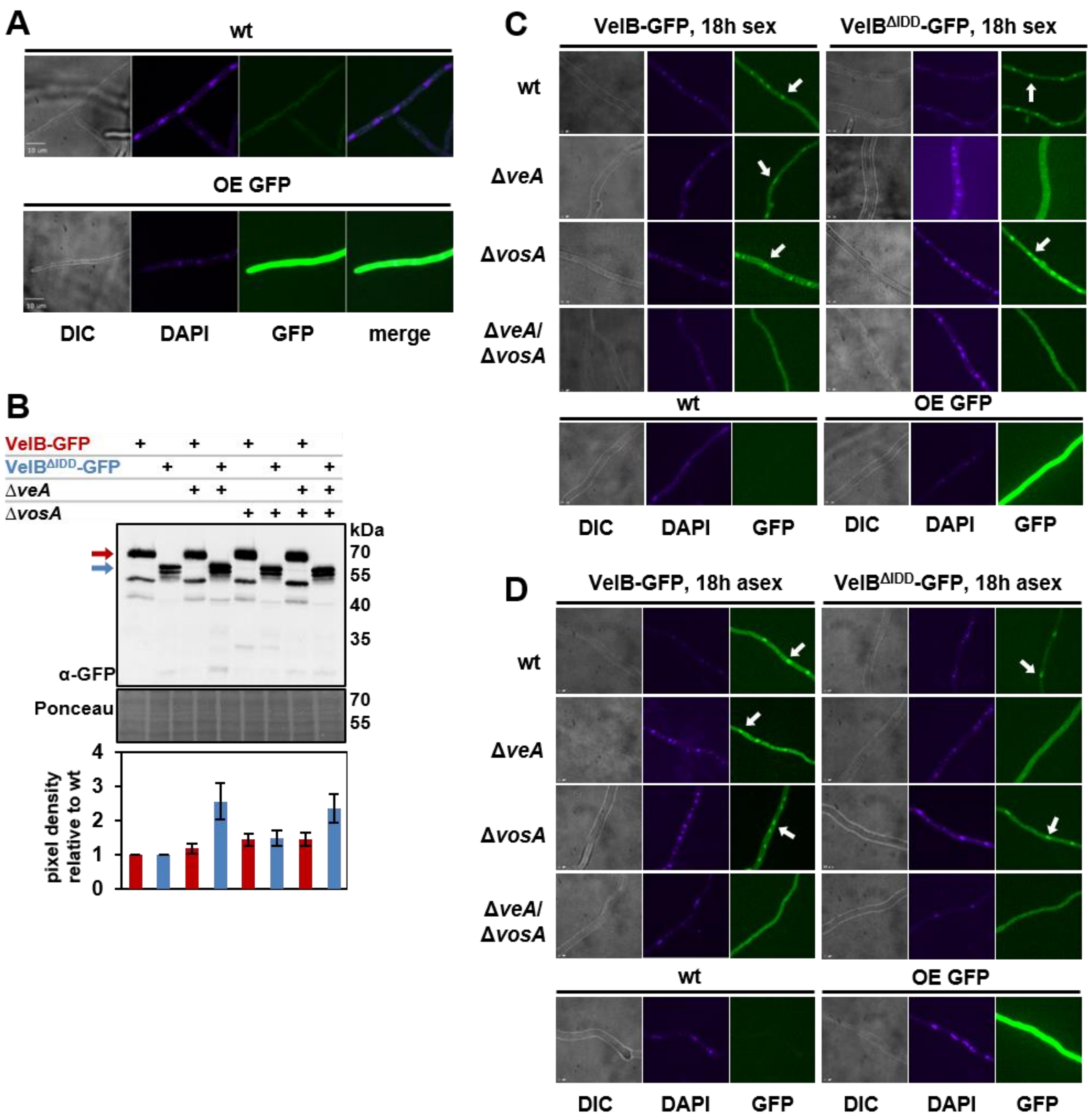

**Fig S5: VelB IDD is required for selective heterodimer formation with VosA and subsequent nuclear localization. (A)** Fluorescence micrographs of 18h vegetatively grown wild type (wt) and constitutively expressing GFP (OE GFP) strains as control for microscopy. **(B)** Western blot experiment for VelB-GFP and VelB$^{\Delta IDD}$-GFP in wild type (wt), *veA* (Δ*veA*) and *vosA* (Δ*vosA*) single and double deletion background used in microscopy. The crude extracts were obtained from mycelium of vegetatively grown cultures (18h). The pixel density was calculated using the BioID software (Vilber Lourmat), normalized against the loading control Ponceau and calculated relative to VelB-GFP (red bars) or VelB$^{\Delta IDD}$-GFP (blue bars) in wild type background. **(C+D)** Fluorescence microscopy of the same strains after growth in 18h sexual **(C)** and 18h asexual **(D)** development inducing conditions. White arrows indicate nuclei. Size bar represents 10 µm.

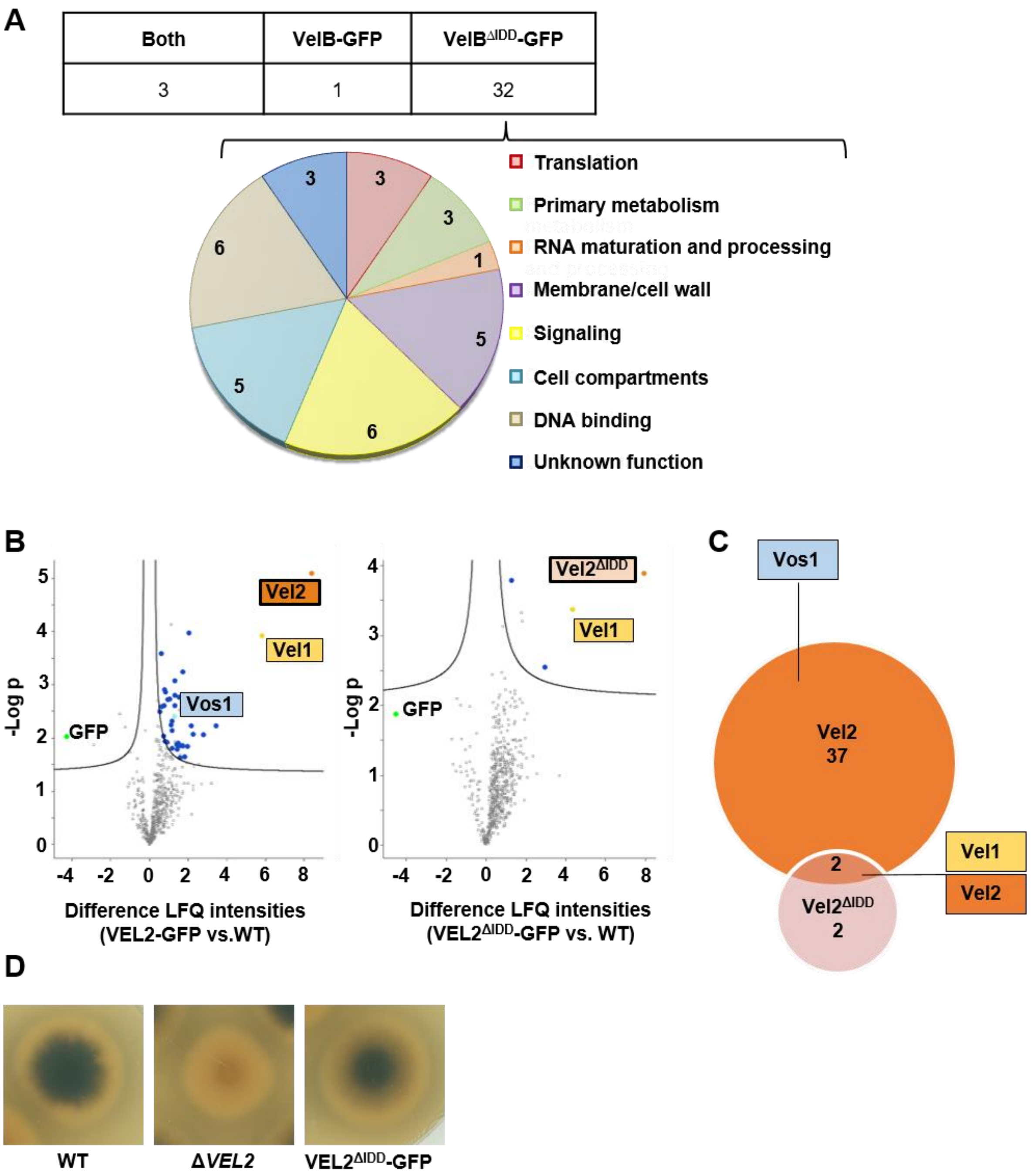

**Fig S6: The VelB IDD allows selective VelB-VosA formation in *A. nidulans* and *V. dahliae*.**

**(A)** GFP pull downs of VelB-GFP and VelB$^{\Delta IDD}$-GFP from vegetatively grown *A. nidulans* cultures followed by LC-MS/MS revealed that the lack of the IDD leads to an increased number of putative interaction partners (Table S3). The three proteins found in both GFP pulldowns (left column) are the corresponding bait protein itself, the catalase CatB and the velvet protein VeA. The single protein, which is only recruited by VelB-GFP (middle column) is the velvet protein VosA. The diagram depicts the predicted cellular function or localization of the 32 putative interaction partners of VelB$^{\Delta IDD}$-GFP. Numbers represent proteins identified in at least two out of three biological repetitions with MS/MS counts ≥ 4, unique peptides ≥ 3 and LFQ intensity ≥ 20 and which were absent in the control strain. **(B)** *V. dahliae* Vel2-GFP or Vel2$^{\Delta IDD}$-GFP strains were used for GFP pull down experiments, followed by LC-MS analysis. The x-axis are fold-differences of LFQ values of the strains indicated (mean of three independent experiments). The y-axis shows the -Log p value resulting from a two-tailed t-test. Significantly interacting proteins are found in the upper right part of the plot (Table S4). Missing values were replaced four times with imputed values and proteins, which showed a significant interaction in all 4 repetitions, are colored (Vel1: yellow, Vel2 respectively Vel2$^{\Delta IDD}$: orange, Vos1: turquois, other proteins: dark blue, GFP: lime). **(C)** Comparison of significantly enriched proteins in Vel2 and Vel2$^{\Delta IDD}$ pull downs. Among the 39 significantly enriched proteins in Vel2 and four enriched proteins in Vel2$^{\Delta IDD}$ are two overlaps, Vel1 and Vel2. Within the proteins enriched in Vel2 is also Vos1. Vos1 was not significantly enriched in the Vel2$^{\Delta IDD}$ strain.

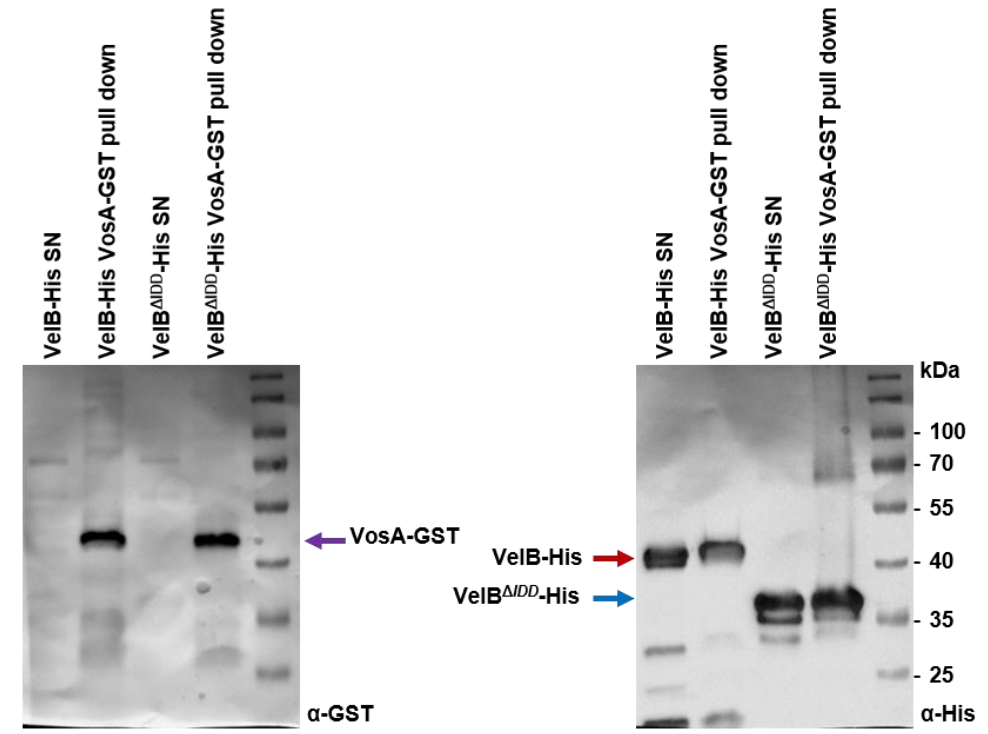

**Fig S7: *In vitro* VelB binds full length VosA independently of the IDD*.*** *In vitro* co-immunoprecipitation of VosA-GST and VelB-His or VelB$^{\Delta IDD}$-His fusion proteins recombinantly expressed in *E. coli.* Specific detection of fusion proteins was achieved with α-His or α-GST antibodies. VosA-GST (violet arrow) pulls VelB-His (red arrow) and VelB$^{\Delta IDD}$-His (blue arrow) fusion proteins. SN = supernatant of lysed cells after centrifugation.

A

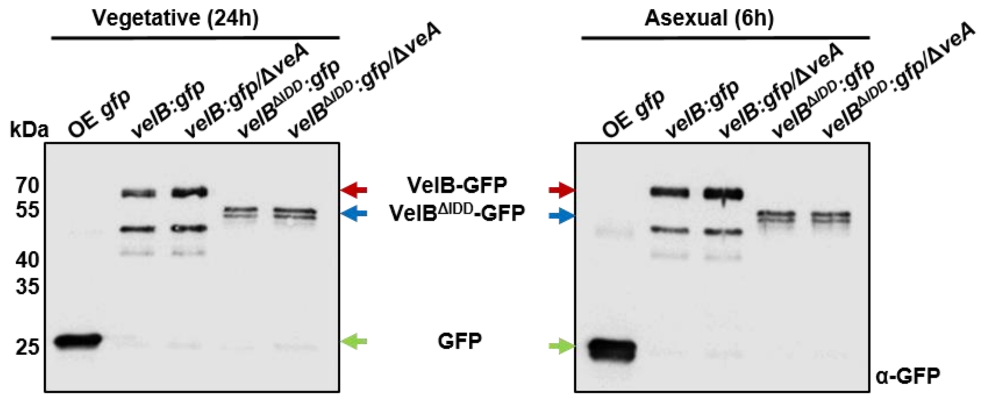

B

| | *velB:gfp* | *velB:gfp/ΔveA* | *velB$^{\Delta IDD}$:gfp* | *velB$^{\Delta IDD}$:gfp/ΔveA* |
|---|---|---|---|---|
| VelB | + | + | + | + |
| VeA | + | - | + | - |
| VosA | + | + | - (*) | + |

**Fig S8: Heterodimer formation of VosA with VelB missing IDD is restored in absence of VeA. (A)** Western experiment of 24h vegetative and six hour light incubated *A. nidulans* strains shows GFP signals of *velB:gfp* and *velB$^{\Delta IDD}$:gfp* in wild type and Δ*veA* background. OE = GFP over expression. **(B)** Summary of identified velvet proteins VelB, VeA and VosA in each GFP pull down (*velB:gfp, velB:gfp/ΔveA, velB$^{\Delta IDD}$:gfp* and *velB$^{\Delta IDD}$:gfp/ΔveA*). Plus (+) means identified as interaction partner in at least two replicates, minus (-) means not identified or only with MS/MS or unique peptide counts lower than one (Table S5).

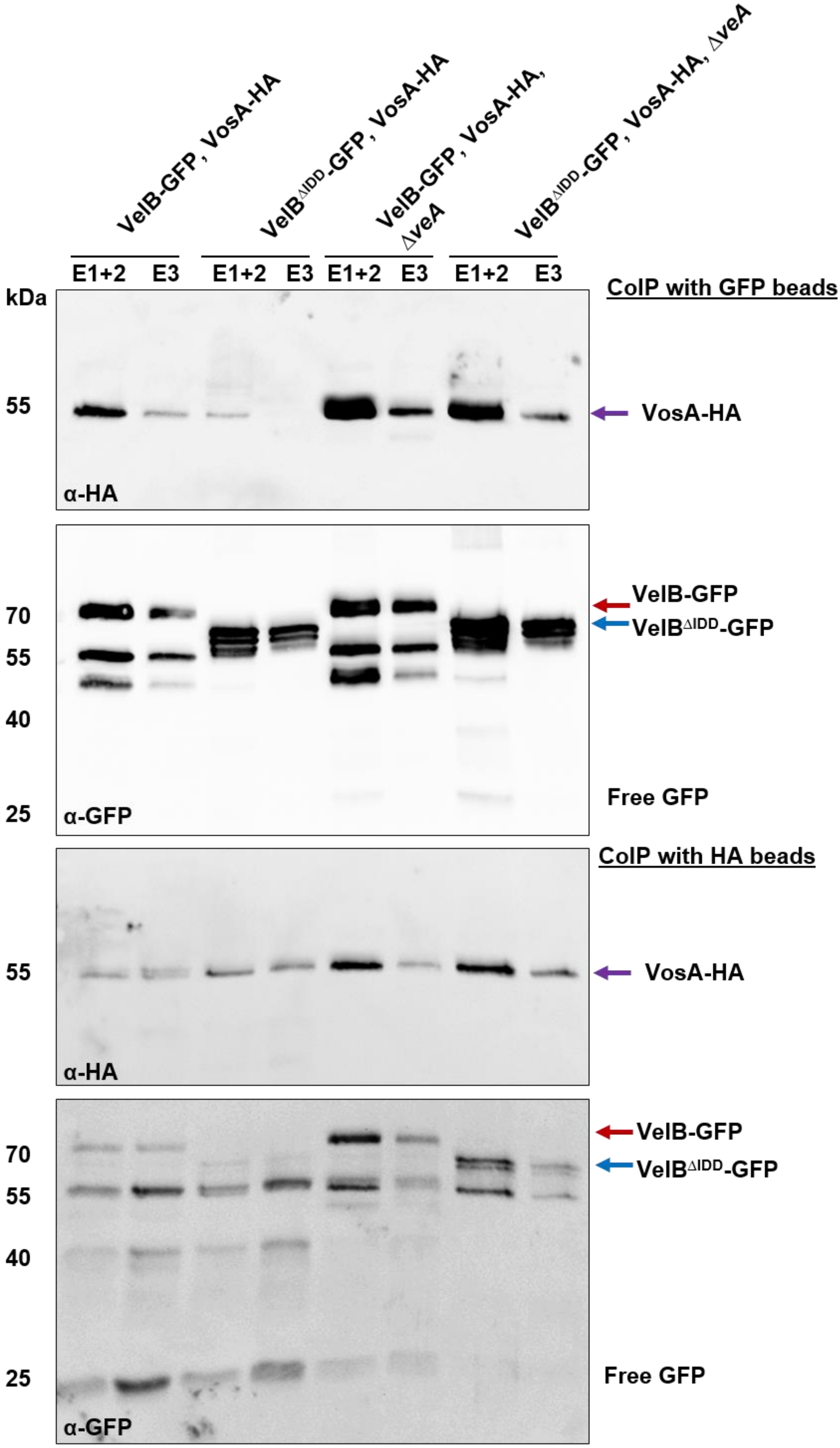
VelB-GFP, VosA-HA
VelB$^{\Delta IDD}$-GFP, VosA-HA
VelB-GFP, VosA-HA, ΔveA
VelB$^{\Delta IDD}$-GFP, VosA-HA, ΔveA
E1+2
E3
kDa
CoIP with GFP beads
55
VosA-HA
α-HA
70
55
40
25
VelB-GFP
VelB$^{\Delta IDD}$-GFP
Free GFP
α-GFP
CoIP with HA beads
55
VosA-HA
α-HA
70
55
40
25
VelB-GFP
VelB$^{\Delta IDD}$-GFP
Free GFP
α-GFP

**Fig S9: VelB-VosA heterodimer formation is reduced in absence of the IDD and *veA.*** Western experiment of CoIP samples from 20h vegetative mycelium of indicated strains. Co-IPs were conducted with GFP and HA beads and the elution fractions (E) were subjected to SDS PAGE following western experiments using anti-GFP and anti-HA antibodies.

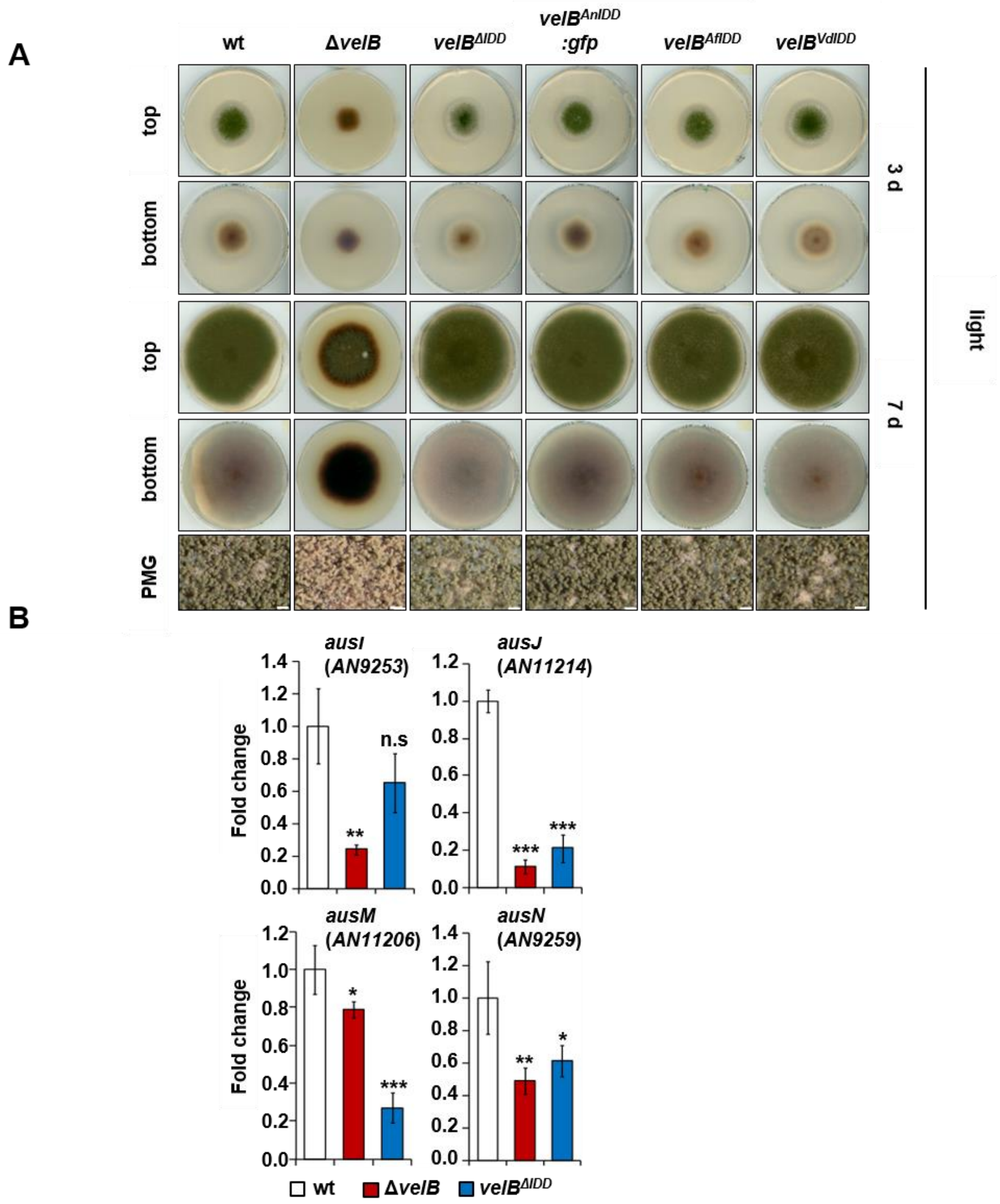

**Fig S10: Phenotypes of *A. nidulans veA* or *vosA* deletion strains are epistatic to *velB*$^{\Delta IDD}$ in corresponding double mutant strains. (A)** Phenotypes of wildtype (wt), *velB* deletion (Δ*velB*), *velB*$^{IDD}$ deletion (*velB*$^{\Delta IDD}$) and complementation strains with the IDD of *A. nidulans, A. fumigatus* or *Verticillium dahliae* into *velB*$^{\Delta IDD}$ (*velB*$^{AnIDD}$*:sgfp, velB*$^{AfIDD}$ and *velB*$^{VdIDD}$) on solid minimal medium. 2000 spores were spotted in the middle of the plate and grown in light for 3 or 7 days (d) at 37°C. PMG photomicrograph, scale bar = 100 µm. **(B)** Differential expression of genes, essential for the synthesis of austinol and dehydroaustinol, in Δ*velB* and *velB*$^{\Delta IDD}$

strains. mRNAs were extracted from mycelia grown in submerged cultures shaking at 37°C for 20h in light. For normalization of gene expression the reference gene *h2A* was used and the expression of all samples was calibrated towards the wt. Data presented are the averages and standard deviations derived by at least three biological replicates, each with at least three technical replicates. Statistics were performed by Student's t test: * $p < 0.05$, ** $p < 0.01$ and *** $p < 0.001$, n.s.: not significant.

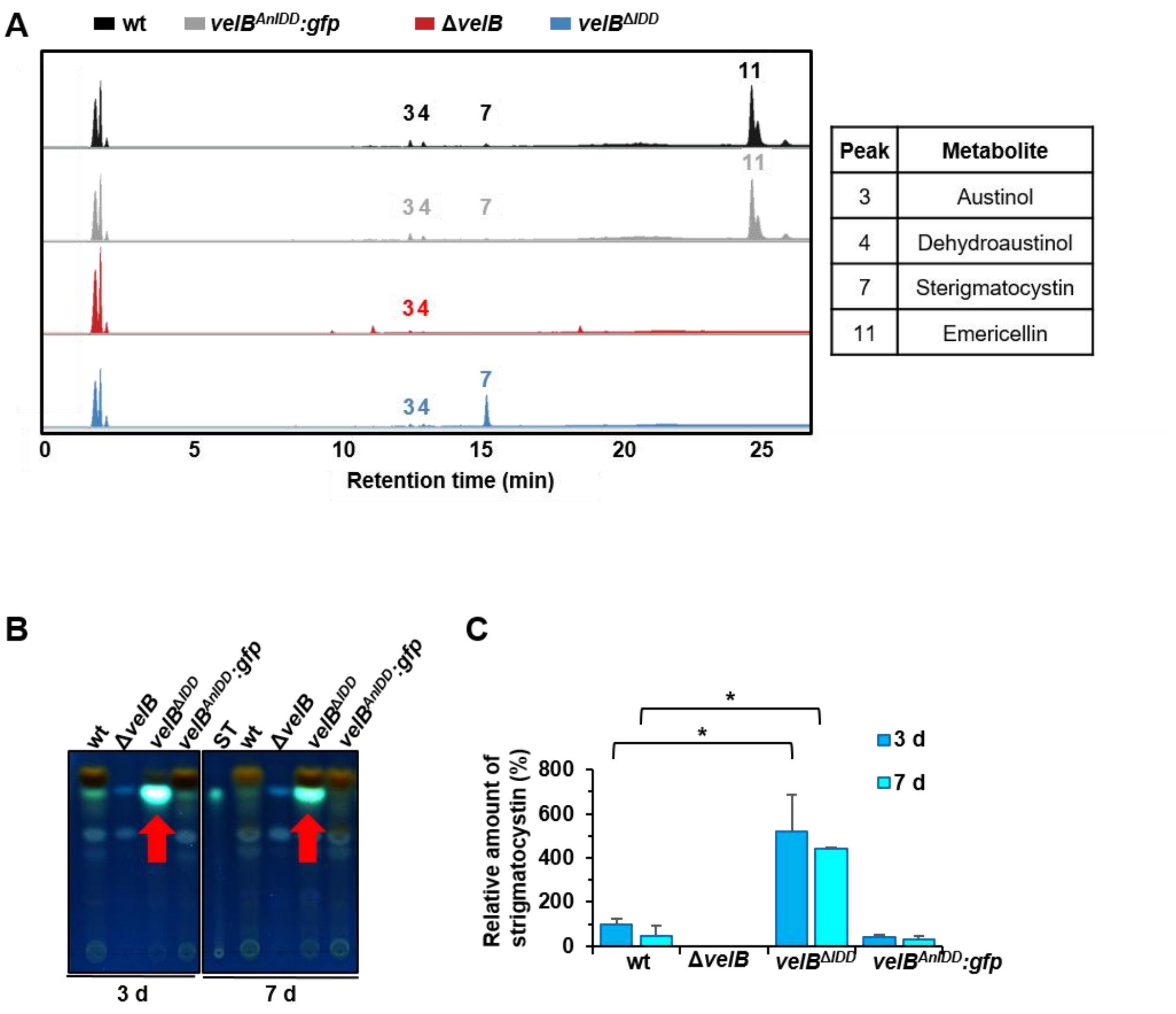

**Fig S11: Sterigmatocystin production depends on the VelB IDD. (A)** Secondary metabolite production is altered in the *velB*$^{\Delta IDD}$ strain. High performance liquid chromatography coupled with mass spectrometry (HPLC-MS) was performed with metabolites extracted after 3 days of growth in light. Metabolites were isolated of wildtype (wt), velB deletion (Δ*velB*), *velB*$^{IDD}$ deletion (*velB*$^{\Delta IDD}$) and *velB*$^{\Delta IDD}$ complementation (*velB*$^{AnIDD}$*:gfp*). The chromatogram shows identified secondary metabolites at 250 nm, indicated with numbers (peak 3: austinol, peak 4: dehydroaustinol, peak 7: sterigmatocystin, peak 11: emericellin) and listed in table S6. **(B)** Thin layer chromatography (TLC) shows increased sterigmatocystin production in the *velB*$^{\Delta IDD}$ strains (red arrow) after 3 or 7 days of growth in the light. The signals of the TLC plate were detected at 366 nm after derivatization with $AlCl_3$. ST = Sterigmatocystin standard.

**(C)** Quantification of the sterigmatocystin signal relative to wt. Error bars indicate the SEM. Statistics were performed by Student's t test: * $p < 0.05$.

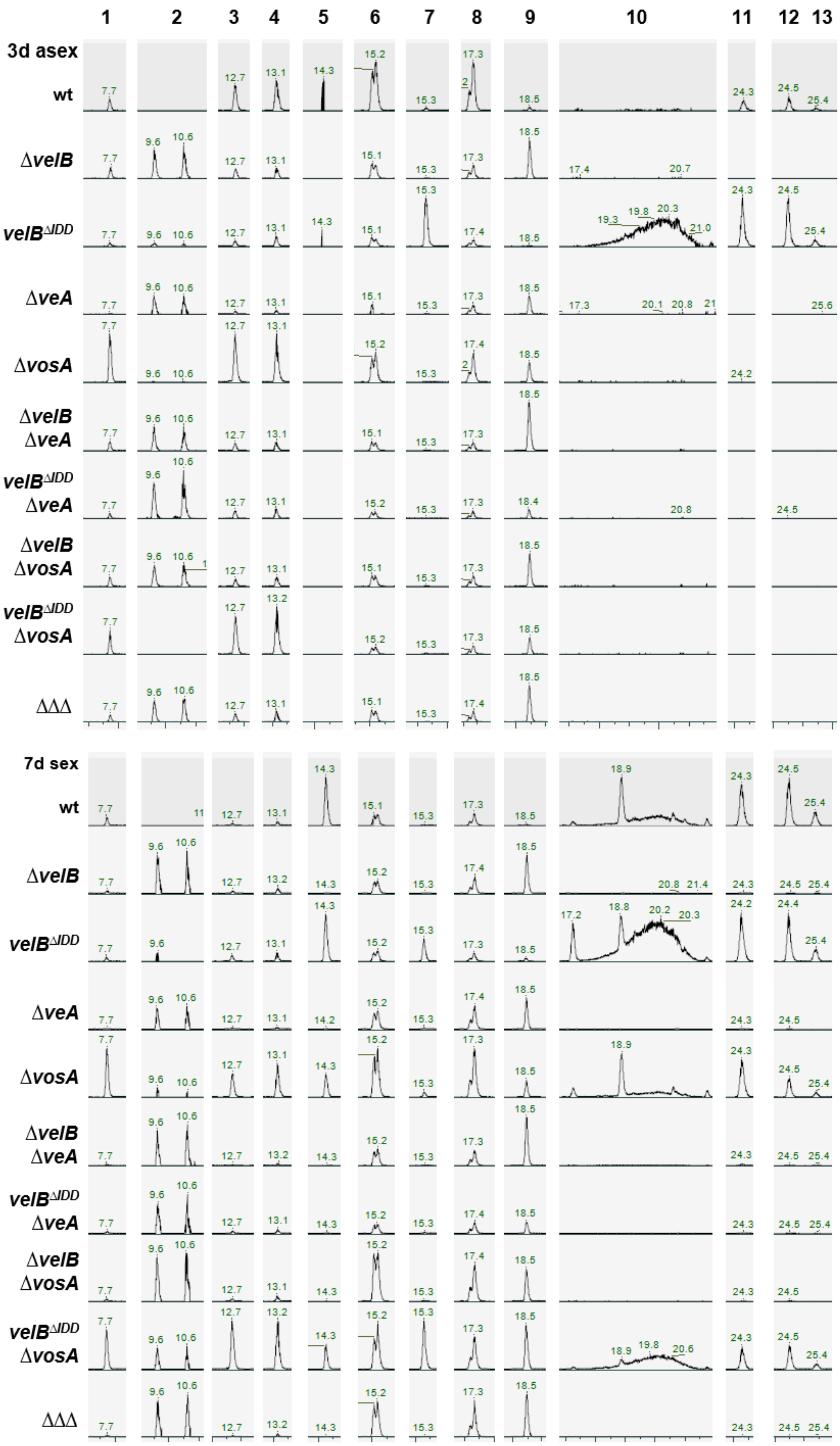
1 2 3 4 5 6 7 8 9 10 11 12 13
3d asex
wt
ΔvelB
velB^ΔIDD
ΔveA
ΔvosA
ΔvelB ΔveA
velB^ΔIDD ΔveA
ΔvelB ΔvosA
velB^ΔIDD ΔvosA
ΔΔΔ
7d sex
wt
ΔvelB
velB^ΔIDD
ΔveA
ΔvosA
ΔvelB ΔveA
velB^ΔIDD ΔveA
ΔvelB ΔvosA
velB^ΔIDD ΔvosA
ΔΔΔ

**Fig S12 Extracted ion chromatograms (EICs) of detected peaks.** The EICs show the identified metabolites (M ± 5 ppm) from table S6 measured from 3 days asexual and 7 days sexual developmental samples in positive ionization mode $[M+H]^+$. Masses are given in table S6. Cichorine (1), F-9775A/B (2), Austinol (3), Dehydroaustinol (4), Arugosin H (5), Emericellamide C (6), Sterigmatocystin (7), Emericellamide E (8), Terrequinone A (9), Arugosin A (10), Emericellin (11), Shamixanthone (12), Epishamixanthone (13). Structures are depicted in Fig S13.

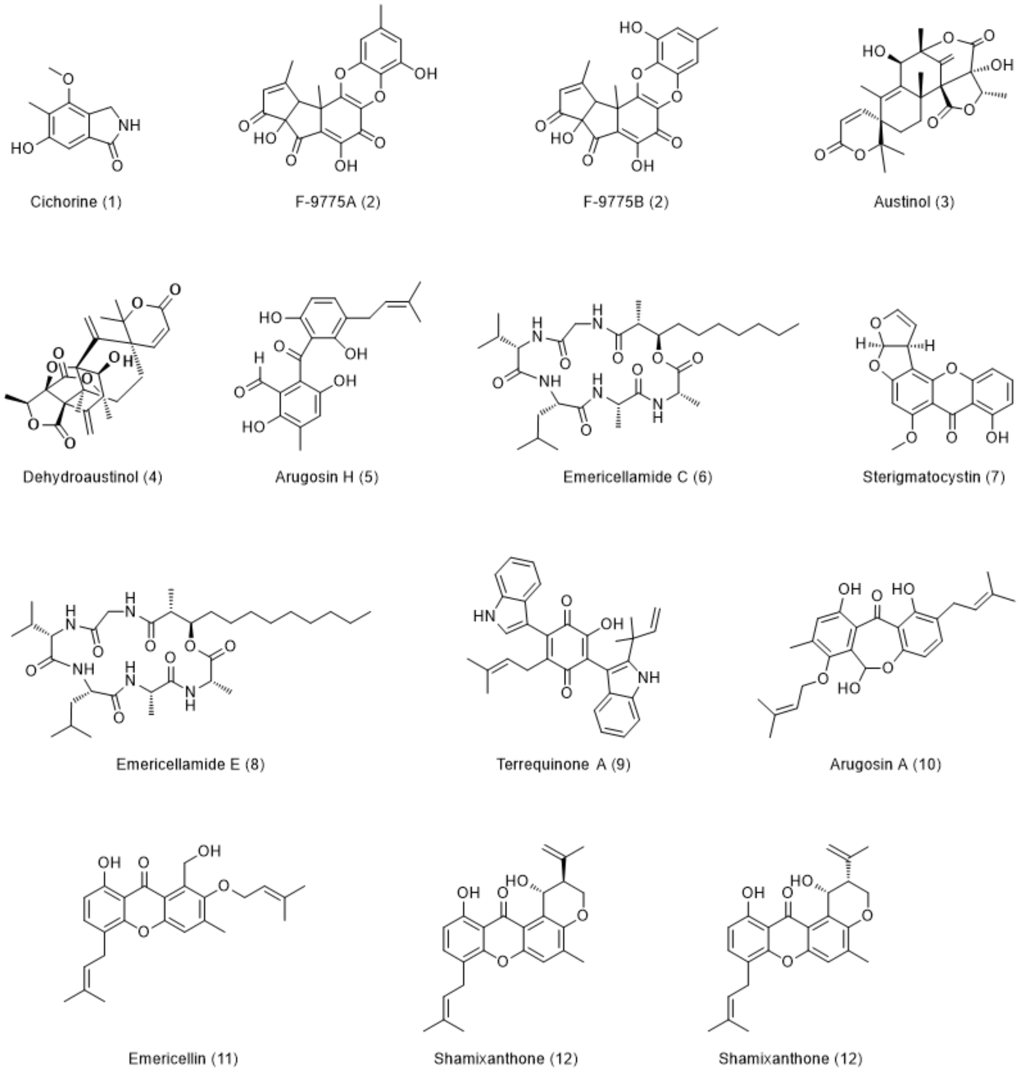

**Fig S13: Overview of identified metabolites and their structures from Table S6.**

**Table S1: Summary on IDD information (Excel file)**

**Table S2: Collection of fungal genomes with multiple VelB homologs (Excel file)**

**Table S3: Putative interaction partners identified in the GFP pull down of VelB-GFP and VelB$^{\Delta IDD}$-GFP expressing strains.**

| **Sys. name** | **Std. name** | **Description** | **VelB-GFP LFQ** | **VelB$^{\Delta IDD}$-GFP LFQ** | **VelB-GFP MS/MS** | **VelB$^{\Delta IDD}$-GFP MS/MS** | **VelB-GFP Unique peptides** | **VelB$^{\Delta IDD}$-GFP Unique peptides** |
|---|---|---|---|---|---|---|---|---|
| **AN0363** | **VelB** | **Bait protein** | 28.88<br>26.18<br>25.22 | 29.77<br>27.77<br>28.05 | 735<br>191<br>112 | 728<br>139<br>36 | 15<br>11<br>13 | 16<br>9<br>9 |
| **AN1052** | **VeA** | Protein involved in light-sensitive control of differentiation and secondary metabolism | 25.51<br>23.75<br>23.78 | 26.35<br>25.85<br>26.01 | 406<br>125<br>89 | 405<br>98<br>47 | 17<br>13<br>16 | 16<br>10<br>12 |
| **AN1959** | **VosA** | Nuclear protein involved in spore formation and trehalose accumulation | 24.32<br>22.03<br>16.89 | 14.92<br>15.86<br>15.24 | 166<br>8<br>1 | 1<br>0<br>0 | 11<br>3<br>1 | 1<br>0<br>0 |
| **AN9339** | **CatB** | Hyphal catalase with a predicted role in gluconic acid and gluconate metabolism | 23.9<br>20.69<br>23.05 | 22.7<br>22.65<br>23.39 | 168<br>8<br>30 | 125<br>40<br>46 | 15<br>4<br>15 | 16<br>10<br>17 |
| Translation | | | | | | | | |
| **AN0907** | **unchar.** | Putative 40S ribosomal protein S23 (Rps23), putative ortholog of *S. cerevisiae* RPS23A | 15.88<br>17.85<br>17.35 | 20.14<br>20.74<br>15.76 | 6<br>2<br>0 | 19<br>10<br>1 | 1<br>1<br>0 | 6<br>2<br>1 |

| | | | | | | | | |
|---|---|---|---|---|---|---|---|---|
| **AN1345** | **unchar.** | Putative 40S ribosomal protein S22 (Rps22), putative ortholog of *S. cerevisiae* RPS22A | 16.39<br>17.58<br>16.24 | 19.39<br>20.05<br>15.41 | 1<br>0<br>0 | 15<br>10<br>2 | 1<br>0<br>0 | 2<br>2<br>1 |
| **AN0314** | **unchar.** | Putative aspartyl-tRNA synthetase, putative ortholog of *S. cerevisiae* DPS1 | 15.39<br>17.09<br>17.03 | 21.86<br>22.11<br>23.06 | 4<br>0<br>0 | 159<br>58<br>81 | 1<br>0<br>0 | 23<br>10<br>24 |
| Primary metabolism | | | | | | | | |
| **AN5226** | **AcpA** | Acetate permease A, involved in acetate uptake | 15.39<br>17.33<br>16.91 | 21.01<br>19.72<br>16.67 | 1<br>0<br>0 | 39<br>4<br>2 | 1<br>0<br>0 | 4<br>2<br>1 |
| **AN9180** | **unchar.** | Putative transketolase, putative ortholog of *A. fumigatus* TktA | 16.88<br>17.1<br>15.96 | 20.42<br>19.98<br>15.46 | 0<br>0<br>0 | 52<br>14<br>1 | 0<br>0<br>0 | 14<br>6<br>1 |
| **AN1967** | **PpoA** | Psi factor producing oxygenase A, responsible for the formation of the oxylipin psiBα | 15.06<br>17.36<br>17.84 | 20.98<br>18.32<br>20.6 | 0<br>0<br>0 | 86<br>5<br>17 | 0<br>0<br>0 | 23<br>3<br>11 |
| RNA maturation and processing | | | | | | | | |
| **AN7474** | **unchar.** | Has domain(s) with predicted RNA binding, putative ortholog of *S. cerevisiae* JSN1 | 16<br>17.14<br>16.1 | 20.65<br>20.73<br>21.04 | 0<br>0<br>0 | 51<br>13<br>15 | 0<br>0<br>0 | 16<br>7<br>9 |
| Membrane/cell wall | | | | | | | | |
| **AN5020** | **ArfB/ AdpA** | ADP ribosylation factor B, required for | 15.05<br>17.41<br>16.55 | 21.33<br>21.80<br>19.1 | 0<br>0<br>0 | 49<br>19<br>4 | 0<br>0<br>0 | 4<br>3<br>3 |

| | | | | | | | | |
|---|---|---|---|---|---|---|---|---|
| | | normal endocytosis and polarized growth | | | | | | |
| **AN9079** | **unchar.** | Putative ortholog of *N. crassa* Ham-10, with role in conidia formation | 15.24<br>16.91<br>16.94 | 20<br>20.13<br>21.26 | 0<br>0<br>0 | 50<br>50<br>36 | 0<br>0<br>0 | 18<br>7<br>20 |
| **AN2928** | **unchar.** | Putative cell wall protein, uncharacterized | 15.09<br>16.25<br>16.29 | 19.08<br>20.56<br>20.17 | 0<br>0<br>1 | 11<br>7<br>4 | 0<br>0<br>1 | 3<br>2<br>2 |
| **AN11139** | **AbpA** | Putative actin-binding protein A of the cortical actin patches | 14.88<br>16.39<br>17.13 | 19.15<br>19.68<br>20.67 | 3<br>0<br>2 | 46<br>21<br>13 | 1<br>0<br>1 | 11<br>5<br>6 |
| **AN3098** | **NsfA** | Putative secretory component, ortholog of *S. cerevisiae* SEC18, has similarity to mammalian N-ethylmaleimide-sensitive factor | 13.88<br>17.07<br>16.29 | 18.5<br>19.77<br>20.64 | 2<br>0<br>0 | 16<br>20<br>32 | 1<br>0<br>0 | 8<br>7<br>16 |
| Signaling | | | | | | | | |
| **AN3102** | **PhkA** | Putative histidine-containing phosphotransfer protein | 16.59<br>17.07<br>17.68 | 19.53<br>17.61<br>20.65 | 0<br>0<br>0 | 46<br>6<br>26 | 0<br>0<br>0 | 16<br>4<br>17 |
| **AN3422** | **Ste7** | MAP kinase kinase (MAPKK), component of a signaling module SteD-SteC-MkkB-MpkB that controls coordination of development and secondary metabolism | 14.02<br>17<br>17.42 | 20.75<br>20.75<br>20.57 | 1<br>0<br>0 | 69<br>20<br>15 | 1<br>0<br>0 | 11<br>5<br>7 |

| | | | | | | | | |
|---|---|---|---|---|---|---|---|---|
| **AN2130** | **unchar.** | Putative Ras guanine-nucleotide exchange factor activity, putative ortholog of *S. cerevisiae* CDC25 | 16.26<br>17.22<br>17.95 | 19.97<br>20.83<br>21.07 | 0<br>0<br>0 | 51<br>26<br>16 | 0<br>0<br>0 | 10<br>8<br>9 |
| **AN7576** | **unchar.** | Predicted Rho GTPase activating protein, putative ortholog of *S. pombe* Rga1 | 14.98<br>16.97<br>16.95 | 19.23<br>20.01<br>19.66 | 0<br>0<br>0 | 32<br>18<br>14 | 0<br>0<br>0 | 8<br>7<br>9 |
| **AN8836** | **Cla4** | Predicted PAK (p21-activated kinase) family protein | 15.06<br>18.01<br>16.83 | 20.79<br>21.62<br>22.53 | 0<br>0<br>0 | 58<br>48<br>46 | 0<br>0<br>0 | 14<br>11<br>18 |
| **AN0463** | **unchar.** | Predicted Rac guanine nucleotide exchange factor, putative ortholog of *C. albicans* DCK2 | 14.68<br>16.46<br>17.32 | 18.9<br>20<br>21.76 | 0<br>0<br>0 | 42<br>18<br>24 | 0<br>0<br>0 | 9<br>9<br>23 |
| Cell compartments | | | | | | | | |
| **AN4207** | **unchar.** | Putative ortholog of *C. albicans* APL4, with role in endosomal transport, vesicle mediated transport and AP-1 adaptor complex | 14.12<br>17.72<br>16.76 | 19.94<br>20.71<br>21.41 | 0<br>0<br>0 | 44<br>26<br>26 | 0<br>0<br>0 | 11<br>9<br>12 |
| **AN0995** | **unchar.** | Putative CLASP family microtubule-associated protein, putative ortholog of *S. pombe* Peg1 | 14.86<br>17.08<br>17.33 | 18.97<br>19.8<br>20.4 | 0<br>0<br>0 | 18<br>15<br>14 | 0<br>0<br>0 | 9<br>4<br>11 |

| | | | | | | | | |
|---|---|---|---|---|---|---|---|---|
| **AN3029** | **unchar.** | Putative AP-1 adaptor complex subunit beta, Putative ortholog of *C. albicans* APL2 | 15.7<br>17.65<br>17.42 | 20.13<br>20.67<br>21.54 | 1<br>0<br>0 | 38<br>36<br>30 | 1<br>0<br>0 | 10<br>9<br>16 |
| **AN0706** | **unchar.** | Ortholog(s) have role in ER to Golgi vesicle-mediated transport, putative ortholog of *C. albicans* USO1 | 15<br>17.79<br>16.9 | 20.22<br>20.38<br>22.36 | 0<br>0<br>1 | 61<br>17<br>42 | 0<br>0<br>1 | 19<br>6<br>24 |
| **AN4168** | **unchar.** | DUF500 and SH3 domain protein, putative ortholog of *S. pombe* actin cortical patch component Lsb4 | 16.7<br>17.45<br>17.45 | 19.95<br>15.67<br>20.16 | 0<br>0<br>0 | 37<br>0<br>14 | 0<br>0<br>0 | 7<br>0<br>7 |
| DNA binding | | | | | | | | |
| **AN0228** | **unchar.** | Putative ortholog of *S. cerevisiae* replication licensing factor MCM6 | 15.82<br>17.3<br>16.84 | 19.2<br>20.24<br>21.01 | 0<br>0<br>0 | 28<br>13<br>25 | 0<br>0<br>0 | 11<br>7<br>13 |
| **AN6070** | **unchar.** | Putative ortholog of *S. cerevisiae* replication licensing factor MCM4 | 14.52<br>17.26<br>16.69 | 18.67<br>19.74<br>20.55 | 0<br>0<br>0 | 17<br>13<br>14 | 0<br>0<br>0 | 8<br>6<br>10 |
| **AN2278** | **unchar.** | Putative ortholog of *N. crassa* SNF2-family ATP dependent chromatin remodeling factor Snf21 (Crf3-1) | 14.56<br>17.69<br>17.05 | 20.02<br>20.34<br>21.76 | 0<br>0<br>1 | 58<br>28<br>41 | 0<br>0<br>1 | 15<br>8<br>17 |
| **AN4187** | **unchar.** | Putative ortholog of *N. crassa* TBP | 15.31<br>17.42<br>16.07 | 19.45<br>18.96<br>21.27 | 2<br>0<br>0 | 42<br>13<br>33 | 1<br>0<br>0 | 13<br>5<br>22 |

| | | | | | | | | |
|---|---|---|---|---|---|---|---|---|
| | | associated factor (Cfr8-1) | | | | | | |
| **<u>AN7222</u>** | **unchar.** | NACHT domain containing protein | 15.09<br>16.34<br>15.09 | 22.49<br>21.55<br>23.74 | 0<br>0<br>0 | 181<br>47<br>84 | 0<br>0<br>0 | 42<br>16<br>35 |
| **<u>AN5168</u>** | **unchar.** | Putative NACHT and Ankyrin domain protein, putative ortholog of *S. pombe* Akr1 | 15.45<br>17.06<br>16.85 | 20.19<br>20.52<br>21.23 | 0<br>0<br>0 | 56<br>27<br>29 | 0<br>0<br>0 | 13<br>8<br>15 |
| Unknown function | | | | | | | | |
| **AN3005** | **unchar.** | Protein of unknown function | 15.17<br>16.81<br>17.11 | 19.2<br>20-9<br>20.87 | 0<br>0<br>0 | 27<br>26<br>36 | 0<br>0<br>0 | 11<br>8<br>16 |
| **<u>AN11181</u>** | **unchar.** | Protein of unknown function | 15.18<br>18.12<br>17.48 | 19.32<br>20.17<br>20.58 | 0<br>0<br>0 | 35<br>12<br>14 | 0<br>0<br>0 | 12<br>6<br>11 |
| **AN5423** | **unchar.** | Protein of unknown function | 13.8<br>17.05<br>18.02 | 21.07<br>16.94<br>20.47 | 0<br>0<br>0 | 68<br>0<br>6 | 0<br>0<br>0 | 13<br>0<br>4 |

Proteins were identified in at least two out of three biological repetitions with MS/MS counts ≥ 4, unique peptides ≥ 3 and LFQ intensity ≥ 20 and which were absent in the control strain. Proteins with underlined AN numbers contain a nuclear localization signal predicted with cNLS mapper *[2]* with a score ≤ 5 (indicating a localization in both, nucleus and cytoplasm), Sys. Name = systematic name, Std. name = standard name, unchar. = uncharacterized. Descriptions were obtained and adapted from FungiDB and NCBI [3,4].

**Table S4: Proteins significantly enriched with *V. dahliae* Vel2-GFP and Vel2$^{\Delta IDD}$-GFP, their predicted domains and potential *A. nidulans* counterparts.**

| ***V. dahliae* Identifier** | **Best Hit in *Aspergillus nidulans*** | **Domain(s)** |
|---|---|---|
| **Proteins co-purified with Vel2-GFP** | | |
| VDAG_JR2_Chr1g11550a | AN7710 | Haloacid dehalogenase-like hydrolase |
| VDAG_JR2_Chr1g13580a | AN7594 | Glutathione-dependent formaldehyde-activating enzyme/centromere protein V |
| VDAG_JR2_Chr1g17980a | AN6337 | Peptidase S8 |
| VDAG_JR2_Chr1g20590a | AN9408 (FasB) | Acyl transferase<br>Fatty acid synthase<br>MaoC-like dehydratase domain<br>Starter unit:ACP transacylase |
| VDAG_JR2_Chr1g20610a | AN7880 | Beta-ketoacyl synthase<br>Fatty acid synthase |
| VDAG_JR2_Chr1g23770a | AN2694 | Short-chain dehydrogenase/reductase SDR |
| VDAG_JR2_Chr2g08140a | AN6399 | Peptidase C1B, bleomycin hydrolase |
| VDAG_JR2_Chr3g01680a | AN8959 | TspO/MBR-related protein |
| VDAG_JR2_Chr3g03700a | AN10709 | Sugar isomerase (SIS) |
| **VDAG_JR2_Chr3g06150a** | **AN0363 (VelB)** | **Velvet domain** |
| VDAG_JR2_Chr3g07650a | AN4914 (Ack) | Aliphatic acid kinase, short-chain |
| VDAG_JR2_Chr3g10860a | AN2068 | K Homology domain |
| **VDAG_JR2_Chr3g12090a** | **AN1959 (VosA)** | **Velvet domain** |
| VDAG_JR2_Chr3g13120a | AN3712 | Dienelactone hydrolase |
| VDAG_JR2_Chr4g02180a | AN5989 | NAD(P)-binding domain |
| VDAG_JR2_Chr4g02640a | AN8979 (AlcA) | Alcohol dehydrogenase |
| VDAG_JR2_Chr4g06150a | AN2435 (AclA) | ATP-citrate synthase, citrate-binding domain |
| VDAG_JR2_Chr4g06160a | AN2436 (AclB) | Citrate synthase |
| VDAG_JR2_Chr4g07720a | AN0443 | Alcohol dehydrogenase |
| VDAG_JR2_Chr4g08010a | AN11985 | Class I glutamine amidotransferase-like |

| | | |
|---|---|---|
| VDAG_JR2_Chr4g08190a | AN6731 | Cytochrome b5-like heme/steroid binding domain<br>Fatty acid desaturase domain |
| VDAG_JR2_Chr4g09880a | AN5544 | HpcH/HpaI aldolase/citrate lyase domain |
| VDAG_JR2_Chr5g00110a | AN3091 | Peptidase M13 |
| VDAG_JR2_Chr5g03080a | AN7937 (CipC) | Protein of unknown function DUF3759 |
| VDAG_JR2_Chr5g05440a | AN2314 (Be1) | Glycosyl hydrolase, family 13 |
| VDAG_JR2_Chr5g09190a | AN3627 | Protein of unknown function DUF4449 |
| VDAG_JR2_Chr5g10680a | AN10301 | NmrA-like domain |
| VDAG_JR2_Chr6g01710a | AN7590 | Enoyl-(Acyl carrier protein) reductase |
| VDAG_JR2_Chr6g06890a | AN10518 | Perilipin protein |
| VDAG_JR2_Chr6g10140a | AN7388 (CpeA) | Haem peroxidase |
| VDAG_JR2_Chr7g00220a | AN3814 (CypA) | Cyclophilin-type peptidyl-prolyl cis-trans isomerase domain |
| VDAG_JR2_Chr7g00600a | AN0248 (PdiB) | Thioredoxin domain |
| **VDAG_JR2_Chr7g04890a** | **AN1052 (VeA)** | **Velvet domain** |
| VDAG_JR2_Chr7g05280a | AN9103 (AifA) | Rieske [2Fe-2S] iron-sulphur domain<br>FAD/NAD(P)-binding domain |
| VDAG_JR2_Chr7g08750a | AN2797 | - |
| VDAG_JR2_Chr8g02960a | AN6048 | Aminotransferase, class I/classII |
| VDAG_JR2_Chr8g04960a | AN9148 (GalF) | UDPGP family |
| VDAG_JR2_Chr8g08780a | AN2867 (PgmB) | Alpha-D-phosphohexomutase |
| VDAG_JR2_Chr8g10760a | AN3414 | NADP-dependent oxidoreductase domain |
| **Proteins co-purified with Vel2-GFP deta insert** | | |
| VDAG_JR2_Chr2g07820a | AN6653 (AcuE) | Malate synthase |
| VDAG_JR2_Chr3g00330a | AN4463 | Clathrin, heavy chain |
| **VDAG_JR2_Chr3g06150a** | **AN0363 (VelB)** | **Velvet domain** |
| **VDAG_JR2_Chr7g04890a** | **AN1052 (VeA)** | **Velvet doamin** |

For identification of proteins and their conserved domains, EnsemblFungi was used [5].

**Table S5: VelB-VosA heterodimer formation is independent of the IDD in a *veA* deletion background.**

| | **Sys. name** | **AN0363** | **AN1052** | **AN1959** | **AN0807** |
|---|---|---|---|---|---|
| | **Std name** | **VelB** | **VeA** | **VosA** | **LaeA** |
| | **bait** | | | | |
| **LFQ intensity** | **VelB-GFP** | 29.33<br>28.63<br>28.43 | 27.27<br>26.97<br>27.1 | 27.08<br>22.69<br>23.94 | 25.4<br>20.97<br>23.28 |
| | **VelB-GFP in Δ*veA* strain** | 28.34<br>29.63<br>28.34 | 17.37<br>17.66<br>19.24 | 25.89<br>26.49<br>19.59 | 16.09<br>17.6<br>20.54 |
| | **VelB$^{\Delta IDD}$-GFP** | 28.29<br>29.17<br>29.3 | 24.39<br>26.25<br>27.36 | 18.41<br>18.58<br>19.82 | 24.39<br>22.14<br>22.66 |
| | **VelB$^{\Delta IDD}$-GFP in Δ*veA* strain** | 28.67<br>31.07<br>25.5 | 19.39<br>18.6<br>19.48 | 24.06<br>23.54<br>18.77 | 19.5<br>17.83<br>18.47 |
| **MS/MS counts** | **VelB-GFP** | 121<br>48<br>27 | 53<br>14<br>13 | 21<br>6<br>3 | 8<br>1<br>3 |
| | **VelB-GFP in Δ*veA* strain** | 101<br>70<br>20 | 0<br>0<br>0 | 17<br>10<br>3 | 0<br>0<br>0 |
| | **VelB$^{\Delta IDD}$-GFP** | 65<br>55<br>37 | 14<br>17<br>9 | 1<br>0<br>0 | 3<br>2<br>6 |
| | **VelB$^{\Delta IDD}$-GFP in Δ*veA* strain** | 106<br>131<br>7 | 0<br>0<br>0 | 8<br>3<br>0 | 0<br>0<br>0 |
| **Unique peptides** | **VelB-GFP** | 17<br>11<br>6 | 17<br>7<br>7 | 8<br>4<br>3 | 4<br>2<br>3 |
| | **VelB-GFP in Δ*veA* strain** | 12<br>11<br>9 | 0<br>0<br>0 | 7<br>6<br>3 | 0<br>0<br>0 |
| | **VelB$^{\Delta IDD}$-GFP** | 14<br>13<br>9 | 8<br>10<br>7 | 1<br>0<br>0 | 3<br>2<br>4 |
| | **VelB$^{\Delta IDD}$-GFP in Δ*veA* strain** | 14<br>15<br>4 | 0<br>0<br>0 | 4<br>3<br>0 | 0<br>0<br>0 |

Identification of VeA, VosA and LaeA as interaction partner of VelB with or without IDD in wild type or *veA* deletion (Δ*veA*) background. Thresholds were set as followed: MS/MS counts ≥ 3, LFQ intensity ≥ 20, unique peptides ≥ 2 and identified proteins are absent in the GFP OE control strain in both biological replicates. Std. name = systematic name, Sys. name = systematic name.

**Table S6: Secondary metabolites identified in this study.**

| Peak | Chemical name | Molecular Formula | Calc. exact mass [M] | Measured exact mass | Rt (min) | Confirmed by | Ref. | Gene cluster, main enzyme |
|---|---|---|---|---|---|---|---|---|
| 1 | Cichorine | $C_{10}H_{11}NO_3$ | 193.07389 | 194.0813 $[M+H]^+$ | 7.7 | A, C | [6] | *cic*, CicF (nrPKS, AN6448) |
| 2 | F9775A/B | $C_{21}H_{16}O_8$ | 396.08452 | 397.0923 $[M+H]^+$ | 9.6, 10.6 | A, C | [7] | *ors*, orsA (nrPKS, AN7909) |
| 3 | Austinol | $C_{25}H_{40}O_8$ | 458.19407 | 459.2014 $[M+H]^+$ | 12.7 | A, B | [8] | *aus*, AusA (nrPKS, AN8383) |
| 4 | Dehydroaustinol | $C_{25}H_{28}O_8$ | 456.17842 | 457.1858 $[M+H]^+$ | 13.1 | A, B | [8] | *aus*, AusA (nrPKS, AN8383) |
| 5 | Arugosin H | $C_{20}H_{20}O_6$ | 356.12599 | 356.1476 $[M+H]^+$ | 14.3 | C | [9] | *mdp*, mdpG (nrPKS, AN0150) |
| 6 | Emericellamide C | $C_{30}H_{53}N_5O_7$ | 595.39450 | 596.40233 $[M+H]^+$ | 15.1 | A | [10] | *eas*, EasA/B (NRPS-PKS, AN2545/ AN2547) |
| 7 | Sterigmatocystin | $C_{18}H_{12}O_6$ | 324.06339 | 325.0706 $[M+H]^+$ | 15.3 | A, B | [11] | *stc*, StcA (nrPKS, AN7825) |
| 8 | Emericellamide E | $C_{32}H_{57}N_5O_7$ | 623.42580 | 624.4323 $[M+H]^+$ | 17.3 | A, D | [10] | *eas*, EasA/B (NRPS-PKS, AN2545/ AN2547) |
| 9 | Terrequinone A | $C_{32}H_{30}O_3N_2$ | 490.22564 | 491.2320 $[M+H]^+$ | 18.5 | A, D | [12] | *tdi*, TdiA (NRPS, AN8513) |
| 10 | Arugosin A | $C_{25}H_{28}O_6$ | 424.18859 | 425.19642 $[M+H]^+$ | 17-21 | A, C | [9] | *mdp*, mdpG (NR-PKS, AN0150) |
| 11 | Emericellin | $C_{25}H_{28}O_5$ | 408.19368 | 409.2002 $[M+H]^+$ | 24.3 | A, B | [13] | mdp, mdpG (NR-PKS, AN0150) |

| | | | | | | | | |
|---|---|---|---|---|---|---|---|---|
| 12 | Shamixanthone | $C_{25}H_{26}O_5$ | 406.17803 | 389.1740 [M-$H_2O$+H]$^+$ | 24.5 | A, B | [13] | mdp, mdpG (NR-PKS, AN0150) |
| 13 | Epishamixanthone | $C_{25}H_{26}O_5$ | 406.17803 | 389.1740 [M-$H_2O$+H]$^+$ | 25.4 | A, B | [13] | mdp, mdpG (NR-PKS, AN0150) |

Table corresponds to Fig 5, Fig S11, Fig S12 and Fig S13. A: Exact mass measurement (for Emericellamide C: [14,15]), B: retention time [16,17]/comparison with commercial standards for sterigmatocystin, C: UV/VIS spectrum [7,18,19], D: MS/MS fragmentation [20,21]. Ref. = reference, PKS = polyketide synthase, NR-PKS = non-reducing polyketide synthase, NRPS = non ribosomal polyketide synthase, *cic* = chicorine, ors = orsellinic acid, *aus* = austinol, *mdp* = monodictophenone, *eas* = emericellamide, *stc* = sterigmatocystein, *tdi* = (tryptophane-derived) terrequinone gene clusters.

**Table S7: Fungal strains used in this study.**

| Strain | Genotype | Reference |
|---|---|---|
| | ***A. nidulans*** **strains** | |
| **FGSC A4** | *veA*$^{+}$ | [22] |
| **AGB551** | Δ*nkuA*::*argB*, *pyrG89*, *pyroA4, veA*$^{+}$ | [23] |
| **AGB596** | $^{P}$*gpdA*::*gfp-phleo*$^{R}$; *pabaA1*, *yA2*, *veA*$^{+}$ | [23] |
| **AGB1057** | Δ*nkuA*::*argB*, *pyroA4*, *pyrG89*, *veA*$^{+}$, Δ*vosA*:*six* | [17] |
| **AGB1064** | Δ*nkuA*::*argB*, *pyroA4*, *pyrG89*, *veA*$^{+}$, Δ*velB*:*six* | [17] |
| **AGB1066** | Δ*nkuA*::*argB*, *pyroA4*, *pyrG89*, *veA*$^{+}$, Δ*veA*:*six* | [17] |
| **AGB1131** | Δ*nkuA::argB, pyroA4, pyrG89, veA*$^{+}$*, velB*$^{133\text{-}231\Delta:}$*six* | This study |
| **AGB1132** | Δ*nkuA*::*argB*, *pyroA4*, *pyrG89*, *veA*$^{+}$, *velB*:*gfp*:*six* | This study |
| **AGB1133** | *nkuA*Δ::*argB*, *pyroA4*, *pyrG89*, *veA*$^{+}$, *velB*$^{\Delta IDD}$:*gfp*:*six* | This study |
| **AGB1134** | Δ*nkuA::argB, pyroA4, pyrG89, veA+, velB*$^{\Delta 133\text{-}231}$*:velB*$^{Af131\text{-}322}$*:six* | This study |
| **AGB1135** | Δ*nkuA::argB, pyroA4, pyrG89, veA+, velB*$^{\Delta 133\text{-}231}$*:velB*$^{Vd210\text{-}327}$*:six* | This study |
| **AGB1136** | Δ*nkuA*::*argB*, *pyroA4*, *pyrG89*, *veA*$^{+}$, *velB*:*gfp*:*six, vosA:ha*:*six* | This study |
| **AGB1137** | Δ*nkuA*::*argB*, *pyroA4*, *pyrG89*, *veA*$^{+}$, *velB*$^{\Delta 133\text{-}231}$*:gfp*:*six, vosA:ha*:*six* | This study |
| **AGB1140** | Δ*nkuA*::*argB*, *pyroA4*, *pyrG89*, *veA*$^{+}$, Δ*veA*:*six, velB*$^{\Delta 133\text{-}231:}$*six* | This study |
| **AGB1142** | Δ*nkuA*::*argB*, *pyroA4*, *pyrG89*, *veA*$^{+}$, Δ*vosA*:*six, velB*$^{\Delta 133\text{-}231:}$*six* | This study |
| **AGB1149** | Δ*nkuA::argB, pyroA4, pyrG89, veA+, velB:gfp:six, veA:ha* | This study |
| **AGB1150** | Δ*nkuA::argB, pyroA4, pyrG89, veA+, velB*$^{\Delta 133\text{-}231}$*:gfp:six, veA:ha* | This study |
| **AGB1190** | *nkuA*Δ*::argB, pyroA4, pyrG89, veA+, velB:gfp:six,* Δ*veA* | This study |
| **AGB1191** | Δ*nkuA::argB, pyroA4, pyrG89, veA+, velB(*Δ*133-231):gfp:six,* Δ*veA* | This study |
| | ***V. dahliae*** **strains** | |
| **JR2** | Wild type isolate from *Solanum lycopersicum* | [24] |
| **VGB58** | Δ*vel2:*$^{P}$*gpdA:nat*$^{R}$ | [25] |
| **VGB450** | $^{P}$*vel2:vel2:gfp:*$^{P}$*gpdA:hyg*$^{R}$*:trpC*$^{T}$*:vel2*$^{T}$ | [25] |
| **VGB468** | $^{P}$*vel2:vel2*$^{\Delta 210\text{-}325}$*:gfp:*$^{P}$*gpdA:hyg*$^{R}$*:trpC*$^{T}$*:vel2*$^{T}$ | This study |

Strains were constructed by employment of recyclable marker cassettes, which only leave a small six site (100 nucleotides) as scar after recycling of the marker off the genome. $^{P}$=promoter, $^{T}$=terminator, NAT$^{R}$=nourseothricin resistance marker, HYG$^{R}$=hygromycin B resistance marker; FGSC= Fungal genetics stock center, Kansas, USA.

**Table S8: Plasmids constructed and used in this study**.

| **Plasmid** | **Description** | **Reference** |
|---|---|---|
| pBluescript SK(+) | Cloning vector, *amp*$^{R}$ | Thermo Scientific |
| pETM13 | Expression vector with C-term. His-tag | EMBL (Heidelberg) |
| pME3815 | *velB:his* from cDNA, cloned in pETM-13 | [26] |
| pME4292 | Plasmid contains *gfp* | [27] |
| pME4304 | *Six:*P*xyl*$^{P}$:β-*rec*:*trpC*$^{t:}$*nat*$^{R:}$*six* with *Sfi*I restriction sites | [17] |
| pME4319 | *six*-P*xyl*$^{P}$:β-*rec*:*trpC*$^{t:}$*phleo*$^{R:}$*six* in *Eco*RV of pBluescript SK+ with *Eco72*I and *Swa*I restriction sites | [28] |
| pME4564 | $^{P}$*TRPC:HYG*$^{R}$*; KAN*$^{R}$ Cloning for *Agrobacterium*-mediated transformation of *V. dahliae* | [29] |
| pME4574 | Δ*veA*::natRM | [17] |
| pME4305 | *six*-P*xyl*$^{P}$:β-*rec*:*trpC*$^{t}$:*phleo*$^{R}$:*six* with *Sfi*I restriction sites | [17] |
| pME4604 | Δ*velB*::natRM | [17] |
| pME4686 | *velB*$^{\Delta IDD}$ = *velB*$^{\Delta 133\text{-}231}$:phleoRM | This study |
| pME4687 | *velB:gfp:*natRM | This study |
| pME4688 | *velB*$^{\Delta IDD}$*:gfp* = *velB*$^{\Delta 133\text{-}231}$*:gfp:*phleoRM | This study |
| pME4689 | 3' flanking region of *velB* in *Eco72*I of pME4319 | This study |
| pME4690 | *velB*$^{AfIDD}$ = *velB*$^{Af131\text{-}322}$:phleoRM | This study |
| pME4691 | *velB*$^{VdIDD}$ *velB*$^{Vd210\text{-}327}$:phleoRM | This study |
| pME4692 | *velB*$^{\Delta 133\text{-}231}$*:his* amplified from pME3815, cloned in pETM-13 | This study |
| pME4693 | *vosA-ha::*phleoRM | This study |
| pME4748 | *veA-ha::*natRM | This study |
| pME4990 | $^{P}$*GPDA:NLP3:GFP:TRPC*$^{T}$*:*$^{P}$*GPDA:HYG*$^{R}$*:TRPC*$^{T}$ in pPK2 | [29] |
| pME5075 | $^{P}$*VEL2:VEL2*$^{\Delta IDD}$*:*$^{P}$*GPDA:HYG*$^{R}$*:TRPC*$^{T}$ in pPK2 | This study |
| pME5077 | $^{P}$*VEL2:VEL2*$^{\Delta IDD}$*:GFP:*$^{P}$*GPDA:HYG*$^{R}$*:TRPC*$^{T}$*:VEL2*$^{T}$ in pME4564 | This study |
| pME5076 | $^{P}$*VEL2:VEL2*$^{\Delta IDD}$*:*$^{P}$*GPDA:HYG*$^{R}$*:TRPC*$^{T}$*:VEL2*$^{T}$ in pME5075 | This study |
| pNI47 | *vosA:gst* in pGEX-5X-1 (Amersham) | [30] |
| pPK2 | $^{P}$*GPDA:HYG*$^{R}$*:TRPC*$^{T}$*; KAN*$^{R}$ Cloning for *Agrobacterium*-mediated transformation of *V. dahliae* | [31] |

pBluescript SK+ was used as backbone for all plasmids constructed in this study, if not stated otherwise. $^{P}$ = promoter, $^{t}$ = terminator, $^{R}$ = resistance, natRM = recyclable nat$^{R}$ resistance cassette from pME4304, phleoRM = recyclable phleo$^{R}$ resistance cassette from pME4305, p.c. = personal communication; hyg$^{R}$ = hygromycine resistance gene; Kan$^{R}$ = kanamycine resistance.

**Table S9: Oligonucleotides used for amplifications and plasmid constructions.**

| Name | 5' – sequence – 3' |
|---|---|
| AO74 | TGC TTG CCA TCT TGC TAC ACC |
| AO75 | GTA CAT CCC CTG AGG GTG GCG GAC GAG GTT CAC CT |
| AO76 | CCT CAG GGG ATG TAC ACG |
| AO77 | CTA ATA ATC GTC ATC GTC GT |
| AO78 | TTT TCT AGA ATG AGG CGC TGG TCA TTC AA |
| AO79 | TAG TCT AGA TGC ATC TTC AGA CAC GCA AA |
| AO165 | GGT GGT AGC GGT GGT GT |
| AO167 | ATT CTT AAT TAA GAT TGC TTG CCA TCT TGC TAC ACC |
| AO168 | ACC ACC GCT ACC ACC ATA ATC GTC ATC GTC GTC A |
| AO169 | AGG TAA TCC TTC TTT CTA GAA TGA GGC GCT GGT |
| AO170 | AGG ACT TCT AGA AGG TGC ATC TTC AGA CAC GCA |
| JG45 | CCC CAT GGC GAT GTA CGC TGT TGA GGA TAG G |
| JG46 | CCC TCG AGG TAT TCG TTA TCC AGA CCA TC |
| KT142 | ATA ATA TGG CCA TCT AAG AAT TCT GCC GGC GTT TAT TTG |
| KT163 | CTA TAG GCC TGA GTG TTA GGC GTA GTC GGG GAC G |
| KT166 | CAA AGC CTG CCA CCA TGC GTT ACC CCT ACG ACG TCC CCG ACT ACG CCT AA |
| KT197 | ATC GAT AAG CTT GAT GTT TAA ACT GGA GTG CCT TTC GTC |
| KT198 | CTG CAG GAA TTC GAT GTT TAA ACA TTC TGG CTC GTC TGC |
| RH514 | ACC GGT CAC TGT ACA TTA CTT GTA CAG CTC GTC CAT |
| RH590 | TGT ACA GTG ACC GGT GA |
| RO4 | AAA GAA GGA TTA CCT CTA AAC AA |
| SR05 | CTG CAG GAA TTC GAT GTT TAA ACC GTG CAG TCA GTC TAC CTA C |
| SR07 | ATA ATA TGG CCA TCT AGA CCG TAT ATT GTT TCA TAA ATC C |
| SR08 | ATC GAT AAG CTT GAT GTT TAA ACC CGC TGT ACA TGT AAT GTC CG |
| SR18 | GGT GGT AGC GGT GGT GTG AGC AAG GGC GAG GAG |
| SR20 | CTA TAG GCC TGA GTG CTA CTT GTA CAGT TCG TCC ATG C |
| SR24 | ACC ACC GCT ACC ACC GTA TTC GTT ATC CAG ACC ATC G |
| SR49 | ATA ATA TGG CCA TCT GGA TTC TCG TTT GTG GAA CAC |

| | |
|---|---|
| SR75 | ATC GAT AAG CTT GAT GTT TAA ACT TTC CGT AGG TCG ATC C |
| SR76 | AGG AAT TCG ATG TTT AAA CAA GGG CTC CTG TCG GA |
| SR108 | GGC ATG TTC ACG CGC AAT CT |
| SR109 | AGA TTG CGC GTG AAC ATG CCG TGC TTC ACA AGA TTT ACT TCA TG |
| SR110 | AGG AAT TCG ATG TTT AAA CGA TTA GGA GAA GTC CAC TTT |
| SR111 | TTG ACC TAT AGG CCT TTA GTA TTC GTT ATC CAG ACC ATC |
| SR112 | ATA ATA TGG CCA TCT AGA CCG TAT ATT GTT TCA TAA ATC C |
| SR113 | ATA AGC TTG ATG TTT AAA CCG CTC GCG CCC CAT |
| SR201 | CTA TAG GCC TGA GTG TTA GGC GTA GTC GGG GAC GTC GTA GGG GTA CCG AGG AGT TCC GTT CGC |
| SR253 | GTG CTT CAC AAG ATT TAC TTC ATG |
| SR254 | AAG TAA ATC TTG TGA AGC ACC AGC GCG AGG TGA ACC TC |
| SR255 | AGA TTG CGC GTG AAC ATG CCC TGA GGC GCA CCT GCG |
| SR266 | AAG TAA ATC TTG TGA AGC ACT CAG CCA CGT CGC CAT CAA TAT C |
| SR267 | AGA TTG CGC GTG AAC ATG CCA CCG GGC CCC GCC GGA A |

**Table S10: Primers used for qRT-PCR in this study.**

| Name | Gene | 5' – sequence – 3' |
|---|---|---|
| jg816 | *velB* A | CCC CTC CGT GTA TCC GTC TAA T |
| jg817 | *velB* B | AGC CGA GTG CTT CAC AAG ATT T |
| kt278 | *15S rRNA* A | GAT CCG CGA AAA ACC TTA CCA C |
| kt279 | *15S rRNA* B | TGG CAC GTC TAT AGC CCA CAG T |
| kt312 | *h2A.X* A | TCT CGA GCT TGC TGG AAA CG |
| kt313 | *h2A.X* B | CAC CCT GGG CAA TAG TGA CG |

### References

1. Ward JJ, McGuffin LJ, Bryson K, Buxton BF, Jones DT. The DISOPRED server for the prediction of protein disorder. Bioinformatics. 2004;20: 2138–2139. doi:10.1093/bioinformatics/bth195
2. Kosugi S, Hasebe M, Tomita M, Yanagawa H. Systematic identification of cell cycle-dependent yeast nucleocytoplasmic shuttling proteins by prediction of composite motifs. Proceedings of the National Academy of Sciences. 2009;106: 10171–10176. doi:10.1073/pnas.0900604106
3. Alvarez-Jarreta J, Amos B, Aurrecoechea C, Bah S, Barba M, Barreto A, et al. VEuPathDB: the eukaryotic pathogen, vector and host bioinformatics resource center in 2023. Nucleic Acids Res. 2024;52: D808–D816. doi:10.1093/NAR/GKAD1003
4. Sayers EW, Bolton EE, Brister JR, Canese K, Chan J, Comeau DC, et al. Database resources of the national center for biotechnology information. Nucleic Acids Res. 2022;50: D20–D26. doi:10.1093/NAR/GKAB1112
5. Harrison PW, Amode MR, Austine-Orimoloye O, Azov AG, Barba M, Barnes I, et al. Ensembl 2024. Nucleic Acids Res. 2024;52: D891–D899. doi:10.1093/nar/gkad1049
6. Sanchez JF, Entwistle R, Corcoran D, Oakley BR, Wang CCC. Identification and molecular genetic analysis of the cichorine gene cluster in *Aspergillus nidulans*. Medchemcomm. 2012;3: 997–1002. doi:10.1039/C2MD20055D
7. Bok JW, Chiang Y-M, Szewczyk E, Reyes-Dominguez Y, Davidson AD, Sanchez JF, et al. Chromatin-level regulation of biosynthetic gene clusters. Nat Chem Biol. 2009;5: 462–464. doi:10.1038/nchembio.177
8. Lo HC, Entwistle R, Guo CJ, Ahuja M, Szewczyk E, Hung JH, et al. Two separate gene clusters encode the biosynthetic pathway for the meroterpenoids austinol and dehydroaustinol in *Aspergillus nidulans*. J Am Chem Soc. 2012;134: 4709–4720. doi:10.1021/ja209809t
9. Nielsen ML, Nielsen JB, Rank C, Klejnstrup ML, Holm DK, Brogaard KH, et al. A genome-wide polyketide synthase deletion library uncovers novel genetic links to polyketides and meroterpenoids in *Aspergillus nidulans*. FEMS Microbiol Lett. 2011;321: 157–166. doi:https://doi.org/10.1111/j.1574-6968.2011.02327.x
10. Chiang Y-M, Szewczyk E, Nayak T, Davidson AD, Sanchez JF, Lo H-C, et al. Molecular Genetic Mining of the Aspergillus Secondary Metabolome: Discovery of the Emericellamide Biosynthetic Pathway. Chem Biol. 2008;15: 527–532. doi:https://doi.org/10.1016/j.chembiol.2008.05.010
11. Yu JH, Leonard TJ. Sterigmatocystin biosynthesis in *Aspergillus nidulans* requires a novel type I polyketide synthase. J Bacteriol. 1995;177: 4792–4800. doi:10.1128/JB.177.16.4792-4800.1995
12. Bouhired S, Weber M, Kempf-Sontag A, Keller NP, Hoffmeister D. Accurate prediction of the *Aspergillus nidulans* terrequinone gene cluster boundaries using the transcriptional regulator LaeA. Fungal Genetics and Biology. 2007;44: 1134–1145. doi:https://doi.org/10.1016/j.fgb.2006.12.010
13. Sanchez JF, Entwistle R, Hung J-H, Yaegashi J, Jain S, Chiang Y-M, et al. Genome-based deletion analysis reveals the prenyl xanthone biosynthesis pathway in *Aspergillus nidulans*. J Am Chem Soc. 2011;133: 4010–7. doi:10.1021/ja1096682
14. Ahmed AM, Ibrahim AM, Yahia R, Shady NH, Mahmoud BK, Abdelmohsen UR, et al. Evaluation of the anti-infective potential of the seed endophytic fungi of Corchorus olitorius through metabolomics and molecular docking approach. BMC Microbiol. 2023;23: 1–19. doi:10.1186/S12866-023-03092-5/FIGURES/11
15. Perlatti B, Lan N, Jiang Y, An Z, Bills G. Identification of Secondary Metabolites from Aspergillus pachycristatus by Untargeted UPLC-ESI-HRMS/MS and Genome Mining. Molecules. 2020;25. doi:10.3390/MOLECULES25040913
16. Liu L, Sasse C, Dirnberger B, Valerius O, Fekete-Szücs E, Harting R, et al. Secondary metabolites of hülle cells mediate protection of fungal reproductive and overwintering structures against fungivorous animals. Elife. 2021;10. doi:10.7554/ELIFE.68058

17. Thieme KG, Gerke J, Sasse C, Valerius O, Thieme S, Karimi R, et al. Velvet domain protein VosA represses the zinc cluster transcription factor SclB regulatory network for *Aspergillus nidulans* asexual development, oxidative stress response and secondary metabolism. Copenhaver GP, editor. PLoS Genet. 2018;14: e1007511. doi:10.1371/journal.pgen.1007511
18. Kralj A, Kehraus S, Krick A, Eguereva E, Kelter G, Maurer M, et al. Arugosins G and H: prenylated polyketides from the marine-derived fungus Emericellanidulans var. acristata. J Nat Prod. 2006;69: 995–1000. doi:10.1021/NP050454F
19. Nielsen KF, Månsson M, Rank C, Frisvad JC, Larsen TO. Dereplication of microbial natural products by LC-DAD-TOFMS. J Nat Prod. 2011;74: 2338–2348. doi:10.1021/NP200254T/SUPPL_FILE/NP200254T_SI_001.ZIP
20. Hamed AA, El-Shiekh RA, Mohamed OG, Aboutabl EA, Fathy FI, Fawzy GA, et al. Cholinesterase Inhibitors from an Endophytic Fungus Aspergillus niveus Fv-er401: Metabolomics, Isolation and Molecular Docking. Molecules. 2023;28: 2559. doi:10.3390/MOLECULES28062559/S1
21. Chiang YM, Szewczyk E, Nayak T, Davidson AD, Sanchez JF, Lo HC, et al. Molecular genetic mining of the Aspergillus secondary metabolome: discovery of the emericellamide biosynthetic pathway. Chem Biol. 2008;15: 527–532. doi:10.1016/J.CHEMBIOL.2008.05.010
22. McCluskey K, Wiest A, Plamann M. The Fungal Genetics Stock Center: a repository for 50 years of fungal genetics research. J Biosci. 2010;35: 119–26.
23. Bayram Ö, Bayram ÖS, Ahmed YL, Maruyama J, Valerius O, Rizzoli SO, et al. The *Aspergillus nidulans* MAPK module AnSte11-Ste50-Ste7-Fus3 controls development and secondary metabolism. Madhani HD, editor. PLoS Genet. 2012;8: e1002816. doi:10.1371/journal.pgen.1002816
24. Fradin EF, Zhang Z, Juarez Ayala JC, Castroverde CDM, Nazar RN, Robb J, et al. Genetic dissection of Verticillium wilt resistance mediated by tomato Ve1. Plant Physiol. 2009;150: 320–32. doi:10.1104/pp.109.136762
25. Höfer AM, Harting R, Aßmann NF, Gerke J, Schmitt K, Starke J, et al. The velvet protein Vel1 controls initial plant root colonization and conidia formation for xylem distribution in Verticillium wilt. PLoS Genet. 2021;17. doi:10.1371/JOURNAL.PGEN.1009434
26. Ahmed YL, Gerke J, Park H-S, Bayram Ö, Neumann P, Ni M, et al. The Velvet family of fungal regulators contains a DNA-binding domain structurally similar to NF-κB. Stock AM, editor. PLoS Biol. 2013;11: e1001750. doi:10.1371/journal.pbio.1001750
27. Jöhnk B, Bayram Ö, Abelmann A, Heinekamp T, Mattern DJ, Brakhage AA, et al. SCF ubiquitin ligase F-box protein Fbx15 controls nuclear co-repressor localization, stress response and virulence of the human pathogen *Aspergillus fumigatus*. Lin X, editor. PLoS Pathog. 2016;12: e1005899. doi:10.1371/journal.ppat.1005899
28. Gerke J, Köhler AM, Wennrich J-P, Große V, Shao L, Heinrich AK, et al. Biosynthesis of Antibacterial Iron-Chelating Tropolones in *Aspergillus nidulans* as Response to Glycopeptide-Producing Streptomycetes. Frontiers in Fungal Biology. 2022;2. doi:10.3389/ffunb.2021.777474
29. Leonard M, Kühn A, Harting R, Maurus I, Nagel A, Starke J, et al. *V. longisporum* elicits media-dependent secretome responses with a further capacity to distinguish between plant-related environments. bioRxiv. 2020; 2020.02.11.943803. doi:10.1101/2020.02.11.943803
30. Park H-S, Nam T-Y, Han K-H, Kim SC, Yu J-H. VelC Positively Controls Sexual Development in Aspergillus nidulans. PLoS One. 2014;9: e89883. doi:10.1371/journal.pone.0089883
31. Covert SF, Kapoor P, Lee M, Briley A, Nairn CJ. *Agrobacterium tumefaciens*-mediated transformation of *Fusarium circinatum*. Mycol Res. 2001;105: 259–264. doi:10.1017/S0953756201003872
